# Replicative aging impedes stress-induced assembly of a key human protein disaggregase

**DOI:** 10.1101/2022.06.25.497591

**Authors:** Yasith Mathangasinghe, Niels Alberts, Carlos J. Rosado, Dezerae Cox, Natalie L. Payne, Angelique R. Ormsby, Keziban Merve Alp, Roman Sakson, Sanjeev Uthishtran, Thomas Ruppert, Senthil Arumugam, Danny M. Hatters, Harm H. Kampinga, Nadinath B. Nillegoda

**Affiliations:** Australian Regenerative Medicine Institute, Monash University, Clayton, Victoria, Australia; Department of Biomedical Sciences of Cells & Systems, University Medical Center Groningen (UMCG) and University of Groningen (RuG), Groningen, The Netherlands; Department of Diabetes, Central Clinical School, Monash University, Melbourne, Victoria, Australia; Department of Biochemistry and Pharmacology, Bio21 Molecular Science and Biotechnology Institute, The University of Melbourne, Melbourne, Victoria, Australia; Max Delbrück Center for Molecular Medicine in the Helmholtz Association, Berlin, Germany; Leibniz-Institut für Analytische Wissenschaften-ISAS-e.V., Dortmund, Germany; Monash Biomedicine Discovery Institute, Monash University, Melbourne, Victoria, Australia; Core facility for Mass Spectrometry and Proteomics, Zentrum für Molekulare Biologie der Universität Heidelberg, Heidelberg, Germany; European Molecular Biological Laboratory Australia (EMBL Australia), Monash University, Melbourne, Victoria, Australia; Australian Research Council Centre of Excellence for Advanced Molecular Imaging, Monash University, Clayton, Victoria, Australia; Centre for Dementia and Brain Repair at the Australian Regenerative Medicine Institute, Monash University, Melbourne, Victoria, Australia

**Keywords:** Protein disaggregation, cellular aging, senescence, heat shock, Hsp70, J-domain proteins, VCP, amyloid

## Abstract

The collapse of protein homeostasis manifests itself in a toxic protein aggregation cascade, which is associated with degenerative diseases and aging. To solubilize aggregates, dedicated protein disaggregases exist in unicellular organisms, but these have no nuclear/cytosolic orthologs in metazoa. Alternative metazoan disaggregation machines have been described, but how these are operated and regulated *in vivo* remained unknown. We show that protein disaggregases are functionally diversified in human cells to efficiently target different types of stress-induced aggregates in sequential and temporally distinct phases. In particular, we show the selective assembly of an Hsp70-DNAJA1-DNAJB1 trimeric disaggregase that forms during late phase of stress recovery., *i.e.*, after VCP-dependent solubilization of non-native proteins that accumulate in cellular condensates such as nucleoli or stress granules. When activated, the trimeric disaggregase provides resistance to stress toxicity and contributes to amyloid disposal. Strikingly, this disaggregase collapses early in cells undergoing replicative aging with important underlining pathophysiological consequences.

## Main

The collapse of protein homeostasis and the increase in protein aggregation are closely associated with aging and many age-related diseases.^1,2^ Healthy cells maintain protein homeostasis by an elaborate protein quality control (PQC) network that supports the proper flux of protein biogenesis (protein synthesis, folding, transport) to protein degradation. Thereby, protein function is ensured, and the accumulation of potentially toxic misfolded or aggregated proteins is prevented. However, these PQC pathways can be overwhelmed by acute or chronic proteotoxic stresses, resulting in the accumulation of toxic protein aggregates that could trigger cell death. Cells have evolved survival mechanisms to reverse such deleterious effects of protein aggregation through protein disaggregation function.^3,4^ Previous studies show that the disruption of protein disaggregation function significantly decreases cell fitness and organismal lifespan.^5–8^

The machinery that performs aggregate solubilization is well understood in unicellular organisms (e.g., bacteria and fungi),^9^ where AAA+ ATPases of the Hsp100 class of chaperones (e.g., bacterial ClpB; yeast Hsp104) use a threading mechanism to untangle and release aggregate-trapped polypeptides.^9–11^ Metazoans lack cytosolic/nuclear Hsp100 disaggregases and instead appear to have evolved aggregate solubilization machines based on Hsp70.^12^ Evidence for metazoan protein disaggregation is provided by proof-of-principle biochemical studies showing the disassembly of artificially formed protein aggregates by reconstituted disaggregases with different Hsp70s and J-domain proteins (JDP).^12–15^ *In vivo* evidence is, however, limited to observational data showing the disappearance of aggregated proteins,^7,16^ reactivation of stress-insolubilized model proteins,^17,18^ and large-scale proteomics studies that monitored proteome solubility dynamics after proteotoxic stress insults.^19,20^ Although the induction of molecular chaperones including various Hsp70s and JDPs and the state of thermotolerance correlate with stress-induced aggregate disappearance,^7,21^ direct visualization of the assembly of functional Hsp70-based disaggregase complexes in metazoan cells has not been achieved. This aspect is challenging due to the unavailability of approaches that disentangle the effects of Hsp70 machineries that prevent protein aggregation (e.g., holdase and foldase) from possible disaggregase function.

Here, we used a protein-protein interaction-based method to track the complete assembly of a major disaggregase consisting of a trimeric Hsp70-DNAJA1-DNAJB1 complex on endogenous protein aggregates, which, for the first time, allowed us to dissect how aggregate solubilization is activated and regulated during repair of human cells after proteotoxic stress in space and time. Strikingly, the assembly of this trimeric disaggregase occurs in the late phase of stress recovery, sequential to and dependent on the resolubilization of stress-unfolded proteins from biomolecular condensates (nucleoli and stress granules). Resolubilization of condensates during the early phase of stress recovery is dependent on the action of another mammalian disaggregase, the AAA+ ATPase Valosin-Containing Protein (VCP).^22–24^ The two distinct disaggregases act on substrates with different solubility properties. Importantly, the late-acting Hsp70-DNAJA1-DNAJB1 trimeric disaggregase is crucial to stress tolerance and enhances the capacity of the cell to dispose of amyloid-type aggregates. Importantly, we show that activation of this Hsp70 disaggregase is severely hampered in human cells that undergo replicative aging, which correlates with increased occurrences of pathological protein aggregation. These findings establish previously unresolved critical crossing points for protein disaggregation, cell repair, and senescence.

### Heat shock triggers selective assembly of the Hsp70-DNAJA1-DNAJB1 trimeric chaperone complex

Previously, we tested the ability of four human class A and B JDP proteins in ten different configurations to assist Hsp70 (HSPA8) in solubilizing preformed aggregates of firefly luciferase biochemically (Supplementary Data Fig. 1a).^12^ A synergistic increase in disaggregation and refolding of aggregated luciferase was observed only for those JDP members that could form interclass JDP complexes with Hsp70 (Supplementary Data Fig. 1b), consistent with previous findings.^12^ The Hsp110 cochaperone, which functions primarily as a nucleotide exchange factor (NEF) for Hsp70, was added to reactions to power protein disaggregation efficiency.^25^

Next, we optimized and used the *in situ* proximity ligation assay (PLA)^14,26^ to trace the assembly of trimeric Hsp70 disaggregases in human cells (Fig. 1a). After signal amplification, six intra- and interclass JDP assemblies could be detected in unstressed (U) HeLa cells, albeit at low frequencies (yellow, false-colored puncta; Fig. 1b; Supplementary Data Fig. 2a). Controls consisting of only one of the JDP-specific primary antibodies did not generate fluorescent puncta (Supplementary Data Fig. 2b). The specificity of the interaction was further confirmed by depleting the JDP members using RNAi, which strongly decreased the PLA signal (Supplementary Data Fig. 2c, d). To induce protein aggregation, we next applied a proteotoxic stress using a 60 min heat shock (HS) at 43°C (Supplementary Data Fig. 2e). This led to a significant increase in the assembly of the DNAJA1-DNAJB1 scaffolds, exclusively (Fig. 1b; Supplementary Data Fig. 2a). We also observed an increase in the interaction between Hsp70 and the two JDPs on a very similar time scale (Fig. 1c). A mutant of DNAJB1 (DNAJB1^RRR^; RRR denotes D4R, E69R, and E70R) that we previously identified as defective in interacting with class A JDPs for disaggregase function *in vitro*,^12^ but fully active in assisting other functions of Hsp70 (e.g., refolding of heat-denatured luciferase^12^ and capable of shuttling into and out of nucleoli during resolution of heat-denatured proteins (Supplementary Data Fig. 3a)),^27–29^ showed a strong decrease in PLA foci formation (Supplementary Data Fig. 3b). A similar observation was made for a DNAJB1 mutant with a disrupted HPD motif (DNAJB1^H32Q^) that cannot bind to Hsp70 (Supplementary Data Fig. 3c).^30^ Together these data indicate that PLA captures direct interactions between specific JDPs and Hsp70s that appear to assemble a trimeric chaperone complex in cells with the same configuration as the biochemically identified Hsp70-DNAJA1-DNAJB1 disaggregase complex (Supplementary Data Fig. 1b). Disassembly of this trimeric chaperone complex to the basal levels occurred within 24h after HS (Fig. 1b, c). Disassembly was delayed in cells that were simultaneously depleted of all three Hsp110-type NEFs that were previously described to specifically increase the efficacy of protein disaggregation, *in vitro*.^25^ Depleting other types of NEFs such as BAG1 and HSPBP1 had no effect (Supplementary Data Fig. 4a, b). This further suggests that the activity of Hsp70-DNAJA1-DNAJB1 complex is related to disaggregase function in cells. The scaffold was also assembled upon relatively mild temperature elevations that resemble mild-moderate fever (37 °C to 39 °C) although to a lesser degree and shorter induction time (Fig. 1d; Supplementary Data Fig. 4c), supporting its physiological relevance.

**Figure 1.**
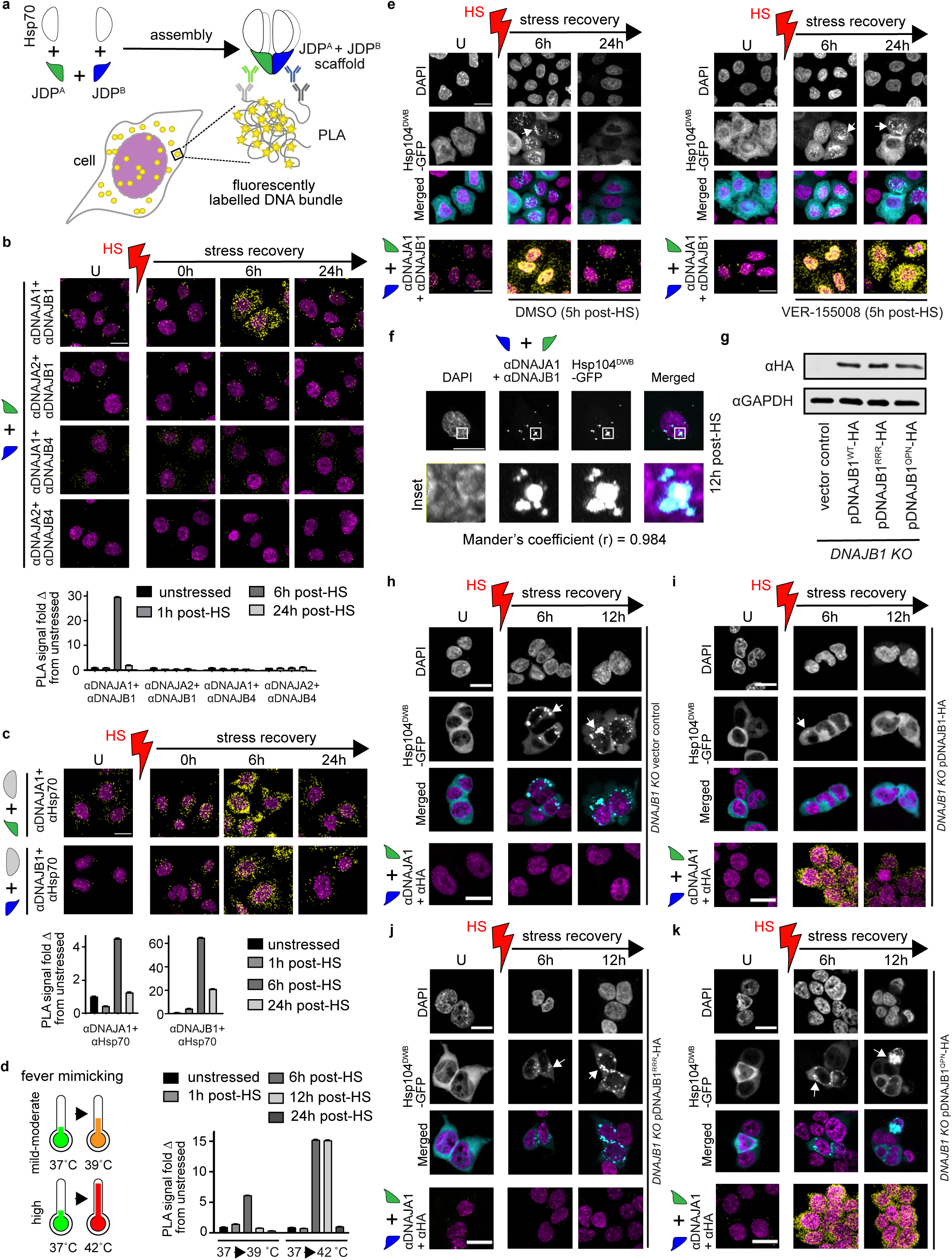
The Hsp70-DNAJA1-DNAJB1 trimeric complex forms a heat-induced protein disaggregase which targets aggregates that escape early clearance. **a**, Detection of Hsp70-JDP and JDP-JDP interactions during the assembly of Hsp70-DNAJA1-DNAJB1 trimeric complex *in situ* through proximity ligation assay (PLA). Captured protein-protein interactions generate fluorescent puncta (yellow) after signal amplification by rolling circle PCR. Fluorescently labeled oligonucleotides hybridize with an amplified DNA bundle to enhance the proximity signal. **b**, Top: Detection of multiple JDP^A^-JDP^B^ scaffolds that support *in vitro* Hsp70 disaggregase activity in unstressed (U) and heat shocked (HS, red) HeLa cells using PLA. Upon HS, cells show selective induction of the DNAJA1-DNAJB1 scaffold (PLA fluorescent signal, yellow). Nuclei stained with DAPI (magenta). HS was performed at 43 ° C for 1h; Stress recovery at 37 ° C for 0h, 6h and 24h. Scale bar 20 µm (n = 3). Images are maximum intensity projections (all channels). Bottom: Quantification of PLA signal from (data are mean +/− s.e.m.; see supplementary methods for details). **c**, Top: PLA-based detection of DNAJA1-Hsp70 and DNAJB1-Hsp70 interactions during the formation of Hsp70-DNAJA1-DNAJB1 trimeric complex. The Hsp70 antibody used in PLA recognizes both constitutively expressed Hsc70 (HAPA8) and the stress-induced Hsp70 paralog (HSPA1A). Scale bar 20 µm (n = 3). Bottom: Quantification of PLA signal (data are mean +/− s.e.m.). **d**, Assembly of DNAJA1-DNAJB1 scaffold at 39 °C *vs* 42 °C for 2h that mimic mild-moderate and high fever, respectively. Quantification of PLA signals (data are mean +/− s.e.m.). **e**, Resolution of heat-induced aggregates in cells +/− Hsp70 inhibitor VER-155008. Top: HS triggered the formation and solubilization of Hsp104^DWB^-GFP positive aggregates (cyan; punctated fluorescent signal pointed by white arrows) in HeLa cells. U denotes unstressed cells. Heat shock (HS) in red. Scale bar 20 µm (n = 3). Bottom: Visualization of assembly and disassembly of Hsp70 disaggregase through formation of the JDP scaffold (PLA; yellow) in HeLa cells recovering from heat shock (HS). Nuclei in magenta. Scale bar 20 µm (n = 3). The inhibitor was added to cells 5h after HS. Scale bar 20 µm (n = 3). DMSO, vehicle control for VER-155008. **f**, Assembly of DNAJA1-DNAJB1 scaffolds (PLA, yellow, see Methods) on Hsp104^DWB^-GFP positive heat-induced aggregates (cyan) during HS recovery (37 ° C for 12h) in HeLa cells. Inset shows zoomed in region of interest. Cells were treated with cycloheximide (CHX) at 3.5h and VER-155008 at 5.5h after HS. Mander’s coefficient (r) = 0.984 (s.e.m. = 0.004). Manders’ overlap correlation coefficients were calculated using thresholded images (n=5). Nuclei in magenta. Scale bar 20 µm. **g**, Immunoblot showing expression levels of wild type DNAJB1-HA, DNAJB1^RRR^-HA and Hsp70-binding defective DNAJB1^QPN^-HA (H32Q+D34N) mutant in *DNAJB1* knocked out (KO) HEK 293 cells. **h-k**, Defective assembly of Hsp70-DNAJA1-DNAJB1 disaggregase results in persistence of heat-induced aggregates in cells. Top: Visualization of solubilization of Hsp104^DWB^-GFP positive heat-induced aggregates during HS recovery in *DNAJB1* KO HEK 293 cells ectopically expressing (**h**) empty vector (control), (**i**) wild type DNAJB1-HA, (**j**) DNAJB1^RRR^-HA, and (**k**) DNAJB1^QPN^-HA. Bottom: (Dis)assembly dynamics of DNAJA1-DNAJB1 scaffold by PLA (yellow) in *DNAJB1* KO HEK 293 cells ectopically expressing (**h**) empty vector (control), (**i**) wild type DNAJB1-HA, (**j**) DNAJB1^RRR^-HA, and (k) DNAJB1^QPN^-HA. U denotes unstressed cells. Scale bar 20 µm (n = 3).

### The late induction of Hsp70-DNAJA1-DNAJB1 disaggregase is geared to specifically target a distinct population of aggregates in human cells

The delayed induction of the Hsp70-DNAJA1-DNAJB1 trimeric complex during stress recovery (which peaked approximately at 6h post-HS) (Fig. 1b) was puzzling because heat-induced aggregate formation (measured as the appearance of denatured protein containing foci or insoluble detergent protein material) occurs almost immediately in cells. Many of these initial aggregating proteins are associated with or sequestered in biomolecular condensates such as nucleoli and stress granules (SG).^27–29,31^ Furthermore, the majority of detergent-insoluble material and condensates were found to disappear before the first 6 hours after HS (Supplementary Data Fig. 5a,b)^27,28,31,32^, *i.e.*, before the trimeric Hsp70-DNAJA1-DNAJB1 complex peaks in its assembly. Thus, the late induction of this complex might be functionally unrelated to the disassembly of these aggregated/ condensated proteins induced by HS. Therefore, we hypothesized that this trimeric Hsp70 disaggregase might be performing a unique task, perhaps targeting a different population of aggregated protein species, different from condensates that escape the early clearance phase. To detect whether such escaping aggregated species exist in human cells, we expressed the catalytically inactive yeast Hsp104 (E285Q, E687Q Double Walker B) tagged with GFP (Hsp104^DWB^-GFP) which decorates stress-induced aggregates.^33^ Hsp104^DWB^-GFP showed a diffuse fluorescent GFP signal in unstressed cells (cyan, false-colored; Fig. 1e; Supplementary Data Fig. 6a). Immediately after HS, fluorescent puncta (white arrows), denoting aggregates, emerged predominantly in the cytosol (Supplementary Data Fig. 6a). Notably, no enrichment of Hsp104^DWB^-GFP positive puncta was detected in nucleoli containing stress-induced condensates immediately after HS. At 6h after HS recovery, the number of Hsp104^DWB^-GFP positive aggregates increased in number and became more compact in appearance in cells. However, 24h after stress recovery, these aggregates were completely resolved (Fig. 1e; Supplementary Data Fig. 6a). The prevalence of Hsp104^DWB^-GFP positive aggregates was consistent with the timing of the assembly and disassembly of the Hsp70-DNAJA1-DNAJB1 complexes. To further test whether Hsp104^DWB^-GFP positive aggregates were targeted by the Hsp70-DNAJA1-DNAJB1 trimeric complex, we used VER-155008, a specific adenosine-derived small molecule inhibitor of Hsp70.^34^ VER-155008 significantly inhibited disaggregase activity mediated by Hsp70-DNAJA1-DNAJB1, *in vitro* (Supplementary Data Fig. 6b). The addition of VER-155008 to cells immediately after HS has been shown to block the early recovery of insoluble proteins accumulated in nucleoli.^27,28^ Therefore, to allow the completion of this recovery and specifically inhibit the activity of the Hsp70-DNAJA1-DNAJB1 trimeric complex, we added VER-155008 5h after HS. Treatment with VER-155008 completely blocked the disappearance of Hsp104^DWB^-GFP positive aggregates (Fig. 1e; Supplementary Data Fig. 6a). In line, DNAJA1-DNAJB1 scaffolds also persisted up to 24h after HS (Supplementary Data Fig. 6a). The presence of VER-155008 caused the PLA signal from DNAJA1 and DNAJB1 complex formation (Fig. 1f) to strongly overlap with Hsp104^DWB^-GFP positive foci (Mander’s overlap correlation coefficient (r) = 0.984), which is consistent with JDP scaffolds forming on aggregate surfaces to drive the assembly of this trimeric Hsp70 disaggregase during protein disaggregation. Washing out VER-155008 12h after HS led to re-initiation of Hsp104^DWB^-GFP positive aggregate resolution (Supplementary Data Fig. 7a, b). These results indicate that the disassembly of Hsp104^DWB^-GFP positive aggregates requires Hsp70 activity. The resolution of the aggregates in the washed cells VER-155008 coincided with the disassembly of the DNAJA1-DNAJB1 scaffolds (Supplementary Data Fig. 7c), suggesting that they help recruit active Hsp70 to form disaggregases on the aggregates. To further test this hypothesis, we expressed wildtype yeast Hsp104, which cooperates with the mammalian Hsp70 system to form very potent disaggregases *in vitro* (Supplementary Data Fig. 8a) and *in vivo*,^35^ in HeLa cells. The frequency of DNAJA1-DNAJB1 scaffolds was strongly reduced in cells expressing Hsp104-GFP 6h after HS (Supplementary Data Fig. 8b) due to a rapid decrease in aggregate load (see below section; Supplementary Data Fig. 8c), indicating a functional redundancy between Hsp104 and the trimeric Hsp70 disaggregase systems. Furthermore, the data suggested that the presence of aggregates could also be a trigger for the assembly of DNAJA1-DNAJB1 scaffolds.

Next, we used DNAJB1 mutants defective in assembling the trimeric complex to assess whether JDP scaffolding is required to solubilize HS-induced Hsp104^DWB^-GFP positive aggregates. The *in vitro* disaggregation of heat aggregated luciferase was significantly reduced in reactions containing DNAJB1^RRR^ and DNAJA1 (Supplementary Data Fig. 9a). Accordingly, ectopically expressed DNAJB1^RRR^ in *DNAJB1* knocked out (KO) HEK 293 cells (confirmed by immunoblotting; Fig. 1g) was unable to solubilize Hsp104^DWB^-GFP positive aggregates induced by HS compared to wild type DNAJB1 (Fig. 1h-j; Supplementary Data Fig. 9b-d). A similar result was observed with DNAJB1^H32Q, D34N^ (DNAJB1^QPN^), which is defective in binding to Hsp70. However, DNAJB1^QPN^ readily complexed with endogenous DNAJA1 after HS (Fig. 1g, k), but these JDP scaffolds could not support disaggregation, presumably due to the decreased ability to recruit Hsp70, and as a consequence prevented efficient aggregate resolution in recovering cells (Fig. 1k, Supplementary Data Fig. 9c). Together, these findings support that the Hsp70-DNAJA1-DNAJB1 trimeric complex represents a real, functional disaggregase in mammalian cells, tentatively crucial for solubilizing a population of stress-induced aggregates that are distinct from those accumulated in condensates.

To investigate whether the DNAJA1-DNAJB1 scaffold is interchangeable with other class A and B members as observed in biochemically reconstituted systems (Supplementary Data Fig. 1),^12^ we knocked down DNAJA1 or DNAJB1 in HeLa cells using RNAi, which, as expected, decreased the assembly of the JDP scaffold after HS. However, knocking down DNAJA1 or DNAJB1 did not result in the remaining unpaired partner complexing with other JDPs (e.g., DNAJB1 with DNAJA2 or DNAJA1 with DNAJB4) in cells (Supplementary Data Fig. 10a). Thus, regardless of the expression and HS induction of other JDP paralogs (Supplementary Data Fig. 10b) in mutual subcellular locations (Supplementary Data Fig. 11), this particular Hsp70 disaggregase is cogged to assemble with DNAJA1 and DNAJB1 during the late phase of stress recovery in human cells. In line with this, DNAJB1 activity could not be fully restored in *DNAJB1* KO cells by overexpressing other JDPs tested (Supplementary Data Fig. 9e). These data highlight the strict dependency on DNAJB1 for the functionality of this disaggregase. The observations also indicate that single JDP-containing disaggregases described biochemically^14,36–39^ are ineffective in solubilizing this persisting fraction of heat-induced aggregates in human cells.

To identify the protein composition of the aggregates targeted by Hsp70-DNAJA1-DNAJB1 disaggregase in HeLa cells, we applied a two-phase fractionation approach coupled to a proteomic workflow (Supplementary Data Fig. 12a).^19^ Data were agnostically clustered into four different response patterns by k-means clustering (see Methods; Supplementary Data Table 1). The proteins in cluster (iii) showed a pattern of aggregation and recovery that was consistent with being potential substrates for this disaggregase activity. Specifically, they appeared to show impaired solubility recovery after treatment with VER-155008 between 6 and 12 hours (Supplementary Data Fig. 12b, c). Approximately half of the hits in this cluster included RNA-associated proteins (Supplementary Data Table 1). Two proteins in cluster (iii), Heterogeneous nuclear ribonucleoprotein K (HNRNPK; solubility profile in Fig. 2a) and Adenosylhomocysteinase (AHCY; solubility profile in Fig. 2b), were selected for further analysis. Both HNRNPK and AHCY formed puncta that persisted at 6h after HS in HeLa cells, but they were completely solubilized and redistributed to original subcellular locations at 12h after stress recovery (Fig. 2c, d). The addition of VER-155008 (Fig. 2c, d; Supplementary Data Fig. 12d) as well as the knocking down of DNAJA1 or DNAJB1 (Fig. 2e) prevented this resolubilization. Fluorescent recovery after photobleaching (FRAP) analysis showed that the HNRNPK puncta were immobile, suggesting tight protein packing in these aggregates (Supplementary Data Fig. 12e). The formation of these HS-induced immobile HNRNPK puncta was completely suppressed after expressing the yeast Hsp104 disaggregase (Supplementary Data Fig. 8c). This explains the reduced assembly of the trimeric disaggregase after HS in cells ectopically expressing Hsp104 (Supplementary Data Fig. 8b).

**Figure 2.**
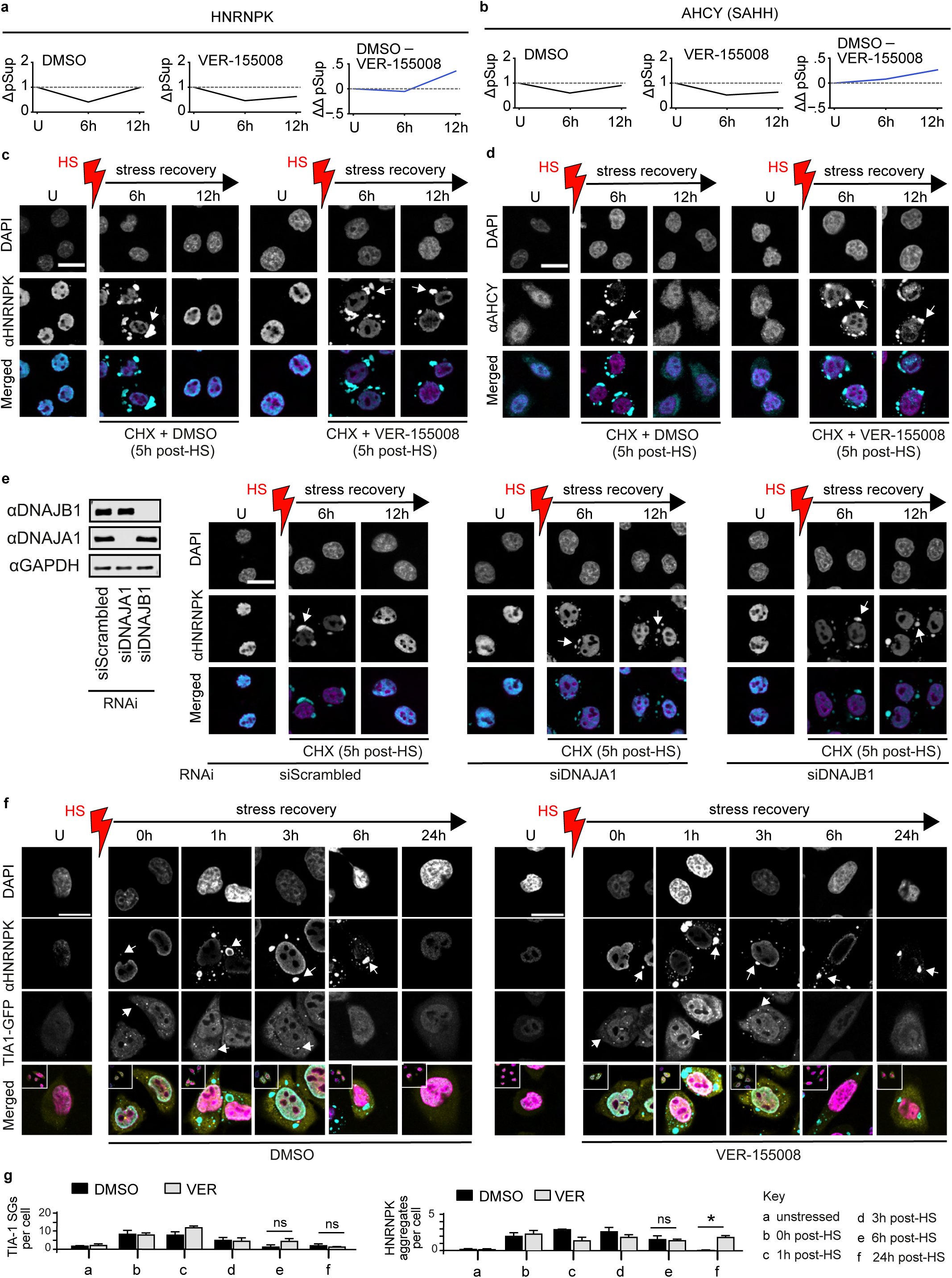
Endogenous substrates of the Hsp70-DNAJA1-DNAJB1 disaggregase. **a-d,** Solubilization of heat aggregated HNRNPK and AHCY (SAHH) requires the induction of Hsp70-DNAJA1-DNAJB1 disaggregase. **a,** Solubility profile of HNRNPK: Graphs show representative traces for ΔpSup in the presence of DMSO (left panel) or VER-155008 (center panel), and the corresponding ΔΔpSup (right panel) (n = 3). See methods for the two-phase fractionation approach coupled to a proteomic workflow used to identify aggregated substrates recovered by Hsp70-disaggregase activity. **b,** As in (**a**), for AHCY (n = 3). **c,** Solubilization of heat aggregated HNRNPK in HeLa cells visualized using immunocytochemistry (HNRNPK, cyan; DAPI stained nuclei, magenta). Hsp70 is inhibited with VER-155008 added 5h after HS. Cycloheximide (CHX) was added to inhibit protein synthesis at 5h after HS. White arrows indicate heat-aggregated HNRNPK. Aggregate solubilization is indicated by the disappearance of the punctated fluorescence signal and the appearance of a diffused signal. DMSO, vehicle control for VER-155008. Heat shock (HS) in red. U denotes unstressed cells. Scale bar 20 µm (n = 3). **d,** As in (**c**), for AHCY (n = 3). **e,** Disruption of DNAJA1-DNAJB1 JDP scaffold formation prevents solubilization of heat-aggregated HNRNPK. Left: Immunoblot showing JDP knockdown efficiencies in HeLa cells. Right: Persistence of heat aggregated HNRNPK in DNAJA1 and DNAJB1 depleted HeLa cells. The non-targeting control siRNA (siScrambled) is used as the control for RNAi. Scale bar 20 µm (n = 3). **f,** Stress-induced Hsp70 disaggregase targets different aggregate types in cells. Top: Hsp70-depended solubilization of HNRNPK aggregates (cyan) *vs* TIA-1-GFP containing SGs (yellow) in HeLa cells recovering from heat stress. The addition of VER-155008 at 0h after HS blocks resolution of HNRNPK aggregates, but not TIA-1 SGs. The inset depicts zoomed-out cells. Scale bar 20 µm. **g,** Quantification of TIA-1 SGs and HNRNPK aggregates (n = 3, data are mean +/− s.e.m. * adjusted p-value < 0.05, one-way ANOVA LSD post hoc test).

### Cellular heterogeneity in aggregate solubilization by sequentially activated Hsp70-DNAJA1-DNAJB1 and VCP-based disaggregase systems

The assembly and disassembly kinetics of the Hsp70-DNAJA1-DNAJB1 disaggregase and the disaggregation of HNRNPK and AHCY are highly distinct from the recovery of protein aggregates that accumulate in biomolecular condensates after HS. Though HNRNPK and TIA-1 are both RNA-binding proteins, they sequestered into spatially distinct aggregates (HNRNPK; cyan) and SGs (TIA-1; yellow) in the cytosol after HS (Fig. 2f). Compared to HNRNPK aggregates, the disassembly of the TIA-1 SGs occurred rapidly within 6h after HS and did not rely on Hsp70-DNAJA1-DNAJB1 disaggregase activity, as evident from VER-155008 treatment and JDP knockdown experiments (Fig. 2f, g; Supplementary Data Fig. 13), suggesting that these SGs are processed through a distinct nonoverlapping disaggregase activity. We also noted the presence of a fraction of highly ubiquitinylated proteins that immediately insolubilized, but rapidly resolved before 6h into recovery from HS (Supplementary data Fig 5a, right panel). Recent findings show that AAA+ ATPase VCP (p97) is not only recruited to ubiquitylated substrates, but is also required for their disaggregation in cooperation with the ubiquitin proteasome machinery.^22–24^ Furthermore, VCP was found to be involved in SG clearance in mammalian cells.^40^ To test whether VCP targets the heat-induced Ub+ insolubilize protein fraction, we depleted the AAA+ ATPase using RNAi, which indeed resulted in a considerable delay in resolving the Ub+ aggregated proteins (Fig. 3a) and TIA-1 SGs (Fig. 3b, c). Surprisingly, however, VCP depletion also delayed the solubilization of HNRNPK and Hsp104^DWB^-GFP positive aggregates (Fig. 3b, c; Supplementary Data Fig. 14a) that largely lacked an Ub signature (Supplementary Data Fig. 14b). In line, the disassembly of the Hsp70-DNAJA1-DNAJB1 disaggregase was also delayed after VCP knockdown (Supplementary Data Fig. 14c). To test whether loss of VCP directly affected the action of Hsp70 disaggregase or whether VCP itself also facilitated the resolution of non-condensate-associated Hsp104^DWB^-GFP positive aggregates, we treated cells with the VCP inhibitor NMS-873^[41]^ at 0h or 5h after HS. Adding NMS-873 at 0h post-HS triggered the same phenotype as that of depleting VCP. However, blocking VCP function using a low (1 µM) or high (10 µM) dose of NMS-873 5h after HS, *i.e.* after clearing of Ub+ aggregates, did not affect Hsp104^DWB^-GFP positive aggregate solubilization (Fig. 3d; Supplementary Data Fig. 14d). Consequently, under this condition, the Hsp70-DNAJA1-DNAJB1 disaggregase disassembled back to basal levels at 24h after HS (Fig. 3e). Inversely, increasing VCP levels did not affect the activation of the Hsp70-DNAJA1-DNAJB1 disaggregase during stress recovery (Supplementary Data Fig. 15), which is in stark contrast to observations made under overexpression of the yeast Hsp104 disaggregase (Supplementary Data Fig. 8). These observations suggest that the function of trimeric Hsp70 disaggregase cannot be compensated for by stimulating the function of VCP in human cells. Together, these data describe for the first time how protein disaggregation activity is organized into two phases (early and late) in living human cells undergoing stress recovery. The stress-induced trimeric complex Hsp70-DNAJA1-DNAJB1 forms a crucial part of an integrated network of at least two distinct disaggregase systems that function sequentially and in a non-redundant manner to target different populations of heat-induced aggregated substrates.

**Figure 3.**
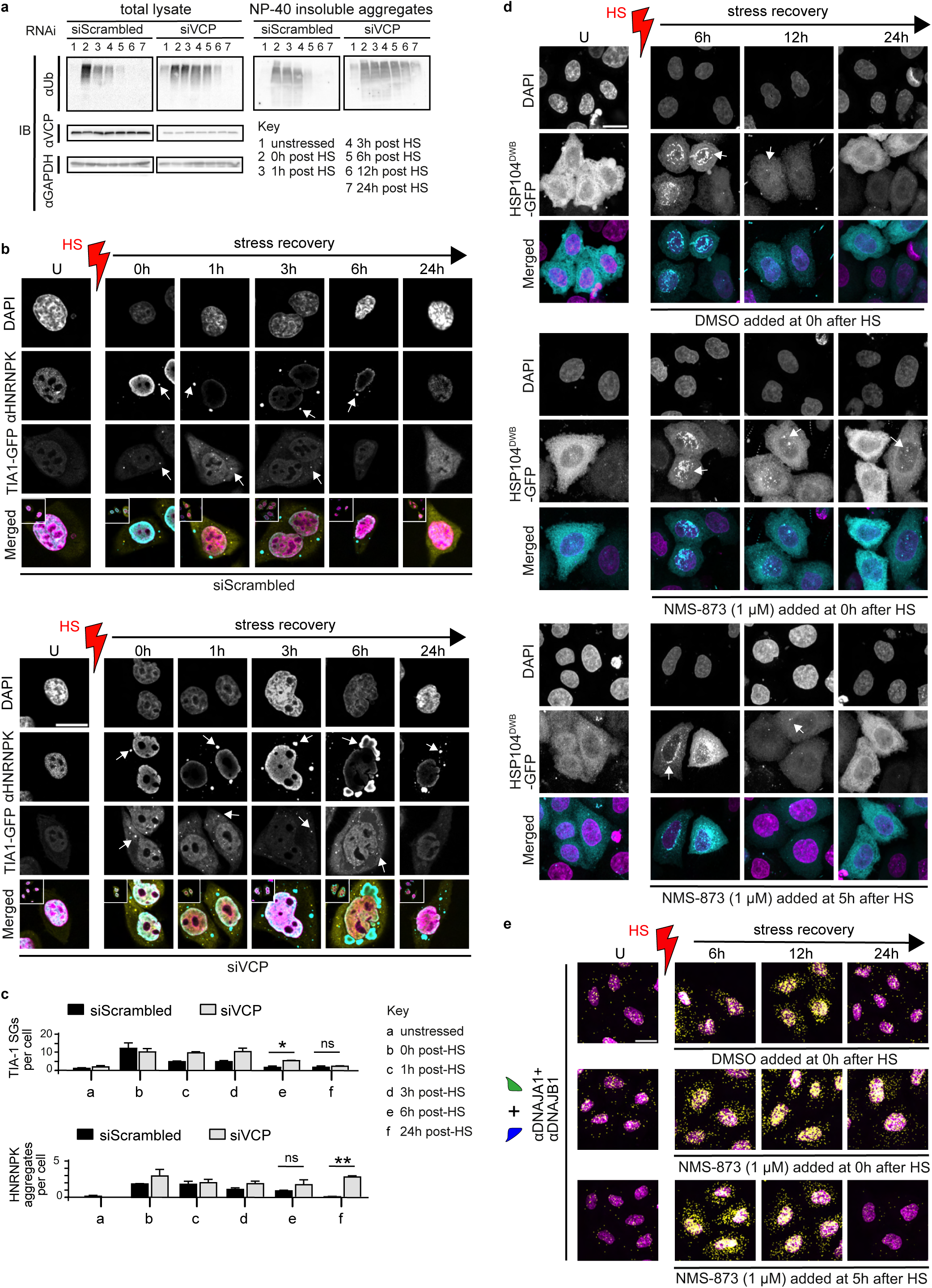
VCP and Hsp70-DNAJA1-DNAJB1 disaggregases operate sequentially and nonredundantly in cells recovering from heat stress. **a,** Disaggregase activity exerted by VCP facilitates rapid resolution of heat-induced ubiquitin (Ub) positive protein aggregates. 1% NP-40 insoluble aggregates were isolated at indicated time points during heat stress recovery using a two-phase fractionation approach and probed with antibody against ubiquitin (n = 3). Immunoblots show depletion of VCP delays the disaggregation and clearance of Ub positive protein aggregates in HeLa cells. VCP protein levels show knockdown efficiency. GAPDH, loading control. **b,** Solubilization of heat-induced HNRNPK aggregates (cyan) and TIA-1-GFP containing SGs (yellow) are affected in VCP depleted HeLa cells. The inset depicts zoomed-out cells. The non-targeting control siRNA (siScrambled) is used as the control for RNAi (same from Supplementary Data Fig. 13). Nuclei stained with DAPI (magenta). Scale bar 20 µm. **c,** Quantification of TIA-1 SGs and HNRNPK aggregates (n = 3, data are mean +/− s.e.m. * adjusted p-value < 0.05, ** adjusted p-value < 0.01, ns - not significant, one-way ANOVA LSD post hoc test). **d,** Blocking VCP disaggregase activity with inhibitor NMS-873 (1 µM, low dose) after 5h post-HS does not inhibit the resolution of Hsp104^DWB^-GFP positive aggregates by Hsp70 disaggregase. On the contrary, the addition of NMS-873 at 0h post-HS results in a phenotype similar to the depletion of VCP. DMSO, vehicle control for the inhibitor (n = 2). Nuclei stained with DAPI (magenta). **e,** Disassembly of DNAJA1-DNAJB1 scaffold is delayed in HeLa cells treated with NMS-873 at 0h post-HS, but not at 5h post-HS (PLA signal, yellow). Nuclei stained with DAPI (magenta). Heat shock (HS) in red. U denotes unstressed cells. Scale bar 20 µm (n = 3).

### Regulation of Hsp70-DNAJA1-DNAJB1 disaggregase

Next, we asked how the kinetics of these sequentially acting disaggregases are regulated by stress signaling. VCP, which acts at the early phase of aggregate solubilization, is not induced by proteotoxic stresses (Fig. 3a, left panel)^42^ and knocking down its levels prior to heat stress affects the disassembly of Ub+ aggregates/SGs (Fig. 3a,b). On the contrary, the Hsp70-DNAJA1-DNAJB1 disaggregase, which shows low abundance in unstressed cells, is massively induced after HS, indicating that this disaggregase is highly regulated. In HeLa cells, the peak assembly of the DNAJA1-DNAJB1 scaffold (Fig. 1b), which initiates the assembly process and provides specificity for this PQC machine, coincided with the highest expression of the disaggregase components in protein (Supplementary Data Fig. 16a), but not mRNA levels (Supplementary Data Fig. 16b). Induction of machine components after stress suggests that the heat shock response (HSR) might modulate the formation of this disaggregase in repairing cells. Knocking down the HSF-1 transcription factor, the central regulator of the HSR, inhibited the synthesis of disaggregase components and reduced the assembly of the disaggregase (Supplementary Data Fig. 16c) and subsequent resolution of its substrates (Supplementary Data Fig. 16d; Supplementary Data Fig. 17a). In contrast, HSF-1 knockdown had no effect on the VCP-dependent disassembly of TIA-1 SGs (Supplementary Data Fig. 17a) again highlighting the nonoverlapping disaggregase activities of the two systems. Similar to HSF-1 depletion, treatment of cells with the translational inhibitor cycloheximide (CHX) immediately after HS also blocked the assembly of Hsp70-DNAJA1-DNAJB1 disaggregase (Supplementary Data Fig. 17b). In contrast, HSF-1 depletion or CHX treatment did not affect the HS-induced localization dynamics of pre-HS synthesized Hsp70s and JDPs (Supplementary Data Fig. 18; Supplementary Data Fig. 19) coinciding with the extraction of denatured/aggregated proteins from nucleoli SGs immediately after HS.^27–29^ This implies that the late phase, but not the early phase of aggregate solubilization is under tight transcriptional regulation. This regulation also allows the assembly of spatially and temporally separated Hsp70 machines through pre- and post-HS synthesized DNAJA1 and DNAJB1 molecules. In essence, this serves as a strategy to efficiently organize different chaperone machines that cater to various cellular needs on a priority basis, for example, to promote rapid recovery of proteins associated with translational machinery crucial for immediate stress adaptation^43–46^ followed by resolving persistent aggregated protein junk to reduce downstream cytotoxicity.

### Physiological relevance of Hsp70-DNAJA1-DNAJB1 disaggregase: heat resistance, replicative aging, and pathological protein aggregation

To address the physiological relevance of the Hsp70-DNAJA1-DNAJB1 disaggregase, we first evaluated the impact of its assembly on the fitness of human cells recovering from heat stress using the lactate dehydrogenase (LDH)-Glo-based cytotoxicity assay (Supplementary Data Fig. 20a). Again, VER-155008 was added after the completion of rapid recovery of heat-denatured proteins from nucleoli immediately after HS.^29^ The Hsp70-DNAJA1-DNAJB1 disaggregase activity inhibited HeLa cells (Supplementary Data Fig. 20b) showed increased heat toxicity compared to control cells treated with DMSO (Supplementary Data Fig. 20c). This shows that the resolution of aggregates escaping early clearance contributes significantly to the cellular tolerance of acute proteotoxic stress.

To address whether the trimeric Hsp70-DNAJA1-DNAJB1 disaggregase also plays a role in chronic stress conditions resulting from aging and age-related diseases, we first assessed its assembly in six lines of primary dermal fibroblasts (HDF) derived from humans with ages ranging from 21 to 75 years. We did not observe a substantial chronological age-driven variation in the inducibility of Hsp70-DNAJA1-DNAJB1 disaggregase after HS in all fibroblast lines tested (Fig. 4a; Supplementary Data Fig. 21a). Similar to immortalized HeLa cells, primary dermal fibroblasts also only induced the DNAJA1-DNAJB1 scaffold after HS (Supplementary Data Fig. 21b), underscoring that this assembly, with the same kinetics and specific components, is also active in normal human diploid cells.

**Figure 4.**
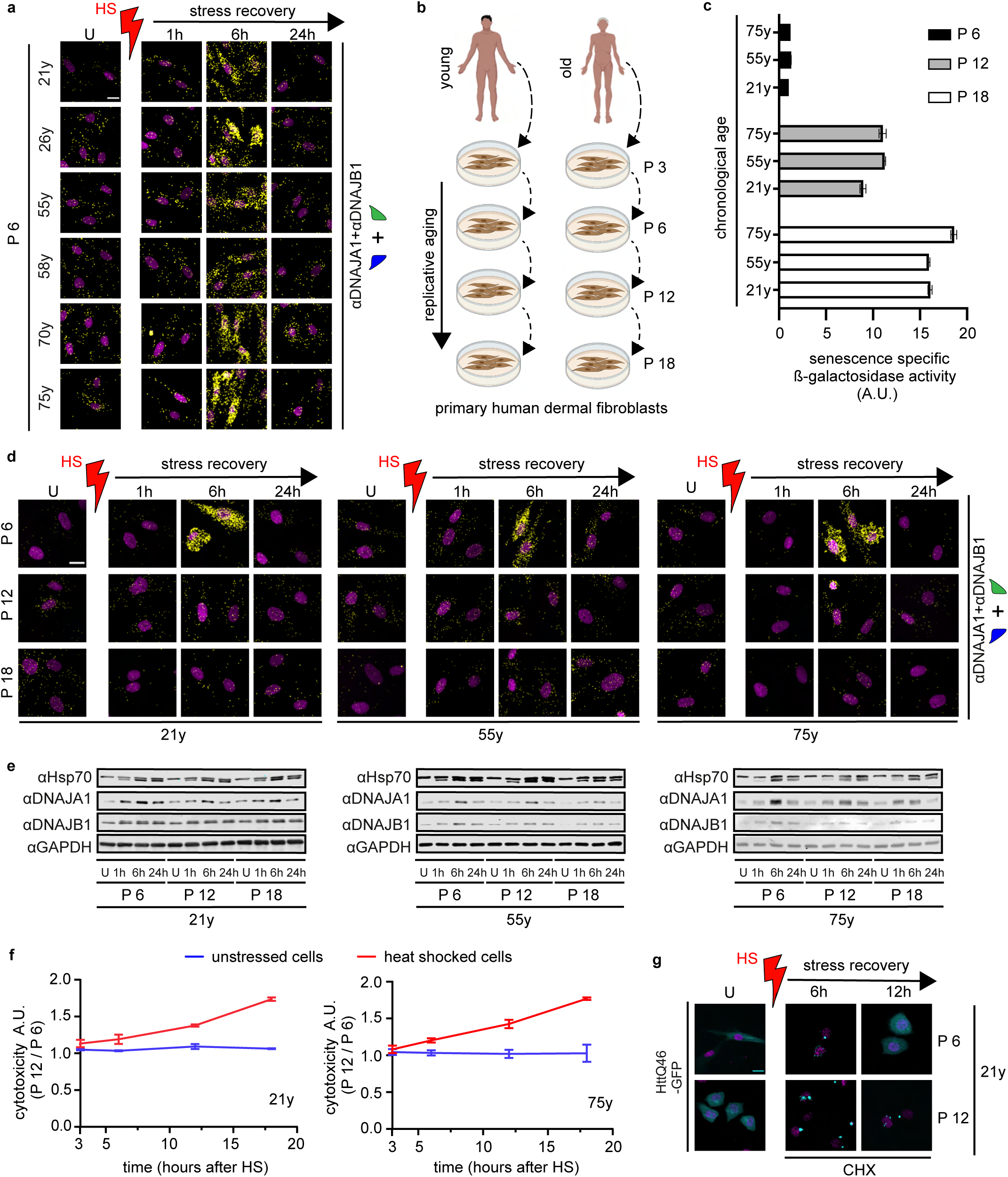
Pathophysiological consequences of the collapse of Hsp70-DNAJA1-DNAJB1 disaggregase in cells undergoing replicative aging. **a,** Chronological age has no effect on Hsp70-DNAJA1-DNAJB1 disaggregase activation in human cells. The PLA signal (yellow) shows the induction of Hsp70-DNAJA1-DNAJB1 disaggregase in heat-stressed primary human dermal fibroblasts derived from 21-year-old (21-y), 55-y and 75-y human subjects. Nuclei stained with DAPI (magenta). U denotes unstressed cells. Heat shock (HS) in red. Scale bar 20 µm (n = 3). **b,** Experimental scheme showing replicative aging of human dermal fibroblasts. P denotes cell passage (see Methods). **c,** Cells at P = 12 and 18 show a considerable increase in the marker of senescence. Cellular senescence correlates with the increased levels of senescence specific β-galactosidase activity in replicating fibroblasts. **d,** HS-induced assembly of Hsp70 disaggregase collapses in cells undergoing replicative aging. The level of induction of Hsp70 disaggregase assemblies after HS in human dermal fibroblasts at P = 6, 12 and 18 using PLA. Scale bar 20 µm (n = 3). **e,** Immunoblot analysis of the induction of components of the disaggregase machine after HS (n = 3). **f,** Cells defective in activating Hsp70-DNAJA1-DNAJB1 disaggregase due to aging (replicative) show reduced cellular fitness after HS. Level of cytotoxicity measured with Ultra-Glo rLuciferase-based cytotoxicity assay in unstressed (blue) and HS recovering (red) human dermal fibroblasts at P 6 and P 12 cells. Error bars depicted as s.e.m. (n = 3). **g,** Human dermal fibroblasts at P 12 lose the ability to solubilize Htt46Q-GFP aggregates (cyan) by assembling Hsp70-DNAJA1-DNAJB1 disaggregase. Cycloheximide (CHX) was added to prevent protein synthesis. Scale bar 20 µm (n = 3).

Recent work shows that during chronological aging, senescent cells resulting from replicative aging accumulate progressively in various mammalian tissues.^47–49^ Therefore, we evaluated whether replicative aging has an effect on activating the Hsp70 disaggregase. Three of the human dermal fibroblast lines (derived from 21-, 55-, and 75-year-old donors) were subjected to replicative aging by *ex vivo* passaging (P) of cells up to 18 times (Fig. 4b, see Methods). Senescence was detected using lysate-based SPiDER-Gal activity assay and direct cell staining for senescence-specific β-galactosidases levels. Compared to non-senescent P 6 cells, cells in P 12 showed a rise in the senescent signal, which increased further in P 18 cells in all lines, regardless of the chronological age of the donor (Fig. 4c; Supplementary Data Fig. 21c, d). Strikingly, the stress-induced assembly of Hsp70-DNAJA1-DNAJB1 disaggregase observed in cells at P 3 or P 6 was completely absent in cells at P 12 and P 18 (Fig. 4d). Importantly, no notable differences were observed in the heat inducibility of disaggregase components between P 6 and P 12 cells (Fig. 4e). Consistently, HSR, which is reported to collapse in cells at late senescence with permanent cell cycle arrest,^50,51^ was still fully intact in still replicating P 12 cells, as indicated by robust induction of the HSF-1 dependent HSPA6 chaperone^52^ (Supplementary Data Fig. 22a). This finding argues that additional pathways beyond the HSR are required for the assembly of the Hsp70-DNAJA1-DNAJB1 disaggregase. Furthermore, as in P 6 cells, P 12 cells also showed similar levels of heat-induced i) nuclear shuttling of ATF-6 to activate the unfolded protein response of the endoplasmic reticulum (UPR_ER_) (Supplementary Data Fig. 22b), and ii) proteasome activity (Supplementary Data Fig. 22c) that decline in human cells at late senescence.^53^ We also did not observe any decrease in VCP levels in fibroblasts in P 6 vs P 12 (Supplementary Data Fig. 22a). Together, these intriguing findings indicate that Hsp70-DNAJA1-DNAJB1 disaggregase is a PQC component that declines very rapidly with replicative aging. The early decline of this disaggregase activity has functional consequences as P 12 fibroblasts showed an inability to resolve heat-induced protein aggregates (Supplementary Data Fig. 23a, b), showed reduced fitness after HS (Fig. 4f), and contained aggregated proteins even under non-stressed conditions (Supplementary Data Fig. 23b), a hallmark of aging cells.

Replicative aging, which drives chronic senescence, has been implicated in the etiopathology of several neurodegenerative diseases.^47^ Therefore, we evaluated how impairment of this Hsp70 disaggregase during replicative aging affects the aggregation of pathological proteins associated with Huntington’s disease (HD). Appearance of intracellular Huntingtin protein (Htt) exon 1 containing expanded polyglutamine (polyQ) amyloid aggregates drives neurotoxicity in HD.^2^ These aggregates form in cells independent of proteotoxic stresses such as HS (Supplementary Data Fig. 24a). Our findings showed that priming of Hsp70-DNAJA1-DNAJB1 disaggregase assembly through HS reduced the accumulation of polyQ aggregates (HttQ46-mEmGFP) in a Hsp70, DNAJA1, and DNAJB1 dependent manner in HeLa cells (Supplementary information 1, Supplementary Data Fig. 24). Unlike HeLa cells, Htt46Q-mEmGFP was largely soluble with only a mild aggregation phenotype in unstressed dermal fibroblasts at P 6 and P 12 (Fig. 4g). However, after HS, the soluble Htt46Q-mEmGFP formed into prominent aggregates (punctated cyan fluorescent signal, 6h after HS) in both P 6 and P 12 cells, which is in line with an acute overloading of the PQC pathways that prevents the initiation of polyQ aggregation in stressed cells. Strikingly, these polyQ aggregates subsequently disappeared in P 6, but not P 12 fibroblasts (Fig. 4g) that can activate the trimeric Hsp70 disaggregase. These proof-of-principle findings unravel a critical breakdown of amyloidogenesis suppression in cells undergoing senescence. Together, our data reveal that the *in vivo* assembly of this Hsp70-DNAJA1-DNAJB1 trimeric chaperone complex vitally contributes to cellular resistance to stress and declines rapidly during replicative aging with putative relevance to disease-related amyloidosis.

## Discussion

The critical role of recovery of aggregated proteins through Hsp100-based disaggregation activity in unicellular organisms such as bacteria and yeast after heat stress has been widely accepted.^54–56^ Whilst metazoa lack cytosolic/nuclear Hsp100 disaggregases,^10^ a number of recent studies suggested that these organisms possess several functionally equivalent protein disaggregase systems composed of different proteins with chaperone or chaperone-like properties (e.g. Hsp70, VCP, and DAXX).^12,23,57^ Importantly, however, it was not understood how these different protein disaggregase systems are assembled (if at all) and regulated similarly or differently in time and space in live cells exposed to protein aggregation stresses. It was also not understood how these disaggregase systems are organized (if at all) in cells during cell repair. Here, we describe for the first time how proteotoxic stress induces the selective assembly of an *in vitro* identified Hsp70-based disaggregase with a specific DNAJA1 and DNAJB1 containing JDP scaffold in primary and immortalized human cells. We show that this trimeric disaggregase operates in the late phase of aggregate solubilization sequentially to the action of a VCP-based disaggregase. The early phase of aggregate solubilization is predominated by the activity of VCP-based disaggregase. VCP largely depends on a Ub signal to recognize its aggregates/SG components, but could also function independently of Ub to disassemble some protein complexes.^58^ Following the initial aggregate solubilization phase, the trimeric Hsp70 disaggregase assembles on remaining aggregates. The data suggest that the Hsp70 disaggregase is particularly effective in “mopping up” tightly packed solid-like cytotoxic protein aggregates that were not sequestered into condensates (e.g., SG or nucleoli) and hence escaped the early phase of clearance. Although the late phase depends on the completion of the early phase, it does not require VCP. In terms of the fate of disaggregated polypeptides, there is likely a culmination of protein recovery and degradation during both phases.^59–62^

The two disaggregase systems affect resistance to cellular stress and appear to be associated with age-associated human diseases. Recent work shows the ability of VCP to target tau amyloids associated with Alzheimer’s disease.^23^ Genetic defects in VCP activity were found to cause inclusion body myopathy associated with Paget’s disease of the bone and front-temporal dementia (FTD).^63^ Additionally, VCP activity was reported to be important for clearing stress granules,^40^ and mutations in VCP alter SG dynamics closely linked to ALS and FTD pathologies.^62^ Our data show that assembly of the Hsp70-DNAJA1-DNAJB1 disaggregase also leads to resistance against acute (heat) stresses. Once induced, this trimeric disaggregase is capable of targeting pathological amyloids (e.g., HttpolyQ amyloids associated with HD).

Importantly, we show that the ability to assemble the Hsp70-DNAJA1-DNAJB1 disaggregase is highly impaired during replicative aging that causes chronic senescence. Mechanistically, the breakdown of this disaggregase activity is due to the inability to effectively assemble the DNAJA1-DNAJB1 JDP scaffold. Importantly, our results revealed that the assembly of Hsp70-DNAJA1-DNAJB1 disaggregase is impaired even before human cells undergoing replicative aging lose several proteotoxic signaling routes, such as the HSR and the UPR_ER_, which have often been considered to precede the collapse of protein homeostasis upon aging.^53,64^ This early collapse could explain the accumulation of difficult-to-resolve aggregates in unstressed cells (Supplementary Data Fig. 23b) and may act as a trigger that contributes to (rather than being the result of) senescence, and thereby rapidly prevent damaged cells from propagation. Interestingly, compared to human cells, non-metazoans such as budding yeast, modulate protein disaggregation function entirely differently during replicative aging. Instead, yeast cells that undergo senescence significantly enhance protein disaggregation function by inducing Hsp104 disaggregase levels to counteract proteostasis defects^65^ suggesting that stress-induced protein disaggregation activity is wired differently in aging unicellular and multicellular organisms.

The implications related to the inactivation of this PQC machine during replicative aging could be significant. As we show, the assembly of this disaggregase already occurs at fever-like temperatures (Fig. 1d). In cells already expressing a high load of aggregation-prone proteins, this mild stress can trigger amyloidogenesis, as here exemplified in polyQ-expressing dermal fibroblasts (Fig. 4g). If Hsp70-DNAJA1-DNAJB1 disaggregase activity is reduced, as we see in cells undergoing replicative aging, these amyloids will not resolve and could lead to disease progression. This may also contribute to the faster decline rates observed for processes associated with aging in HD models and HD patients.^66^ Collectively, our data show how metazoa have evolved an intricate network of protein disaggregase machines that perform different sequential tasks, all required to ensure cell resilience to acute and chronic stresses, relevant to mammalian cellular aging and organismal health span.

## Methods

### Plasmids

Exon 1 carrying C-terminally tagged HttQ25-EmGFP and HttQ46-EmGFP were generated in the mammalian expression vector pTREX (Invitrogen). VCP (wt)-EGFP (Addgene, catalog number: 23971), pFRT/TO eGFP TIA1 (Addgene, catalog number: 106094) and pCMV3-GFP-hnRNPK (LSBio, catalog number: LS-N40093) were purchased. pcDNA5/FRT/TO HttQ119-eGFP was previously generated.^67^ pEGFP-N1 Hsp104 and pEGFP-N1 Hsp104 ^E285Q, E687Q^ were kind gifts from Bernd Bukau, Ph.D. (Heidelberg University). The generation of Hsp70 (HSPA8), Hsp110 (HSPH2), DNAJA1, DNAJA2, DNAJB1, DNAJB4, and mutant variants tagged with 6xHis-Smt3 in the vector pCA528 for recombinant expression and purification was previously described.^12,25^ Standard recombinant DNA techniques described by Sambrook et al.^68^ combined with standard site-directed mutagenesis with Phusion Plus DNA Polymerase (Thermo Fisher Scientific) were used to generate V5-DNAJA1, V5-DNAJA2, V5-DNAJB1, V5-DNAJB4, V5-DNAJB1^H32Q^, DNAJB1-HA, DNAJB1^D4R,E69R,E70R^-HA, DNAJB1^H32Q,D34N^-HA in pcDNA5/FRT/TO mammalian expression plasmid (Thermo Fisher Scientific).^69^ Gene sequences were verified by sanger sequencing.

### Protein purification

Human Hsp70, Hsp110, DNAJA1, DNAJA2, DNAJB1, DNAJB4, JDP mutants and firefly luciferase were recombinantly expressed and purified as previously described.^12,25^ Briefly, *E. coli* strains BL21 (D3E)/ pRARE carrying the corresponding expression vectors were induced for protein expression with 0.5 mM isopropyl-1-thio-D-galactopyranoside (IPTG, Sigma-Aldrich) for three hours at 30 ° C. Cells were lysed in 50 mM HEPES-KOH, pH 7.5, 750 mM KCl, 5 mM MgCl_2_, protease inhibitor cocktail (Roche), 2 mM phenylmethylsulphonyl fluoride, 10% glycerol (for human JDP purifications) or 50 mM HEPES-KOH, pH 7.5, 300 mM KCl, 5 mM MgCl_2_, protease inhibitor cocktail (Roche), 2 mM phenylmethylsulphonyl fluoride, 10% glycerol (for human Hsp70/110 purifications). After centrifugation at 30000 x g (30 min, 4 ° C) the resulting supernatants were applied to the Ni-NTA/Ni-IDA matrix and incubated for 60 min at 4 ° C. Subsequent washing steps were performed with high salt buffers (50 mM HEPES-KOH, pH 7.5, 750 and 500 mM KCl, 5 mM MgCl_2_ and 10% glycerol or 50 mM HEPES-KOH, pH 7.5, 300 and 100 mM KCl, 5 mM MgCl_2_ and 10% glycerol for human JDP and Hsp70/110 purifications, respectively). In the case of Hsp70/110 purification, an additional wash step was included with ATP buffer (50 mM HEPES-KOH, pH 7.5, 100 mM KCl, 5 mM MgCl_2_, 5 mM ATP). Protein elution was performed with 300 mM imidazole in the corresponding low salt buffers. Dialysis was performed overnight at 4 ° C in the presence of 4 µg of the 6xHis tagged Ulp1 per mg of substrate protein for proteolytic cleavage of the 6xHis-Smt3 tag. The 6xHis-Smt3 tag and Ulp1 were removed by incubating the dialyzed proteins in a Ni-NTA/ Ni-IDA matrix for 60 min at 4 ° C. Pyruvate kinase was purchased from Sigma-Aldrich. Proteins were further purified using Superdex 200 (GE Healthcare).

### Luciferase disaggregation assay

Luciferase disaggregation assays were performed as previously described.^14^ Briefly, 20 nM firefly luciferase was incubated at 45 ° C for 15 minutes in HKM buffer (50 mM Hepes-KOH pH 7.5, 50 mM KCl, 5 mM MgCl_2_, 2 mM DTT, 2 mM ATP pH 7.0, 10 μM BSA) to generate thermally denatured luciferase. Protein disaggregation was initiated by adding 1 µM HSPA8, 100 nM HSPH2, 250 nM JDP/mutant variants (total) and an ATP regenerating system (3 mM phosphoenolpyruvate (PEP) and 20 ng/μl pyruvate kinase) to the preformed luciferase aggregates followed by shifting the reaction to 30 ° C. 1 µM of recombinant yeast Hsp104 (kind gift from Yuki Oguchi, Ph.D.) was added to generate the bichaperone disaggregation system. Luciferase activity was measured at the indicated time points with a Lumat tube luminometer (Berthold Technologies) by transferring 1 µL of the sample to 100 µL of assay buffer (25 mM Glycylglycin, pH 7.4, 5 mM ATP pH 7, 100 mM KCl, 15 mM MgCl_2_) mixed with 100 µL of 0.25 mM luciferin. The Hsp70 inhibitor VER-155008 (Sigma, SML0271) was added at a working concentration of 5 µM.

### Cell culture

Human adenocarcinoma epithelial cell line HeLa (ATCC CRM-CCL-2) and human embryonic kidney HEK 293 cells (ATCC CRL-3216) were obtained from the American Type Culture Collection. HEK 293 *JB1* KO cells were kindly provided by Christian Hansen, Ph.D. (Lund University). These cell lines were cultured in DMEM (Gibco, Thermo Fisher Scientific, catalog number: 10569044) supplemented with 10% (v/v) heat inactivated fetal bovine serum (FBS: Gibco catalog number: 10082147), 1x L-alanine-L-glutamine dipeptide (Gibco Glutamax 100x catalog number: 35050061), 10 mM sodium pyruvate (Gibco catalog number: 11360070) and 100 U/ml penicillin/streptomycin (Gibco 1x 104 U/ml catalog number: 15140122) at 37 ° C in 5% CO_2_. Human dermal fibroblasts were isolated and grown from 6 mm skin punch biopsies as previously described.^70^ Dermal fibroblasts were cultured on 0.1% gelatin (Sigma catalog number: G1890) treated wells using fibroblast media consisting of DMEM (Gibco catalog number: 10569044), 10% FBS, 0.1 mM MEM non-essential amino acids solution (Gibco catalog number: 11140050), 100 U/ml penicillin/streptomycin, 0.1mM 2-Mercaptoethanol, (Gibco catalog number: 21985023). Cell passaging: 0.25 million dermal fibroblasts were seeded in a 10 cm culture plate until cell confluency reached 70%. This was defined as a passage (P), and the process was repeated to generate replicatively aged cell lines. Mycoplasma contamination of cells was tested and ruled out with the LookOut Mycoplasma Detection kit (Sigma-Aldrich; catalog number: MP0035).

### Heat shock and drug treatments

Heat shock was performed by exposing HeLa cells to 43 ° C for 1 hour in a water bath or otherwise indicated. A 42 ° C, 2 hours heat shock in an incubator was applied to HEK 293 and human dermal fibroblasts. Following heat shock, cells were returned to 37 ° C for stress recovery. Temperature measurements for heat shock conditions in the water bath and incubator were determined using a VOLTCRAFT K101 digital thermometer (Conrad). Hsp70 inhibitor VER-155008 (Sigma, catalog number: SML0271) was used at a working concentration of 5 µM (*in vitro*) or 50 µM (in cells) or otherwise stated. Protein synthesis was inhibited by adding cycloheximide (CHX) (Sigma, catalog number: 01810) at a final concentration of 100 µg/mL. NMS-873 (Sigma, catalog number: SML 1182) was used for VCP inhibition at a final concentration of 1 or 10 µM.

### *In situ* proximity ligation assay (PLA)

PLA was performed as previously described.^14,26^. Cells were washed 2x with PBS, fixed with 4% paraformaldehyde in PBS, and the proximity ligation assay was performed according to the DUOLINK manufacturer’s guideline for mammalian cells (Sigma-Aldrich). DNA was stained with DAPI-containing mounting medium (Sigma-Aldrich, catalog number: DUO82040). The staining, imaging, processing, and analysis conditions were kept consistent for each experiment.

### Immunocytochemistry and staining

Cells were cultured on soda lime glass diagnostic slides masked with epoxy resin (Marienfield-superior, catalog number: 1216530) until 70-80% confluence. Cells were washed 2x with PBS, fixed with 4% paraformaldehyde in PBS for 10 minutes, and washed 2x again with PBS. Cells were permeabilized with 0.2% Triton X-100 in PBS for 10 minutes. After permeabilization, cells were washed 2x with PBS and blocked for 10 minutes in PBS containing 0.5% bovine serum albumin and 0.15% Glycine (PBS+ buffer). Primary antibodies were diluted in PBS+ buffer and incubated for 1.5 hours at room temperature (RT) or overnight at 4 ° C. After incubation, cells were washed 4x with PBS+ buffer. Secondary antibodies were diluted in PBS+ buffer and incubated for 1.5 hours at RT, followed by 2x washes in the same buffer. DNA was stained with DAPI-containing mounting medium (Sigma-Aldrich, catalog number: DUO82040) or Hoechst staining (Thermo Scientific, catalog number: 62249) according to the manufacturer’s guidelines.

### Transfections and siRNA knockdowns

Plasmid transfections were performed according to standard protocols using Lipofectamine 2000 or 3000 (Thermo Fisher Scientific). siRNA transfections were carried out according to standard protocols using DharmaFECT one transfection reagent (Dharmacon, DHA-T-2001). HeLa cells were transfected with 25-50 nM of Dharmacon onTarget Plus smartpool siRNA against DNAJA1 (Dharmacon, L-019617-00), DNAJA2 (Dharmacon, L-012104–01), DNAJB1 (Dharmacon, L-012735–01), DNAJB4 (Dharmacon, L-015678-01), HSF-1 (Dharmacon, L-012109-02), BAG1 (Dharmacon, L-003871-00), HSPBP1 (Dharmacon, L-016607-00), HSPH1 (Dharmacon, L-004972-00), HSPA4 (Dharmacon, L-012636-00), HSPA4L (Dharmacon, L-017633-00), VCP (Dharmacon, L-008727-00), or scrambled non-targeting siRNA (Dharmacon, D-001810–10) for 48 hours. The knockdown efficiencies were verified by immunoblotting using specific antibodies.

### Confocal microscopy

LSM 780 and LSM 980 laser scanning confocal systems on inverted stands with diode 405 nm, argon 488 nm, and DPSS 561 nm lasers (Carl Zeiss, Germany) were used for microscopy. The images were taken with the Plan-Apochromat 40x/1.4 oil objective (Carl Zeiss) and identical acquisition settings with a pinhole of approximately 1 AU. Maximum projection images were generated from acquired z-stacks. Linear adjustments of brightness and contrast were performed to the same degree within each experiment. The DAPI, PLA, and GFP signals were pseudo-colored in magenta, yellow, and cyan. Fluorescence intensity profile plots (line scans) were generated on the raw images using the RGB Profiler plugin in Fiji.

### PLA quantification

The PLA signal was quantified using custom code (https://github.com/Sanudish/Punctate-Detection.git) in MATLAB R2021a using functions from the Image Processing Toolbox, and Statistics and Machine Learning Toolbox. Briefly, the brightest slice of each volume, determined by the cumulative intensity of the slice, was extracted to perform the quantification. The nuclei-stained channel of the images was first processed with a median filter followed by a maximum filter, both of which utilized a 4×4 kernel. The user input-based percentile thresholding was then applied to the processed images. The connected nuclei were then further segmented through the implementation of a Watershed transformation. To refine nuclei segmentation, user inputs for the degree of connectivity and area were utilized. The output enabled quantification of the number of cells per image. Next, for extraction of puncta, logarithmic frequency histograms of the intensity profiles were plotted to generate a normalized user input range between −1 and 1 to threshold out puncta. To improve the segmentation of the puncta from noise, area thresholding was implemented. The resulting number of puncta, and their characteristics were measured. The product of the average punctate mean intensity per cell and the number of punctate structures per cell was calculated to obtain the PLA raw parameter. The final induction of Hsp70 and JDP assemblies were reported as a PLA fold change to an unstressed condition.

### Colocalization between PLA signal and Hsp104^DWB^-GFP positive heat-induced foci

Hela cells expressing Hsp104^DWB^-GFP were heat shocked at 42 ° C for 2h and allowed to recovery at 37 ° C for 12h. Cells were treated with cycloheximide (CHX) at 3.5h and VER-155008 at 5.5h after HS. Super-resolution images were acquired using LSM 980 Airyscan super-resolution module (Carl Zeiss, Germany). Raw images were processed using the Zen Blue software’s Airyscan Processing function in 3D mode. Colocalization analysis between PLA signal and Hsp104^DWB^-GFP foci was conducted using Fiji plugin Just Another Colocalization Plugin (JACoP).^71^ Manders’ overlap correlation coefficients were calculated using thresholded images (n=5).

### Aggregate quantification

Endogenous protein aggregates and TIA-1 biomolecular condensates in HeLa cells were quantified using a Fiji macro termed AggreCount as previously described.^72^ Briefly, the aggregates were extracted by processing images with gamma enhancement, mean background reduction, and a user-based threshold. The contrast enhancement of the nuclei enabled segmentation of nuclei regions, which were then utilized to generate a composite image with the preprocessed image. The signal-enhanced composite image enables the detection of local maxima of cell regions. Implementing the watershed algorithm, a Voronoi diagram was generated and the proportion of cells with aggregates was measured. The fold change was determined by calculating the difference of each parameter in the stress recovery conditions and the unstressed condition. When segmenting hnRNPK aggregates, the threshold diameter of inclusions was set at 1 µm to avoid intranuclear inhomogeneities being falsely detected as aggregates. The aggregates in HEK 293 cells were calculated manually as previously described^73^ because AggreCount segmentation was less efficient.

### Fluorescence Recovery After Photobleaching

HeLa cells expressing GFP tagged hnRNPK were exposed to 43 ° C for 1 hour in a water bath and returned to 37 ° C for 1 hour. FRAP was performed on heat-induced hnRNPK aggregates using an LSM 780 laser scanning confocal system on inverted stands at 37 ° C with argon 488 nm laser (Carl Zeiss, Germany). The images were taken with the Plan-Apochromat 40x/1.4 oil objective (Carl Zeiss). Aggregates were photobleached using 100% FRAP laser power for 10 ms. Images were obtained every 500 ms to observe fluorescence recovery up to 40 s. Images were analyzed using easyFRAP.^74^ Recovery curves were normalized to background fluorescence for subtracting noise, and adjacent non-bleached aggregates for fluorescence intensity fluctuations. Average and standard errors were calculated from seven individual traces.

### Immunoblotting

Immunoblot analysis was performed using standard methodologies.^14^ Briefly, cells were lysed in RIPA buffer with a protease inhibitor cocktail (Roche cOmplete - EDTA free, catalog number: 05056489001) and sonicated (3x cycles, each cycle at 5 seconds and 50% amplitude) in a Branson SFX150 Digital Sonifier (Emerson). Protein concentrations were determined using the bicinchoninic acid protein assay kit (PierceTM, BCA, catalog numbers: Reagent-A: 23228, Reagent-B: 23224). Protein samples were stabilized with 1x sample buffer and boiled for 5 minutes at 95 ° C. For SDS-PAGE, 25 µg total protein was loaded into 4% stacking gel and resolved through 12% running gel. Protein transfer to a PVDF membrane was performed using a Bio-Rad transfer tank (Mini Trans-Blot Cell). Anti-rabbit (800CW; C90220-05) and anti-mouse (680RD; C90219-05) antibodies from LI-COR Biosciences were used as secondary antibodies. The detection was carried out with an Odyssey Immunoblotting Imaging System (LI-COR).

### Supernatant-pellet assay

HeLa cells were grown in 100 mm cell culture plates. Before lysis, cells were washed 1x with ice-cold PBS, scraped, and collected with 500 µl of ice-cold PBS. Cells were collected at 600 x g for 5 minutes at 4 ° C. Cell pellets were resuspended in lysis buffer containing 25 mM HEPES, 75 mM NaCl, 1 mM MgCl_2_, 1% NP-40, 75 U/ml Benzonase (EMD Millipore) and Protease Inhibitors cocktail (Roche) and lysed by vortexing for 20 seconds and subsequently for 5 seconds every 10 minutes over the course of 1 hour. Protein quantification was performed using the DC-protein assay (Bio-Rad) and the samples were equalized. 10% of the total protein lysate was transferred to a fresh Eppendorf tube and boiled in Laemmli buffer (60 mM Tris pH 6.8, 2% SDS, 10% glycerol, 5% β-mercaptoethanol). From the remaining lysate, the pellet fractions were isolated by centrifuging the samples at 20,817 x g for 30 minutes at 4 ° C and removing the supernatant. The pellet fractions were washed and resuspended in urea buffer (8M Urea, 50 mM Tris pH 7.4, 2% SDS, 50 mM DTT). The resuspended samples were shaken at 1000 rpm in a heating block at 60 ° C for 1 hour and subsequently at 1000 rpm in RT, overnight, and boiled prior to SDS-PAGE.

### qRT-PCR

HeLa cells were washed with ice cold PBS, scraped in TRIzol reagent (Invitrogen) and incubated at RT for 5 minutes. Chloroform was added to cell lysates, vortexed for 15 seconds and incubated for 2-3 minutes before centrifugation at 18,000 x g for 15 minutes at 4 ° C. The aqueous phase was transferred to a tube without DNAase/RNAase, isopropanol was added and incubated in RT for 10 minutes. RNA was precipitated by centrifugation at 18,000 x g for 10 minutes at 4 ° C. The supernatants were removed, 75% ethanol was added to the pellets, and the samples were centrifugated at 18,000 x g for 5 minutes at 4 °C. The supernatants were removed and the pellets were resuspended in DNA/RNAase-free H_2_O. DNAases were removed with the Turbo DNAase free kit (Invitrogen) according to the manufacturer’s instructions. cDNA synthesis was performed using M-MLV reverse transcriptase (Invitrogen) according to the manufacturer’s instructions. The primers used in the PCR step were previously described.^75^

### LDH-Glo cytotoxicity assay

Heat shock-induced cytotoxicity was determined using the LDH-Glo cytotoxicity assay (Promega, catalog number: J2380) according to the manufacturer’s guidelines. Briefly, HeLa cells and human dermal fibroblasts were cultured in a 6-well plate at 37 ° C. Heat shock was performed at 43 ° C in a water bath for 1 hour and returned to the 37 ° C incubator for recovery. VER-155008 (5 µM) and cycloheximide (100 µg/mL) treatments were performed 2.5 hours after heat shock. 5 µL samples from the cell culture media were collected at indicated time points during heat shock recovery, and immediately mixed with 95 µL of LDH storage buffer and kept at 4 ° C. The LDH-Glo cytotoxicity assay was performed using a plate reader (CLARIOstar^Plus^, BMG Labtech).

### Cellular senescence detection assay

Cellular senescence was assessed using fluorescence- and immunocytochemistry-based assays. The fluorescence-based assay was performed with the Cellular Senescence Plate Assay Kit-SPiDER-βGal (Dogindo, catalog number: 05-05) according to the manufacturer’s guidelines. Fluorescence intensity was measured using a plate reader (CLARIOstar^Plus^, BMG Labtech; Excitation: 535 nm/ emission: 580 nm). SA-β-Gal activity was normalized to cell count determined using a hemocytometer. The immunocytochemistry-based assay was performed with Senescence β-Galactosidase Staining Kit (Cell Signaling Technology, catalog number: 9860) according to the manufacturer’s guidelines. DIC images were obtained on a Leica DMi8 system using Plan-Apochromat 40x/1.4 oil objective. The percentage of cells positive with β-Galactosidase signal was quantified manually.

### Proteasome activity assay

The chymotrypsin-like activity of the 20S proteasome was assessed using a commercially available fluorometric proteasome activity assay kit (Abcam, catalog number: ab107921) according to the manufacturer’s guidelines. Fluorescence intensity was measured using a plate reader (CLARIOstar^Plus^, BMG Labtech; Excitation: 350 nm/ emission: 440 nm). The readout was normalized to the protein concentrations of the lysate samples determined using the bicinchoninic acid protein assay kit (PierceTM, BCA, catalog numbers: Reagent-A: 23228, Reagent-B: 23224).

### Multiple reaction monitoring mass spectrometry

HeLa cells were cultured in 150 mm cell culture plates until 70-80% confluence. Light samples were grown as previously described. To generate a heavy internal standard, cells were grown in arginine and lysine-depleted DMEM (Thermo Fisher Scientific) supplemented with 100 mg/L L-arginine-HCl, ^13^C_6_, ^15^N_4_ (Thermo Fisher Scientific) and 100 mg/L L-Lysine-2HCl ^13^C_6_, ^15^N_2_ (Thermo Fisher Scientific). The cells were grown in the heavy medium for 6 passages until the incorporation of heavy amino acids was at least 95%. To ensure that all proteins of interest were detectable, a heavy standard was generated from lysates of unstressed and heat shocked cells for 1 hour at 43 ° C. Before lysis, cells were washed with ice-cold PBS and detached by scraping with 1 ml of ice-cold PBS. Cells were pelleted at 500 x g for 5 minutes at 4 ° C and the supernatant was discarded. Cell pellets were snap frozen in liquid nitrogen and stored at −80 ° C or processed immediately. Cell pellets were resuspended in Benzonase containing urea buffer (6 M Urea, 50 mM Tris pH = 7.5, 2 mM MgCl_2_, 50 mM DTT, 0.25 U µL^−1^ Benzonase) and lysed for 10 minutes at RT. The lysate was centrifuged at 5,000 x g for 10 minutes in RT. The supernatant was transferred to a fresh tube. Protein quantification was performed using the DC Protein Assay (Bio-Rad).

Light samples and heavy internal standards were mixed in a 1:1 (w/w) ratio and 100 µg of total protein was precipitated using a mixture of methanol and chloroform according to a published protocol ^76^. For in-solution digestion, precipitated protein pellets were resolved in 50 µL of urea buffer (8 M Urea, Carl Roth, 100 mM NaCl, Applichem, in 50 mM triethylammonium bicarbonate (TEAB), pH 8.5, Sigma-Aldrich). To reduce disulfide bridges, the sample was incubated with 10 mM 1,4-dithiothreitol (Sigma-Aldrich) for 30 min at RT, constantly agitated at 350 rpm. Next, alkylation was performed with 30 mM iodacetamide (Sigma-Aldrich) in 100 mM TEAB buffer for 30 min at RT in the dark. Sample predigestion was performed by adding 2.5 µg Lysyl Endopeptidase® (Wako Chemicals) and incubating for 4 hours at 28 °C and 350 rpm agitation. After diluting the urea concentration to 2 M by adding 50 mM TEAB buffer, 1 µg of trypsin (Thermo Fisher Scientific) was added and the samples were incubated for 16 hours at 28 °C and 350 rpm agitation. The digestion process was stopped by reducing pH < 2 by adding trifluoroacetic acid (TFA, Sigma-Aldrich) to a final concentration of 0.4 – 0.8% (v/v). The samples were centrifuged for 10 min at 20,000 x g in RT and the supernatant was desalted using SepPak C18 cartridges (500 mg; Waters) and the Waters Extraction Manifold. The SepPak C18 cartridges were equilibrated by adding 3 ml of 100% acetonitrile (ACN), then 3 ml of 50% ACN / 0.1% TFA and finally 3 ml of 0.1% TFA. Digested peptide samples were loaded onto SepPak cartridges and washed with 3 ml of 0.1% TFA. Finally, the samples were eluted into fresh LoBind tubes using 1 ml of 50% ACN / 0.1% TFA. The samples were dried in a vacuum centrifuge and stored at −20 ° C until LC-MRM analysis. Before LC-MRM analysis, the peptides were resolubilized in 10 µL of 20% ACN / 0.1% (v/v) TFA and incubated for 5 min in RT, then diluted 10-fold with 0.1% TFA to obtain a final concentration of 2% ACN. 5 μg of tryptic digest including a set of 11 synthetic peptides for indexed retention time scheduling^77^ were analyzed per LC-MRM injection. The samples were measured in a randomized order using a nanoACQUITY UPLC System (Waters) online-coupled to an ESI-QTrap 5500 via a NanoSpray III Source (both Sciex). The UPLC was equipped with a trapping column (Waters, Symmetry C18; 2 cm length, 180 μm inner diameter, 5 μm C18 particle size, 100 Å pore size) and an analytical column (Waters, M-Class Peptide BEH C18; 25 cm length, 75 μm inner diameter, 1.7 μm C18 particle size, 130 Å pore size). The peptides were trapped for 7 min at a flow rate of 10 µL min^−1^ with 99.4% of buffer A (1% v/v ACN, 0.1% v/v FA) and 0.6% of buffer B (89.9% v/v ACN, 0.1% v/v FA) and separated at the analytical column temperature of 60°C, with a flow rate of 300 nl min^−1^ and a 37 min linear gradient of buffer B in buffer A from 3 to 37% B, followed by washing and reconditioning of the column to 3% B. Uncoated precut emitters (Silica Tip, 20 μm inner diameter, 10 μm tip inner diameter, New Objectives, Woburn, MA) were used and a voltage of approximately 2.6 kV was applied for electrospray ionization. QTrap 5500 was operated in scheduled mode at Unit resolution for Q1 and Q3 with 180 sec MRM detection window width and 2.5 sec target scan time. Peptide retention times for scheduling were previously determined using commercial synthetic peptide standards (JPT).

### LC-MRM data and statistical analysis

The MRM data was first manually reviewed using Skyline 64-bit version 20.1. Data sets were imported into Skyline, and signal quality was assessed using the Peak Areas and Retention Time visualization features. We ensured via the Results Grid that all reference dot-product (rdotp) correlations for ratios of the observed transition intensities between the corresponding light and heavy precursor pairs were equal to or above 0.9. Then, all data were exported from Skyline and imported into R version 4.0.3 (2020-10-10) “Bunny-Wunnies Freak Out” for further analysis using a custom script. Briefly, all quantitative transitions were summed up across proteins, and log2 light/heavy ratios were computed to account for technical variance. In addition, all samples were normalized to their respective tubulin alpha-1A protein level to account for biological variance. Then, pairwise group comparisons were performed using linear regression models between protein ratios of different conditions and the unstressed control. The slope of the linear regression corresponded to the fold change between the respective condition and the control and was plotted together with its respective standard error.

### Identification of aggregated proteins recovered by Hsp70 disaggregase activity (post 6h heat shock)

Briefly, HeLa cells were heat-shocked followed by sustained recovery over 12h. We chose recovery time points that encompass the evolution and resolution (i.e. 6h and 12h post-HS) of aggregates that appeared to be the main target of Hsp70-DNAJA1-DNAJB1. To quantify cellular proteins that change solubility at different harvest time points, we used isobaric labeling (Tandem mass tag; TMT), a method that allows for multiplexing of samples collected at different time points in a recovery course into a single experiment (Supplementary Data Fig. 12a). The gist of TMT is that peptides are labeled with one of several distinct agents of the same mass, but cleave into distinct reporter ions during mass spectrometry fragmentation. The workflow included a parallel examination of stress recovery in which Hsp70-DNAJA1-DNAJB1 disaggregase activity was inhibited using VER-155008 to identify potential substrates. The proportion of proteins in the supernatant compared to the total proteome (pSup) was normalized to solubility prior to HS (ΔpSup), and then the effect of VER-155008 was compared to the DMSO control (ΔΔpSup). As expected, we observed a significant decrease in the median pSup at 6h after HS in both DMSO and VER-155008 treated samples (Supplementary Data Fig. 12b; p value < 0.0001, paired Wilcoxon rank test) that supports aggregation throughout the proteome as a result of heat denaturation. After 12 hours of recovery, more than 75% of proteins had recovered their solubility. However, as expected, a subset of proteins recovered differently in the presence of VER-155008, as measured by ΔΔpSup. Proteins with ΔΔpSup > +/ 0.26 (determined by Gaussian thresholding at the 99^th^ percentile of pseudo pre-HS ΔpSup) were considered to have significantly altered solubility due to the presence of VER-155008. Of these proteins, those that were less soluble in the presence of VER-155008 compared to DMSO (ΔΔpSup > 0) at at least one time point were identified as potential clients solubilized by the disaggregase machinery (a total of 51 proteins, Supplementary Data Table 1). We then sought to describe the general behavior of ΔΔpSup among these proteins using unsupervised k-means clustering. The proteins of interest were sorted into four distinct patterns (Supplementary Data Fig. 12c); (i) those that were more soluble at 6h-HS in the presence of VER-155008 compared to the DMSO control, but whose recovery was inhibited compared to the DMSO control at the 12h timepoint, (ii) those that were less soluble at 6h post-HS in the presence of VER155008 but then recovered similarly to the DMSO control at 12h, (iii) those with no difference in solubility due to VER-155008 at 6h post-HS, but whose recovery was inhibited compared to the DMSO control at the 12h timepoint, and (iv) those that were less soluble at both 6h and 12h post-HS in the presence of VER-155008 (Supplementary Data Table 1). Protein degradation and processes such as the recently discovered ubiquitinome reshaping during stress recovery could have in part influenced the generated patterns we observed.^61^ Given that the addition of VER-155008 was at 5h post-HS, we postulated that its impact on disaggregation would be most visible at the 12h timepoint. Therefore, we considered proteins with minimal differences in solubility at 6h but inhibited recovery at 12h in the presence of VER-155008 (cluster iii) for validation.

#### Cell fractionation by ultracentrifugation

HeLa cells were grown in T75 cell culture flasks (Nunc EasYFlask, catalog number: nun156472). Before lysis, cells were washed with 1x ice-cold PBS, scraped and collected with 10 mL of ice-cold PBS. The cells were then pelleted by a mild centrifugation step at 120 x g for 6 minutes at 4 ° C twice and the supernatant was discarded. Cell pellets were resuspended in 500 µL of lysis buffer 1 containing 50 mM Tris-HCl pH 7.5, 150 mM NaCl, 1% (v/v) IGEPAL CA-630, 10 units/mL DNase I, 1:1,000 protease inhibitor, and Protease Inhibitors cocktail (Roche, catalog number: 11836145001) and lysed by extrusion through a 27-gauge needle for 25 times, followed by a 31-gauge needle for 10 times. Subsequently, EDTA was added to a final concentration of 2 mM. Cell lysate was then divided into two, one for the assessment of total protein and the other pelleted at 100,000 x *g* for 20 min at 4 ° C. The resulting supernatant was the collected sample. The volume of total protein and supernatant samples was measured and resuspended in an equal volume of a lysis buffer 2 containing lysis buffer 1, 4% sodium dodecyl sulfate (SDS, Thermo Scientific, catalog number: 28365) and 4 mM 1,4-dithiothreitol (DTT, Thermo Scientific, catalog number: R0861). Subsequently, the samples were incubated at 95 ° C for 20 minutes.

#### NanoESI-LC-MS/MS

Samples were prepared for mass spectrometry as previously detailed.^19^ Freeze-dried peptides were resuspended in 80 µl of distilled water and quantified by the BCA assay (Thermo Fisher Scientific) according to the manufacturer’s instruction. 20 µl of each sample was diluted into a final volume of 40 µl containing 100 mM TEAB and 30% (v/v) acetonitrile. The samples were then labeled with TMT 11-plex (Thermo Scientific, catalog number: A37725) according to the manufacturer’s protocol. The samples were analyzed by nanoESI-LC-MS/MS using an Orbitrap Exploris mass spectrometer (Thermo Scientific) equipped with a nanoflow reversed-phase-HPLC (Ultimate 3000 RSLC, Dionex) fitted with an Acclaim Pepmap nano-trap column (Dionex—C18, 100 Å, 75 µm × 2 cm) and an Acclaim Pepmap RSLC analytical column (Dionex—C18, 100 Å, 75 µm × 50 cm). For each iteration of LC-MS/MS, 0.6 µg of the peptide mix was loaded onto the enrichment (trap) column at an isocratic flow of 5 µl min^−1^ of 2% ACN containing 0.1% (v/v) trifluoroacetic acid for 5 min before the enrichment column was switched in-line with the analytical column. The eluents were 0.1% (v/v) formic acid/5% DMSO (v/v) (solvent A) and 100% ACN/0.1% formic acid/5% DMSO (v/v) (v/v) (solvent B), flowed at 300 nL min^−1^ using a gradient of 3–22% B in 90 min, 22–40% B in 10 min and 40–80% B in 5 min then maintained for 5 min before re-equilibration for 8 min at 3% B prior to the next analysis. All spectra were acquired in positive ionization mode with full scan MS acquired from m/z 300-1600 in the FT mode at a mass resolving power of 120 000, after accumulating to an AGC target value of 3.00e^6^, with a maximum accumulation time of 25 ms. The RunStart EASY-IC lock internal lockmass was used. Data-dependent HCD MS/MS of charge states > 1 was performed using a 3 s scan method, at a normalized AGC target of 75%, automatic injection, a normalized collision energy of 30%, and with spectra acquired at a resolving power of 15000. Dynamic exclusion was used for 20s.

#### Peptide identification

The initial analysis of raw mass spectra was carried out using MaxQuant (v 1.6.14.0) against the Swissprot Homo Sapiens database (downloaded 04/03/2020; containing 20350 entries). Searches were conducted with 20 ppm mass tolerance for MS, and 0.2 Da for MS/MS, with one missed cleavage allowed and match between runs enabled. The MS2 reporter ion was set to TMT 11-plex and isotopic distribution correction was applied according to the product data sheet. Variable modifications included methionine oxidation and N-terminal protein acetylation, while the carbamidomethylcysteine modification was fixed. The false discovery rate maximum was set to 0.005% at the peptide identification level (actual was 0.005 for each replicate) and 1% at the protein identification level. All other parameters were left as default.

#### Ratio correction and scaling

Further analysis was performed with custom Python scripts. The logic was as follows. First, common contaminant proteins were removed, and quantified proteins were considered as those identified by at least one unique peptide. The ratio of abundance for each protein in the supernatant as compared to total samples (pSup) was corrected for differences in total protein amount as quantified via BCA assay described previously.^19^ The relative change in solubility for individual proteins after heat shock (ΔpSup) was then determined by normalizing to the preheat shock value, such that a protein with no change in solubility due to heat shock would have a value of 1. The resulting data were then scaled using a p-value weighted correction.^78^ This correction weights the mean ratio of biological replicates (*n=4*) according to the relative confidence with which it deviates from the expected value (in this case, 1) as per equation 1:

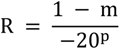

where *m* corresponds to the mean of the ΔpSup ratios and *p* corresponds to the p-value derived from a one-sample t-test of the ratios against the expected value under a single timepoint and treatment regime. Finally, the difference in relative solubility at 6 and 12h post-heat shock between the DMSO and VER-155008 treated samples (ΔΔpSup) was calculated as ΔpSup_DMSO_ – ΔpSup_VER_, such that no difference in relative solubility between the DMSO and VER-155008 treatments would yield a ratio of 0.

#### Identifying proteins of interest

To define a ΔΔpSup at which the changes in solubility could be considered biologically relevant, we first generated a pseudo-control ΔΔpSup by subtracting per-protein randomized pre-heat shock ratios (ΔpSup_R-Control_) from the observed pre-heat shock ratios (ΔpSup_Control_). The resulting frequency histogram of pseudo-control ΔΔpSup values was then fit to a Gaussian distribution, and by calculating three times the derived variance parameter (σ) we determined the interval encompassing 99.7% of the pseudo-control ΔΔpSup values. The boundaries of this interval (in this case ± 0.26, consistent with previous estimates of ± 0.3 applied to similar data sets^19^ were then used to threshold the ΔΔpSup values at each time point. Proteins that exceeded the ΔΔpSup threshold at least one time point were collected for additional analyzes. Functional overrepresentation was tested using PantherGOSlim (http://pantherdb.org), with a Bonferroni-corrected significance cutoff of p < 0.05 against the complete list of proteins identified. Annotated protein-protein interactions were collected from STRINGdb^79^ via Cytoscape (v 3.7.2).^80^ Finally, ΔΔpSup traces over time were clustered using k-means as implemented via the scikit-learn package.^81^

## Statistical Methods

All quantified experiments were performed at least in triplicate. Data are shown as mean ± s.e.m. and error bars indicating s.e.m. All statistical analyzes not described elsewhere were performed on RStudio (Version 1.1.447) or GraphPad Prism 8. One-way between groups analysis of variance (ANOVA) was conducted for multiple group comparisons. All analyses were considered statistically significant at p<0.05.

## Ethics

Derivation of human dermal fibroblasts from skin punch biopsies was performed with approval from the Monash University and Monash Medical Centre Human Ethics Committees (approval number 10006A).

## Data availability and code availability

The mass spectrometry proteomics data have been deposited with the ProteomeXchange Consortium through the PRIDE^82^ partner repository with the data set identifier PXD029161. All analysis scripts are available on request. The multiple reaction monitoring data will be deposited in the <u>ProteomeXchange</u> Consortium through the <u>Panorama</u> Public^83^ partner repository upon manuscript acceptance and can be reviewed at https://panoramaweb.org/siDAP.url

## Acknowledgements

We thank Paolo D. L. Rios, Ph.D., and Joachim Berger, Ph.D., and others for scientific discussions and critical reading of the manuscript. Special acknowledgements go to Oleksandr Chernyavskiy, Ph.D. (Monash Micro Imaging Facility) and Tim E. Powers, Ph.D. (Monash eResearch Center) for assistance with microscopy and statistical analysis. We thank Xinran Cheng, Ph.D., Peck S. Tan, Ph.D., Sophie Kopetschke, and Jeanette F. Brunsting for experimental/technical support. Some figures were created with BioRender.com.

## Funding

National Health and Medical Research Council of Australia Investigator Grant APP1197021 to NBN, APP1161803, and APP1154352 to DMH.

Recruitment Grant from Monash University Faculty of Medicine Nursing and Health Sciences with funding from the State Government of Victoria and the Australian Government to NBN. Yulgilbar Foundation, Australia to NBN

Australian Research Council of Australia Grant DP170103093 to DMH.

Deutsche Forschungsgemeinschaft Project ID FOR2509 P05 to TR.

Bundesministerium für Bildung und Forschung (BMBF), Ministerium für Kultur und Wissenschaft des Landes Nordrhein-Westfalen (MKW) and Regierender Bürgermeister von Berlin to RS.

We thank Merck PTY LTD Australia for its concessional product pricing for the work.

## Authors’ contributions

Conceptualization: NBN

Methodology: YM, NA, DC, CJR, RS, KMA and NBN

Software development: SU, YM and SA

Investigation: YM, NA, CJR, DC, NLP, ARO, RS, KMA, and NBN

Formal analysis: YM, NA, CJR, DC, RS, KMA, SU, TR, SA, DMH, HHK and NBN

Original draft writing: NBN

Editing: YM, NA, CJR, DC, RS, KMA, SU, TR, SA, DMH, HHK and NBN

Reviewing: YM, NA, DMH, HHK and NBN

## Competing interests

The authors declare no competing interests.

## Materials and correspondence

Correspondence and requests for materials should be addressed to

## Supplementary Materials

Supplementary Information is available for this paper.

## Rights and permissions

Reprints and permissions information is available at www.nature.com/reprints

## Supplementary Data Figure legends

**Supplementary Data Figure 1.**
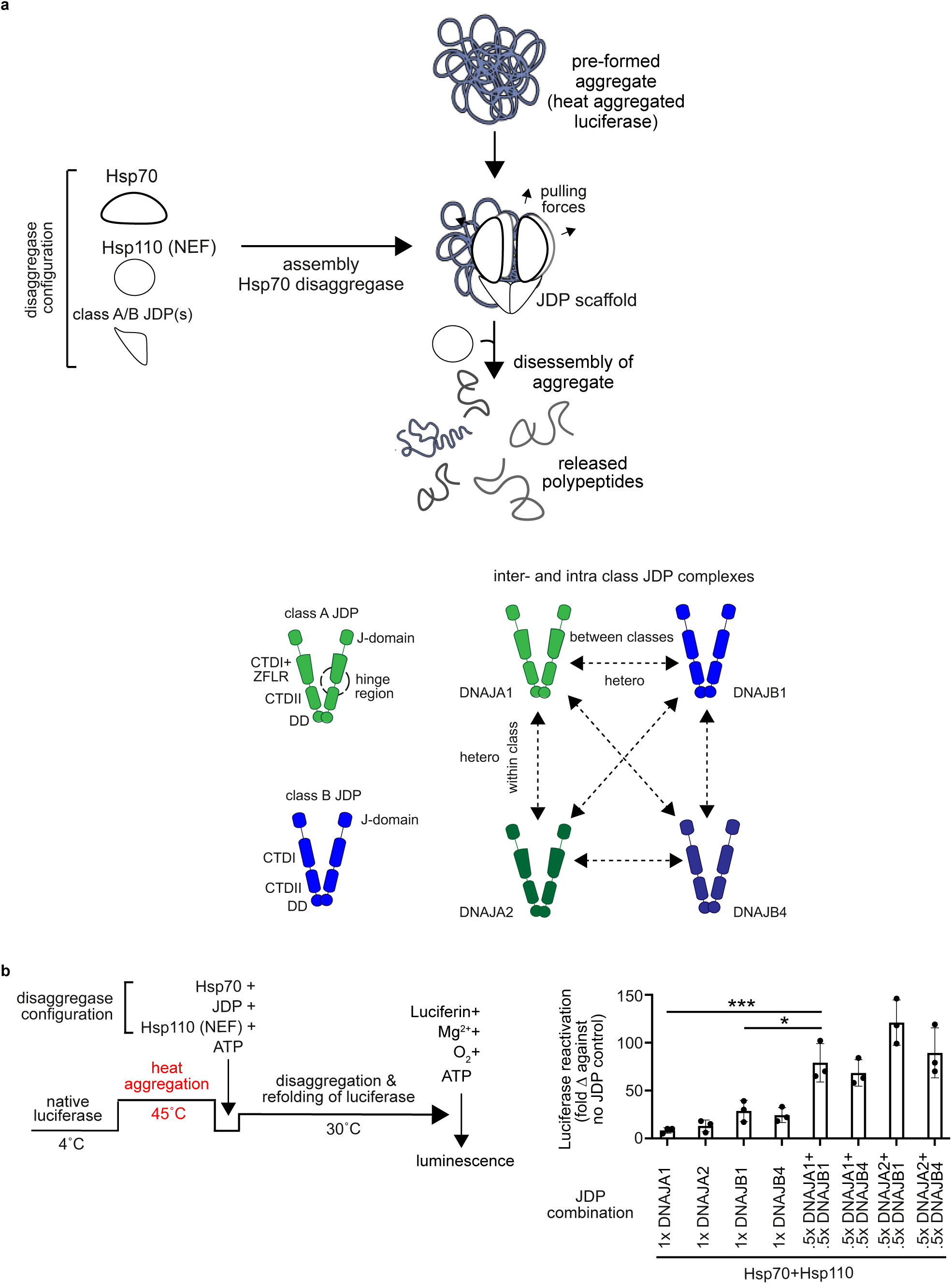
Reactions containing interclass JDP scaffolds show increased protein disaggregation/refolding activity, *in vitro*. **a**, Top: Schematic diagram of Hsp70, JDP and Hsp110 mediated protein disaggregation. JDP scaffolds presumably assemble on aggregates and recruit Hsp70. Hsp110’s primarily function as nucleotide exchange factors (NEFs) for Hsp70. Bottom left: Domain architectures of class A (green) and class B (blue) JDPs (depicted as dimers). JD, J-domain; CTD, C-terminal domain; ZFLR, Zink finger like region; DD, dimerization domain. Bottom right: Intra and interclass JDP complexation which results in the formation of homo or hetero JDP scaffolds (minimum is a dimer of a dimer) is driven by intermolecular interactions between JD and CTD.^12^ **b**, Left: Luciferase disaggregation/refolding assay. Heat denaturation triggers misfolding and aggregation of luciferase. Right: Disaggregation/refolding efficiencies by different JDP scaffolds containing Hsp70 disaggregases, *in vitro* (n = 3, data are mean +/− s.e.m., * adjusted p-value < 0.05, ** adjusted p-value < 0.01, one-way ANOVA Tukey post hoc test). Hsp110 functions as a NEF.

**Supplementary Data Figure 2.**
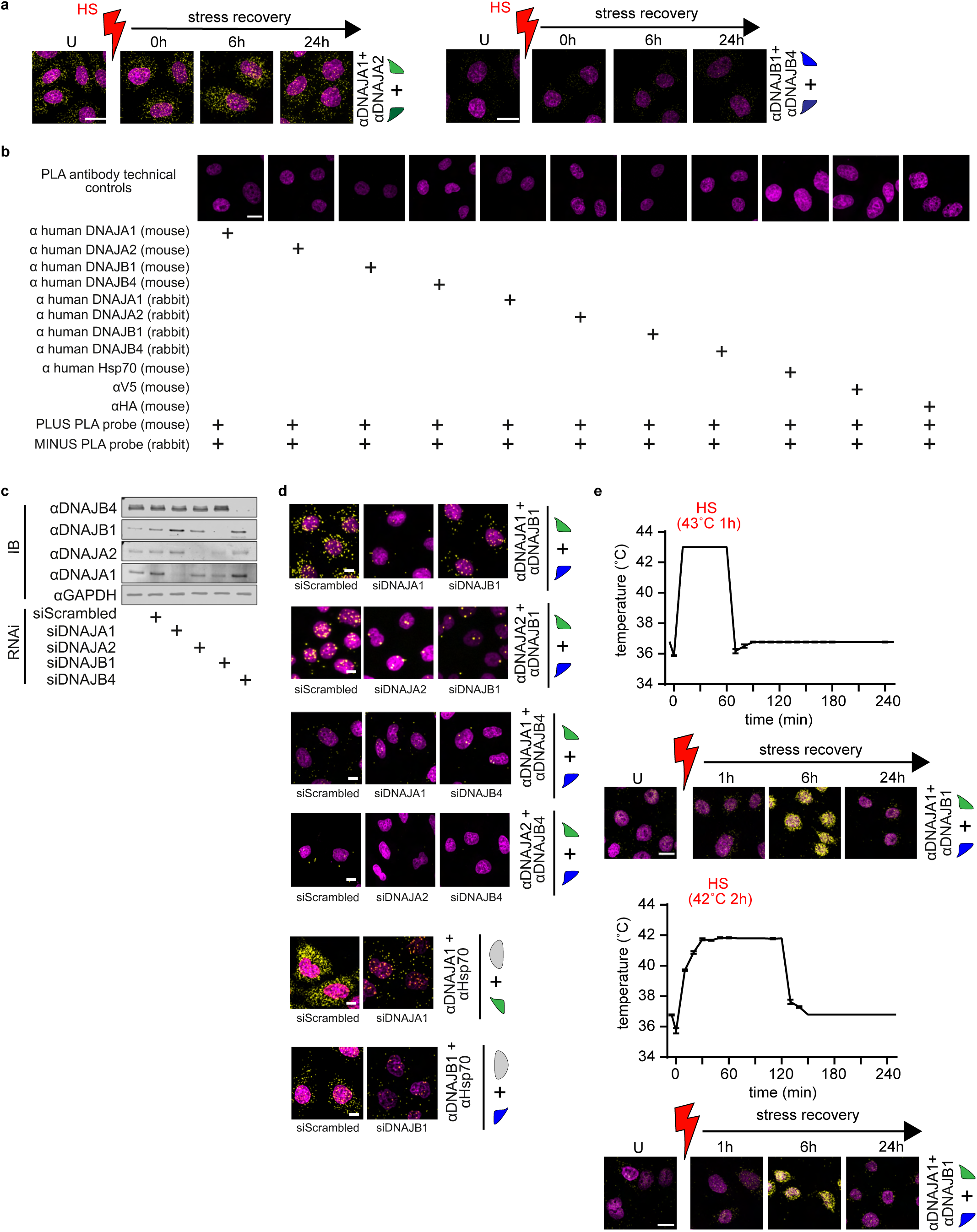
JDP-JDP and JDP-Hsp70 complex formation during disaggregase assembly monitored by PLA in heat shocked human cells. **a**, Formation of intraclass JDP scaffolds between DNAJA1 and DNAJA2, and DNAJB1 and DNAJB4 in HeLa cells during HS recovery. PLA signal in yellow. DAPI (magenta) staining indicates nuclei. HS indicated in red. Unstressed cells indicated as U. Scale bar 20 µm (n = 3). **b**, Technical controls of PLA for antibody specificity. PLA was performed in HeLa cells with single primary antibodies only. Signal amplification (yellow) was not observed in any of the controls. Scale bar 20 µm (n = 2). **c**, Immunoblot (IB) showing RNAi knockdown efficiencies for DNAJA1, DNAJA2, DNAJB1, DNAJB4 in HeLa cells. GAPDH, loading control. **d**, RNAi knockdown-based controls of PLA for antibody and complexation specificity. RNAi knockdown of single JDPs decreased JDP^A^-JDP^B^ and JDP^A/B^-Hsp70 complex formation (yellow puncta) in HeLa cells. The non-targeting control siRNA (siScrambled) is used as the control for RNAi. Scale bar 10 µm (n = 2). **e**, Top: The graphs show the temperature changes observed in two HS regimes (at 42 °C 2h in an incubator *vs.* 43 °C 1h in a water bath) used in the study. Bottom: Induction of the DNAJA1-DNAJB1 scaffold (PLA signal, yellow) after applying the two HS regimes to HeLa cells. Nuclei stained with DAPI (magenta). HS indicated in red. Unstressed cells indicated as U. Scale bar 20 µm (n = 3).

**Supplementary Data Figure 3.**
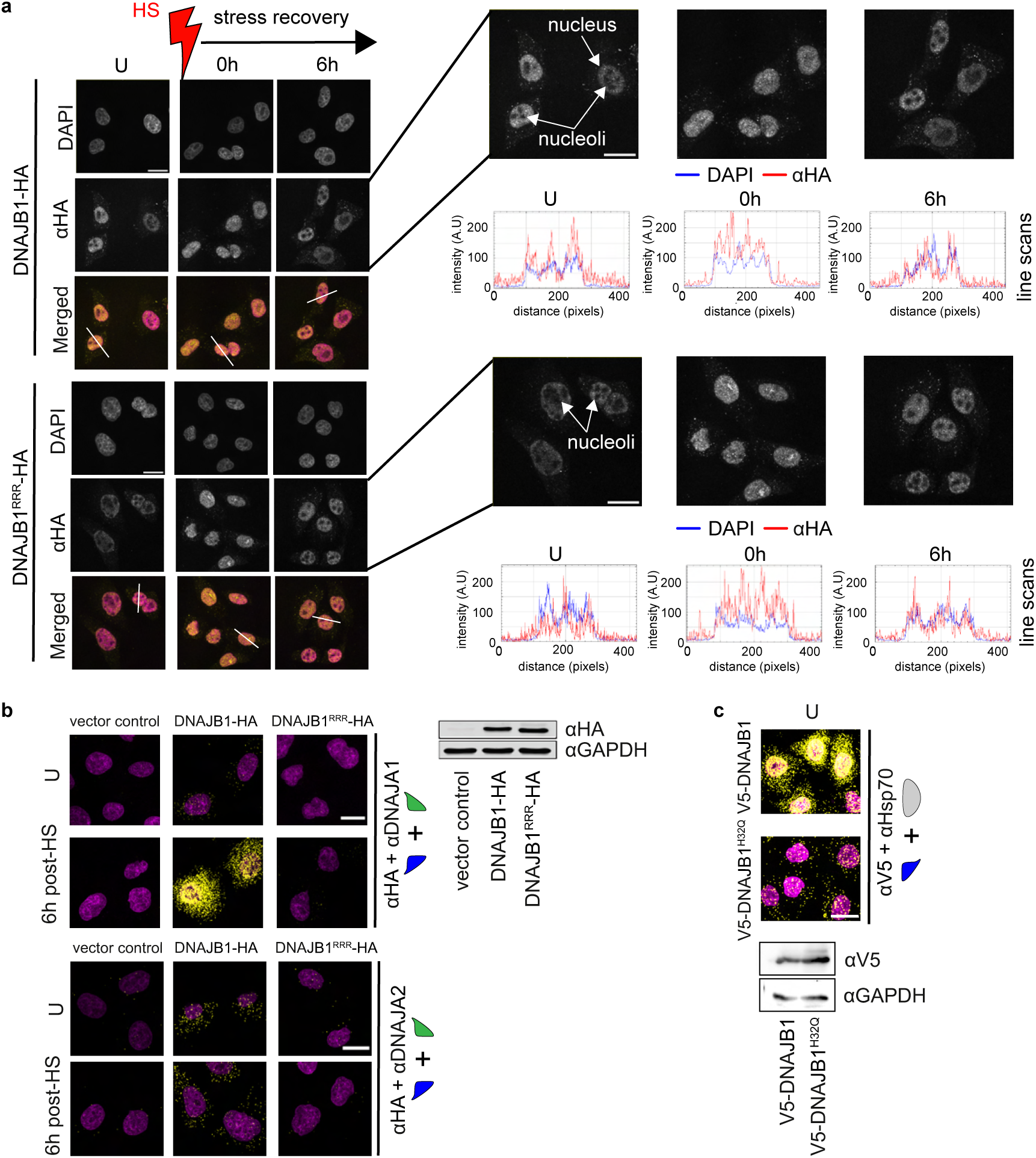
PLA signals capture direct interactions between disaggregase machine components during assembly. **a**, DNAJB1^RRR^ can shuttle in and out of nucleoli immediately after heat shock to support chaperoning function(s) similar to wild type DNAJB1. The immunocytochemical signal from the HA tag (yellow) shows that both DNAJB1-HA and the JDP scaffold forming defective mutant DNAJB1^RRR^-HA rapidly localize into the nucleoli of HeLa cells immediately after heat shock (HS), and relocate out of the nucleoli before 6h post-HS. RRR denotes the D4R + E69R + E70R mutations in DNAJB1. Unstressed cells are denoted as U. Nuclei stained with DAPI (magenta). The nucleoli are unstained by DAPI and appear as dark spots in the nucleus. Line scans across the nucleoli are indicated in white. Corresponding line scan plots are shown on the right. **b**, The PLA signal captures the direct interaction between DNAJA1 and DNAJB1 during the formation of the JDP scaffold. A decreased PLA signal (yellow) is observed for the interaction between the JDP scaffold forming defective mutant DNAJB1^RRR^-HA and endogenous DNAJA1 or DNAJA2. Nuclei stained with DAPI (magenta). Scale bar 20 µm (n = 2). Immunoblot shows expression levels of mutant and wild type DNAJB1. **c**, As in (**b**), a similar decrease in the PLA signal is observed for the interaction between Hsp70 binding defective mutant V5-DNAJB1^H32Q^ and endogenous Hsp70. Scale bar 20 µm (n = 2). Immunoblot shows expression levels of mutant and wild type DNAJB1.

**Supplementary Data Figure 4.**
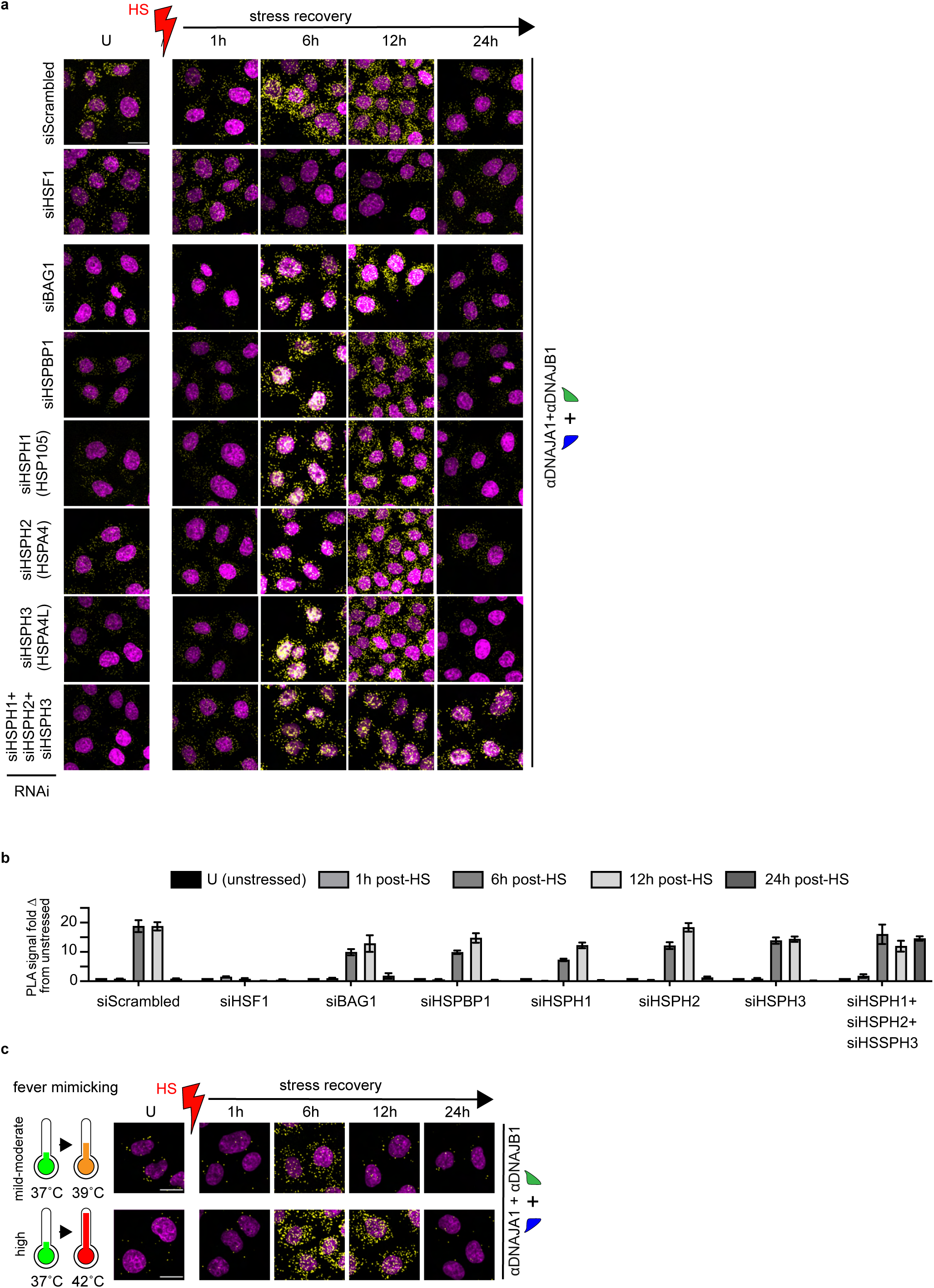
Hsp110-type nucleotide exchange factors (NEF) impact the disassembly of DNAJA1-DNAJB1 scaffold. The Hsp110 chaperone family of NEFs functions redundantly to facilitate the dissociation of ADP from Hsp70 and stimulate the disaggregation activity. **a**, Mini RNAi screen targeting NEFs from Hsp110 (HSPH1, HSPH2, and HSPH3), Bag (BAG-1), and HSPB1 (HSPB-1) families in HeLa cells. Single NEF depletions do not affect the assembly or disassembly of DNAJA1-DNAJB1 scaffold (PLA signal, yellow) during heat stress recovery (HS, red). Triple knockdown of the Hsp110s, which removes functional redundancy, results in delayed disassembly of the JDP scaffold. The non-targeting control siRNA (siScrambled) and depletion of HSF-1 (see section on HSF-1 mediated assembly of DNAJA-DNAJB1 scaffold) are used as controls for RNAi. Nuclei stained with DAPI (magenta). Scale bar 20 µm (n = 3). **b**, Quantification of the PLA signals from (a). Data are mean +/− s.e.m. **c**, Cells tune the activation of DNAJA1-DNAJB1 scaffold according to the level of protein damage sustained after HS. Heat stress at 39 °C *vs* 42 °C for 2h mimics mild-moderate and high fever, respectively. PLA signal in yellow. Quantification of PLA signal reported in Fig 1d. The nuclei stained with DAPI (magenta). Scale bar 20 µm (n = 3).

**Supplementary Data Figure 5.**
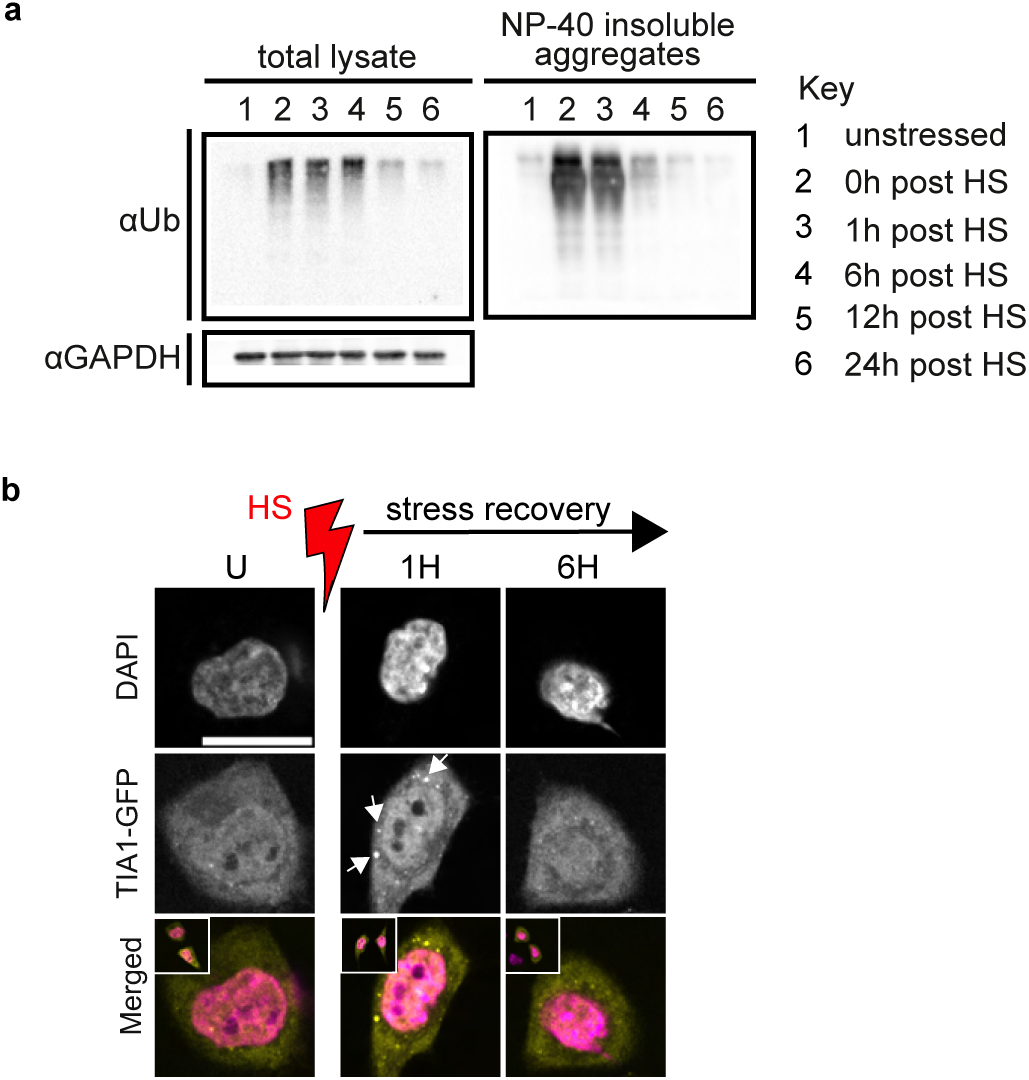
Resolution of different heat-induced aggregates/SGs in cells. **a**, Detection of heat-induced ubiquitin (Ub) positive protein aggregates. 1% NP-40 insoluble aggregates were isolated at indicated time points during heat stress recovery using a two-phase fractionation approach and probed with antibody against ubiquitin (n = 3). Immunoblots show the clearance of Ub-positive protein aggregates in HeLa cells before 6h post-HS. GAPDH, loading control. **b**, Resolution of TIA-1-GFP containing stress granules (SGs; yellow) also occurs in HeLa cells before 6h post-HS. Nuclei stained with DAPI (magenta) (n = 3).

**Supplementary Data Figure 6.**
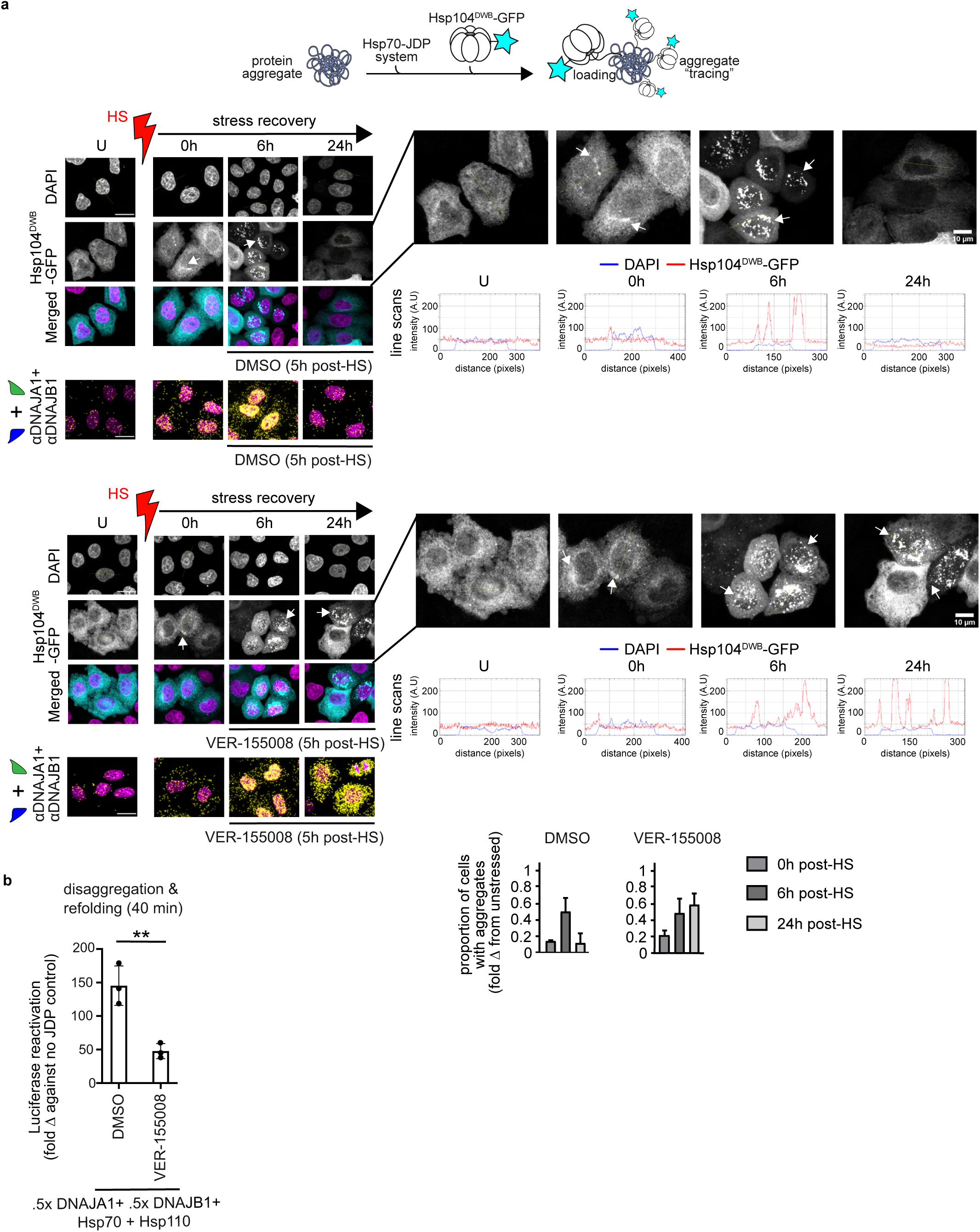
Heat-induced aggregate solubilization after inhibition of Hsp70 with VER-155008. **a**, Resolution of heat-induced aggregates in cells +/− Hsp70 inhibitor VER-155008. Top: Schematic diagram illustrating the decoration of heat induced aggregates by the yeast HSP104^DWB^-GFP. Hsp70-JDP chaperone system facilitates the loading of the inactive mutant of Hsp104 onto aggregate surface. Bottom: HS triggered the formation and solubilization of Hsp104^DWB^-GFP positive aggregates (cyan; punctated fluorescent signal pointed by white arrows) in HeLa cells. U denotes unstressed cells. Heat shock (HS) in red. Scale bar 20 µm (n = 3). Below: Visualization of assembly and disassembly of the DNAJA1-DNAJB1 scaffold (PLA; yellow) in HeLa cells recovering from heat shock (HS). The inhibitor was added to cells 5h after HS. DMSO, vehicle control for VER-155008. Nuclei in magenta. Scale bar 20 µm (n = 3). Quantification of Hsp104^DWB^-GFP positive aggregate levels in HeLa cells during HS recovery (see Methods). Data are mean +/− s.e.m. **b**, Inhibition of disaggregation/refolding of preformed luciferase aggregates by the Hsp70 inhibitor VER-155008, *in vitro* (n = 3, data are mean +/− s.e.m. ** adjusted p < 0.01, one-way ANOVA Tukey post hoc test).

**Supplementary Data Figure 7.**
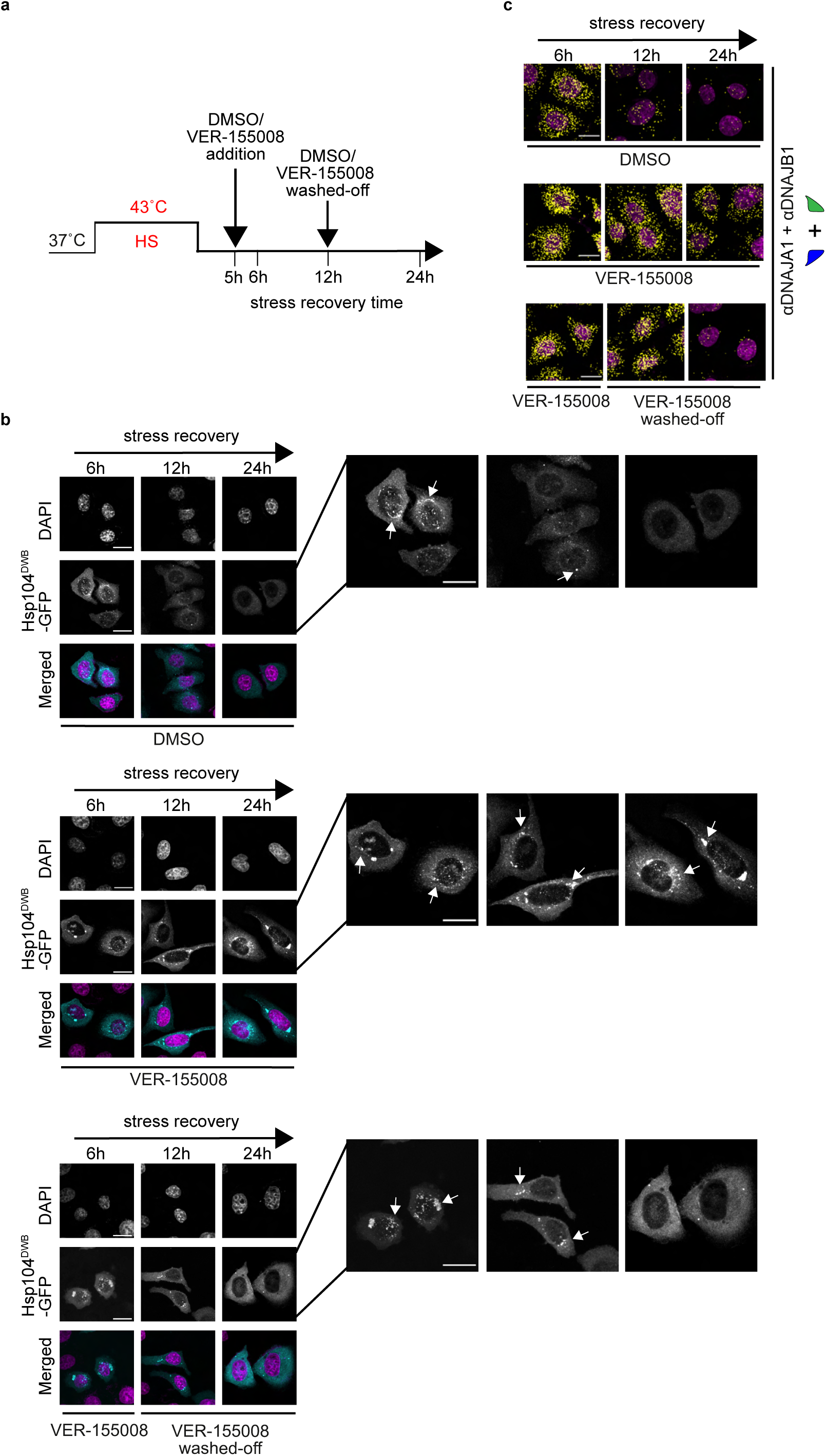
Disaggregation of heat-induced aggregates re-initiates after washing off VER-155008. **a**, VER-155008 wash-off experimental setup. **b**, Washing off VER-155008 by cell culture medium change at 12h post-HS results in the resumption of Hsp104^DWB^-GFP (cyan) positive aggregate solubilization in HeLa cells recovering from heat stress. VER-155008 was added to cells at 5h post-HS. DMSO, vehicle control for VER-155008. Nuclei stained with DAPI (magenta). Aggregates are indicated by white arrows. Scale bar 20 µm (n = 3). **c**, The assembled DNAJA1-DNAJB1 scaffolds (PLA signal, yellow) support the resumption of protein disaggregation after washing off the Hsp70 inhibitor. VER-155008 washed off HeLa cells show efficient disassembly of the DNAJA1-DNAJB1 scaffold of the disaggregase at 24h after HS compared to control with the inhibitor. DMSO, vehicle control for VER-155008. Nuclei stained with DAPI (magenta). Scale bar 20 µm. (n = 3).

**Supplementary Data Figure 8.**
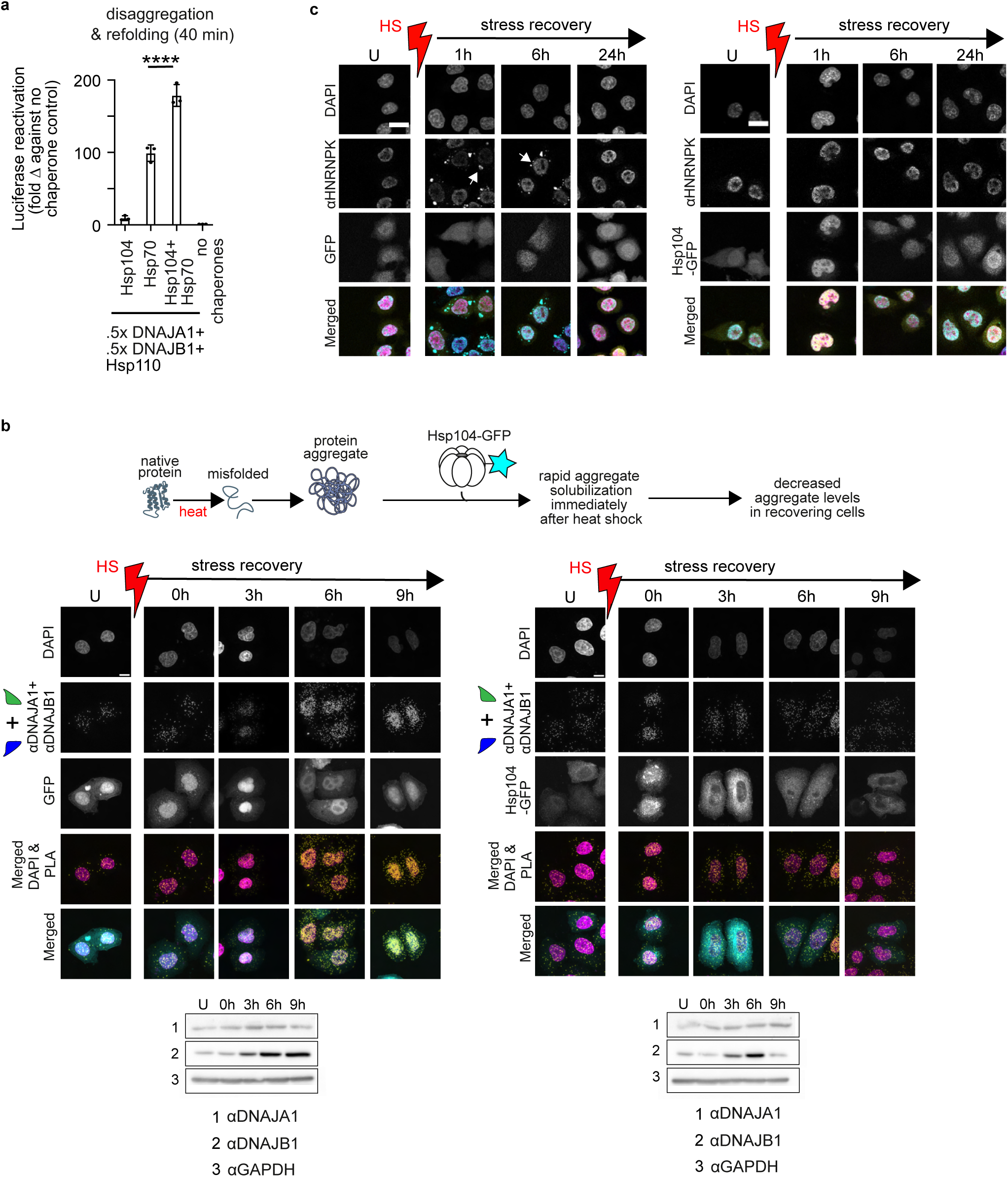
Expression of yeast Hsp104 disaggregase in human cells decreases the assembly of DNAJA1-DNAJB1 scaffold of the Hsp70 disaggregase. **a**, Yeast Hsp104 cooperates with human Hsp70-JDP system to target and solubilize preformed luciferase aggregates more effectively compared to Hsp70-JDP disaggregase alone, *in vitro* (n = 3, data are mean +/− s.e.m., **** adjusted p value < 0.0001, one-way ANOVA with Tukey post hoc test). Hsp110 functions as NEF. **b**, Assembly of DNAJA1-DNAJB1 scaffold of the Hsp70 disaggregase (PLA signal, yellow) is reduced in HeLa cells overexpressing wild-type Hsp104-GFP (cyan) compared to GFP control. Nuclei stained with DAPI (magenta). Heat shock (HS) in red. Scale bar = 20 µm (n = 3). Immunoblots show the induction of DNAJA1 and DNAJB1 after HS. GAPDH, loading control. **c**, Rapid solubilization of HNRNPK aggregates (cyan, indicated by white arrows, see section on endogenous substrates of Hsp70-DNAJA1-DNAJB1 disaggregase) immediately after heat shock in HeLa cells overexpressing Hsp104-GFP (yellow). Control cells express GFP alone (yellow). Nuclei stained with DAPI (magenta). Heat shock (HS) in red. Scale bar = 20 µm (n = 3).

**Supplementary Data Figure 9.**
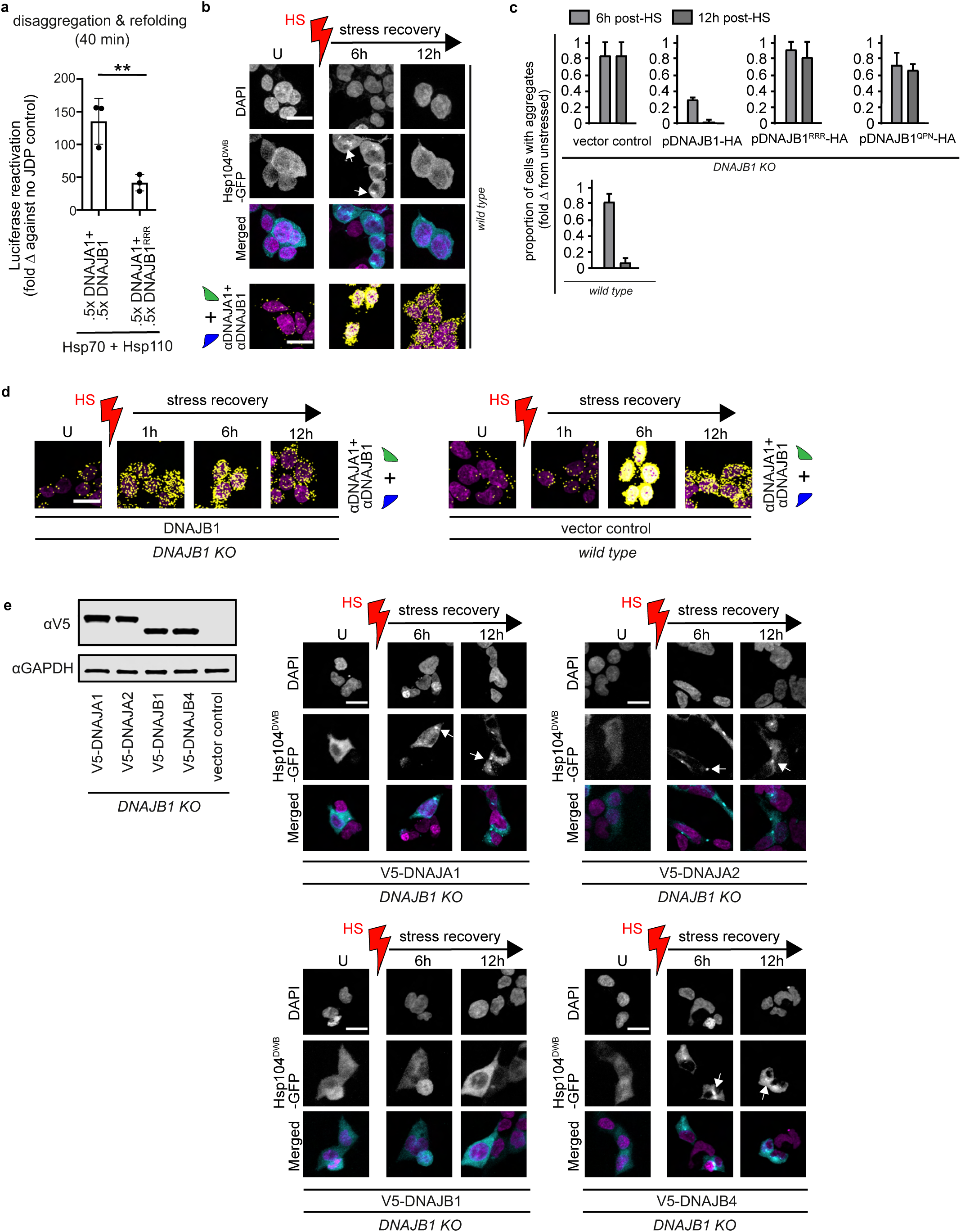
Cooperation between DNAJB1 and DNAJA1 is crucial for efficient Hsp70 disaggregase function in cells. **a**, The reaction containing class A JDP binding defective mutant DNAJB1^RRR^ (RRR denotes D4R+E69R+E70R) shows decreased disaggregation of heat-induced luciferase aggregates, *in vitro* (n = 3, data are mean +/− s.e.m. ** adjusted p-value < 0.01, one-way ANOVA Tukey post hoc test). **b**, Top: Solubilization of heat-induced Hsp104^DWB^-GFP (cyan) positive aggregates in HEK 293 cells recovering from heat stress. The nuclei stained with DAPI (magenta). Aggregated proteins are indicated by white arrows. Heat shock (HS) in red. Scale bar 20 µm (n = 3). Bottom: Visualization of the (dis)assembly dynamics of DNAJA1-DNAJB1 scaffold of the Hsp70 disaggregase (PLA signal, yellow) in HEK 293 cells recovering from heat shock. Scale bar 20 µm (n = 3). **c**, Quantification of heat-induced Hsp104^DWB^-GFP positive protein aggregates from Figure 1h-k and Supplementary Data Figure 9b (data are mean +/− s.e.m.). **d**, Rapid assembly of DNAJA1-DNAJB1 scaffold in heat shocked *DNAJB1* KO HEK 293 cells overexpressing DNAJB1 *vs* wild-type HEK 293 cells (PLA, yellow). Nuclei stained with DAPI (magenta). Scale bar 20 µm (n = 3). **e**, The role of DNAJB1 in Hsp70-mediated disaggregase function cannot be substituted for by other class A and B JDPs in stress recovering cells. Left: Overexpression levels of DNAJA1, DNAJA2, DNAJB1 and DNAJB4 tagged with V5 in *DNAJB1* KO HEK 293 cells visualized by immunoblotting. Right: Resolution of heat-induced Hsp104^DWB^-GFP positive aggregates (cyan) in *DNAJB1* KO cells ectopically expressing V5-tagged DNAJA1, DNAJA2, DNAJB1 or DNAJB4. Nuclei stained with DAPI (magenta). Aggregated proteins are indicated by white arrows. Scale bar 20 µm (n = 3).

**Supplementary Data Figure 10.**
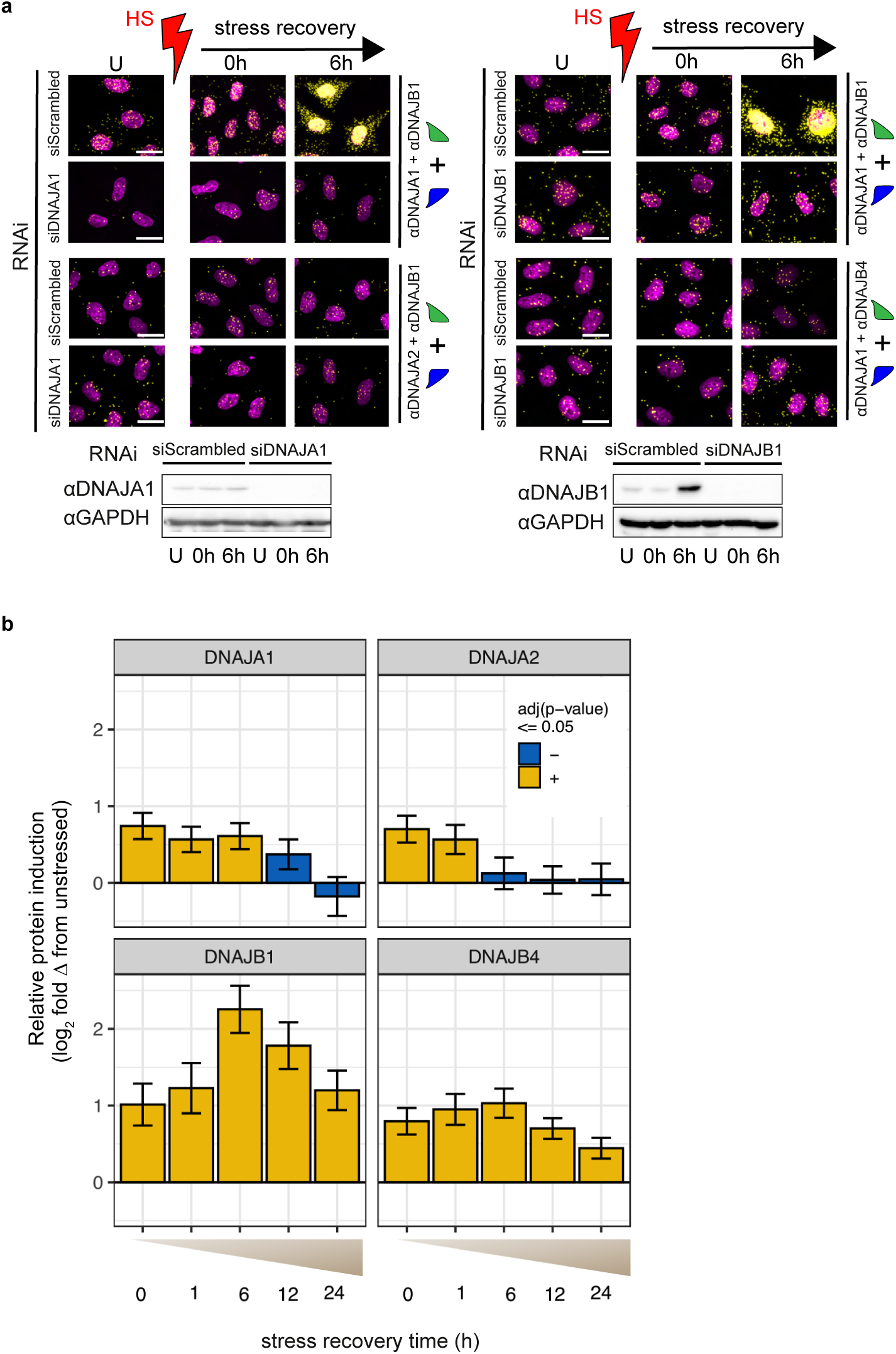
Heat induced Hsp70 disaggregase is cogged to assemble with DNAJA1-DNAJB1 scaffold. **a**, depleting either DNAJA1 or DNAJB1 via RNAi does not result in unpaired JDPs forming alternative scaffolds with DNAJA2 or DNAJB4. Left: Levels of DNAJA1-DNAJB1 and DNAJA2-DNAJB1 scaffolds in DNAJA1 depleted HeLa cells. Right: Levels of DNAJA1-DNAJB1 and DNAJA1-DNAJB4 scaffolds in DNAJB1 depleted HeLa cells. The non-targeting control siRNA (siScrambled) is used as the control for RNAi. Scale bar 20 µm (n = 3). Corresponding immunoblots show JDP knockdown efficiencies. GAPDH, loading control. **b**, HS triggered increase in DNAJA1, DNAJA2, DNAJB1 and DNAJB4 protein levels in recovering HeLa cells measured through multiple reaction monitoring (MRM, see Methods) mass spectrometry (normalized to tubulin alpha-1A and unstressed JDP levels). Error bars reflect fold change +/− s.e. of regression (n = 4).

**Supplementary Data Figure 11.**
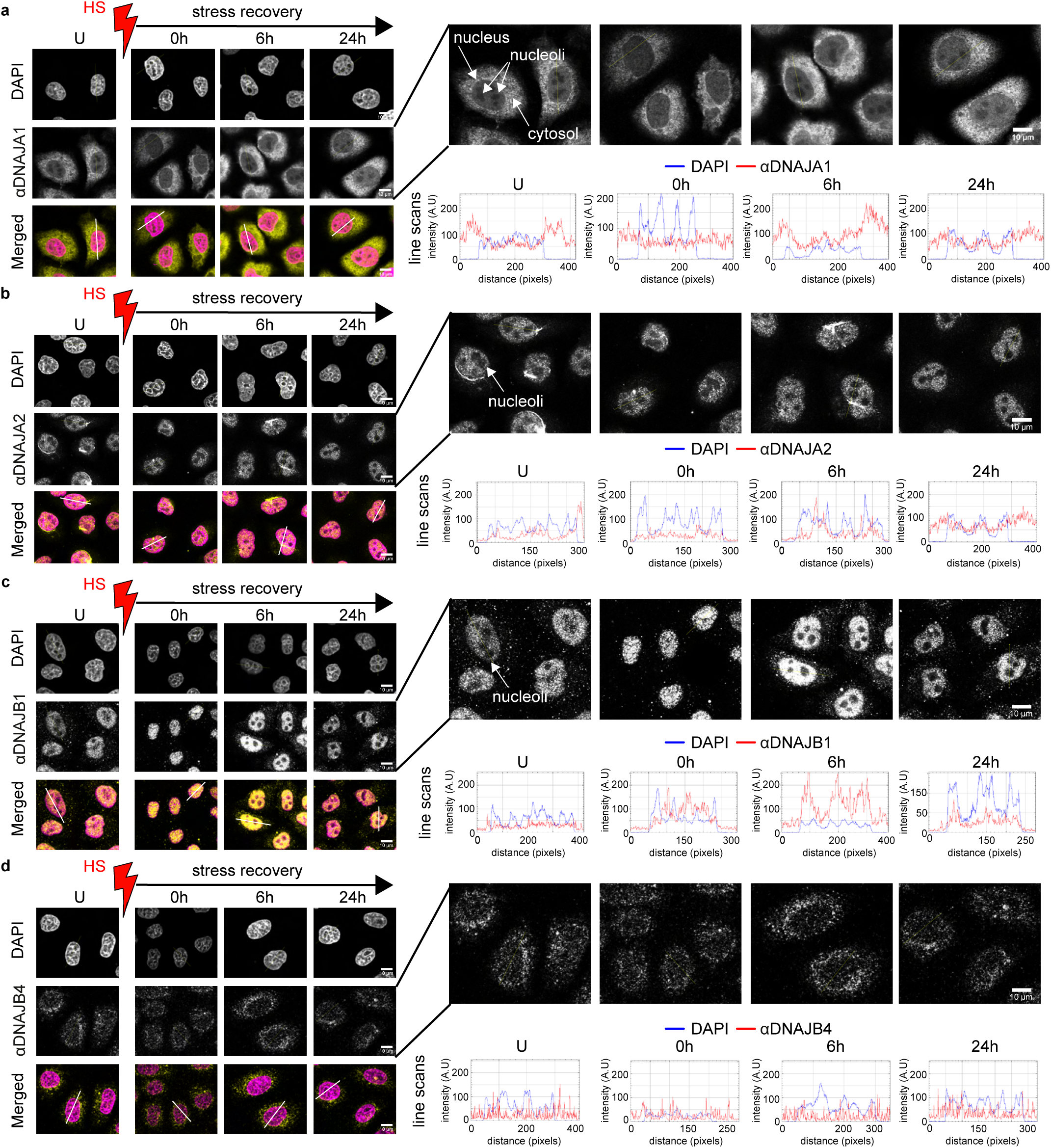
Localization of DNAJA1, DNAJA2, DNAJB1 and DNAJB4 in unstressed and heat shocked cells. **a**, DNAJA1 is localized in both cytosol and nucleus in unstressed (U) HeLa cells monitored by immunocytochemistry (yellow). DNAJA1 rapidly localizes into nucleoli immediately after heat shock, and relocates out of nucleoli before 6h post-HS. Nuclei stained with DAPI (magenta). The nucleoli are unstained by DAPI and appear as dark spots in the nucleus. Line scans across the nucleoli are indicated in white. Corresponding line scan plots are shown on the right. Scale bar 10 µm (n = 3). (**b-d**), As in (**a**). (**b**) DNAJA2 shows a predominantly nuclear localization signal in both unstressed and heat shocked cells. (**c**) DNAJB1 is localized predominantly in the nucleus in unstressed (U) cells and rapidly localizes into nucleoli immediately after heat shock, and relocates out of nucleoli before 6h post-HS. (**d**) DNAJB4 displays both cytosolic and nuclear localization and does not show detectible heat stress induced dynamics in HeLa cells.

**Supplementary Data Figure 12.**
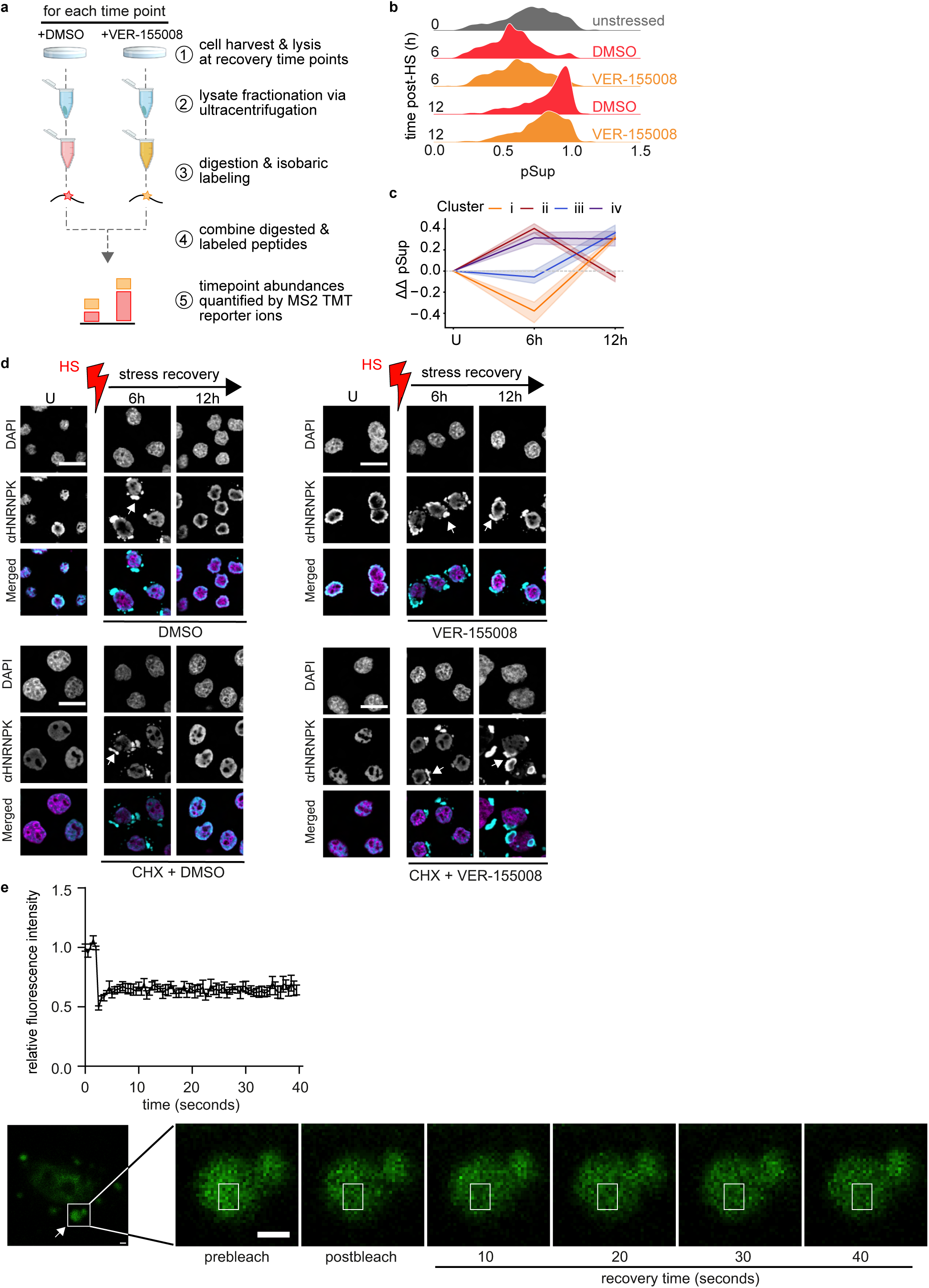
Identification of aggregated proteins recovered by stress-induced Hsp70 disaggregase activity. **a**, Schematic representation of the proteomics workflow used to identify aggregated proteins rescued by stress-induced Hsp70 disaggregase. The fractions were distinguished by isobaric labeling and proteins that regained solubility between 6-12h in DMSO, but not in cells treated with VER-155008 were identified and quantified by LC-MS/MS (see Methods). **b**, Ridge plot showing the distribution of solubility (pSup) for all proteins during post-HS recovery. **c**, Proteins for which the change in relative solubility due to the VER-155008 treatment (ΔΔpSup) exceeding 0.26 in at least one time point were clustered via k-means into four patterns. Mean ΔΔpSup for all proteins in each cluster is shown, where colored bands represent ± s.d. (n = 4). **d**, Visualization of Hsp70-based disaggregation of heat aggregated HNRNPK in HeLa cells by immunocytochemistry (HNRNPK, cyan; DAPI stained nuclei, magenta). Heat-induced Hsp70 disaggregase was inhibited with VER-155008 +/− cycloheximide (CHX) at 5h after HS. CHX inhibits protein synthesis. The white arrows indicate HNRNPK aggregated with heat. Aggregate solubilization is indicated by the disappearance of the punctated fluorescence signal and the appearance of a diffused signal. DMSO, vehicle control for VER-155008. Heat shock (HS) in red. U denotes unstressed cells. Scale bar 20 µm (n = 3). **e**, Fluorescence recovery after photobleaching (FRAP) analysis of heat-induced HNRNPK-containing aggregates shows poor recovery of the fluorescent signal after photobleaching. GFP-HNRNPK was transiently expressed in HeLa cells, and FRAP was performed after heat shock. The fluorescence signal of the selected inclusions was reduced to ~50% by photobleaching with the 488-nm laser (photobleached area indicated by the white squire), and its recovery was recorded for 40 seconds. The fluorescence intensity of GFP-HNRNPK after photobleaching was normalized as described in Methods. Selected images of a cell containing GFP-HNRNPK positive aggregates (green) before photobleaching and at indicated intervals after photobleaching are shown. The white arrows indicate GFP-HNRNPK aggregates. Scale bar 2 µm (n = 7).

**Supplementary Data Figure 13.**
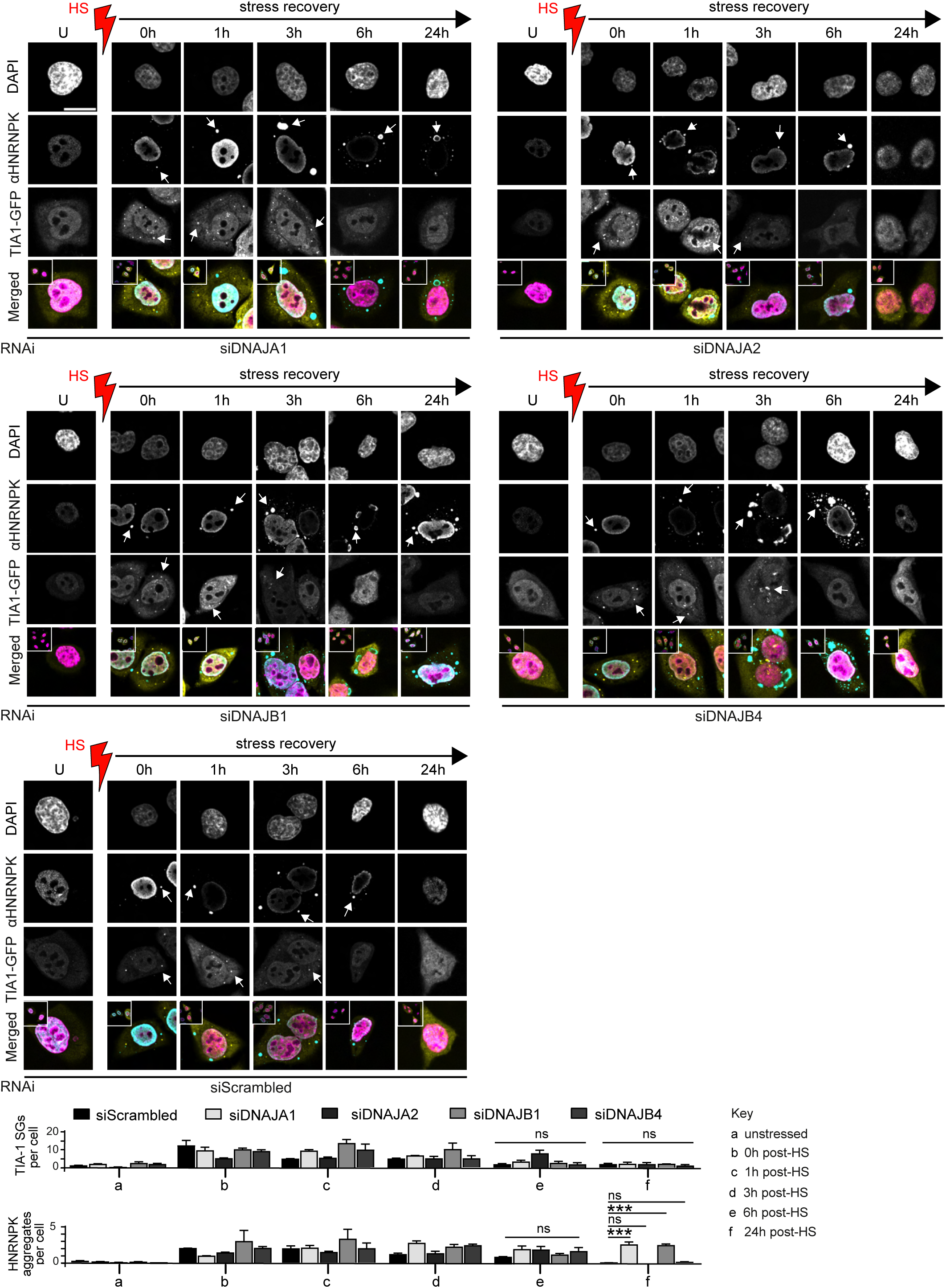
TIA-1 containing stress granules disassembly is not affected in cells depleted of class A and B JDPs. Depletion of DNAJA1, DNAJA2, DNAJB1, and DNAJB4 via RNAi has no effect on the resolution of TIA-1 SGs (yellow) in heat stress recovering HeLa cells. DNAJA1 and DNAJB1, but not DNAJA2 and DNAJB4 knockdown results in inhibition of HNRNPK (cyan) aggregate solubilization. The inset depicts zoomed out cells. The non-targeting control siRNA (siScrambled) is used as the control for RNAi. Nuclei stained with DAPI (magenta). Scale bar 20 µm. Below: Quantification of TIA-1 SGs and HNRNPK aggregates (n = 3, data are mean +/− s.e.m. * adjusted p-value < 0.05, ** adjusted p-value < 0.01, ns - not significant, one-way ANOVA LSD post hoc test).

**Supplementary Data Figure 14.**
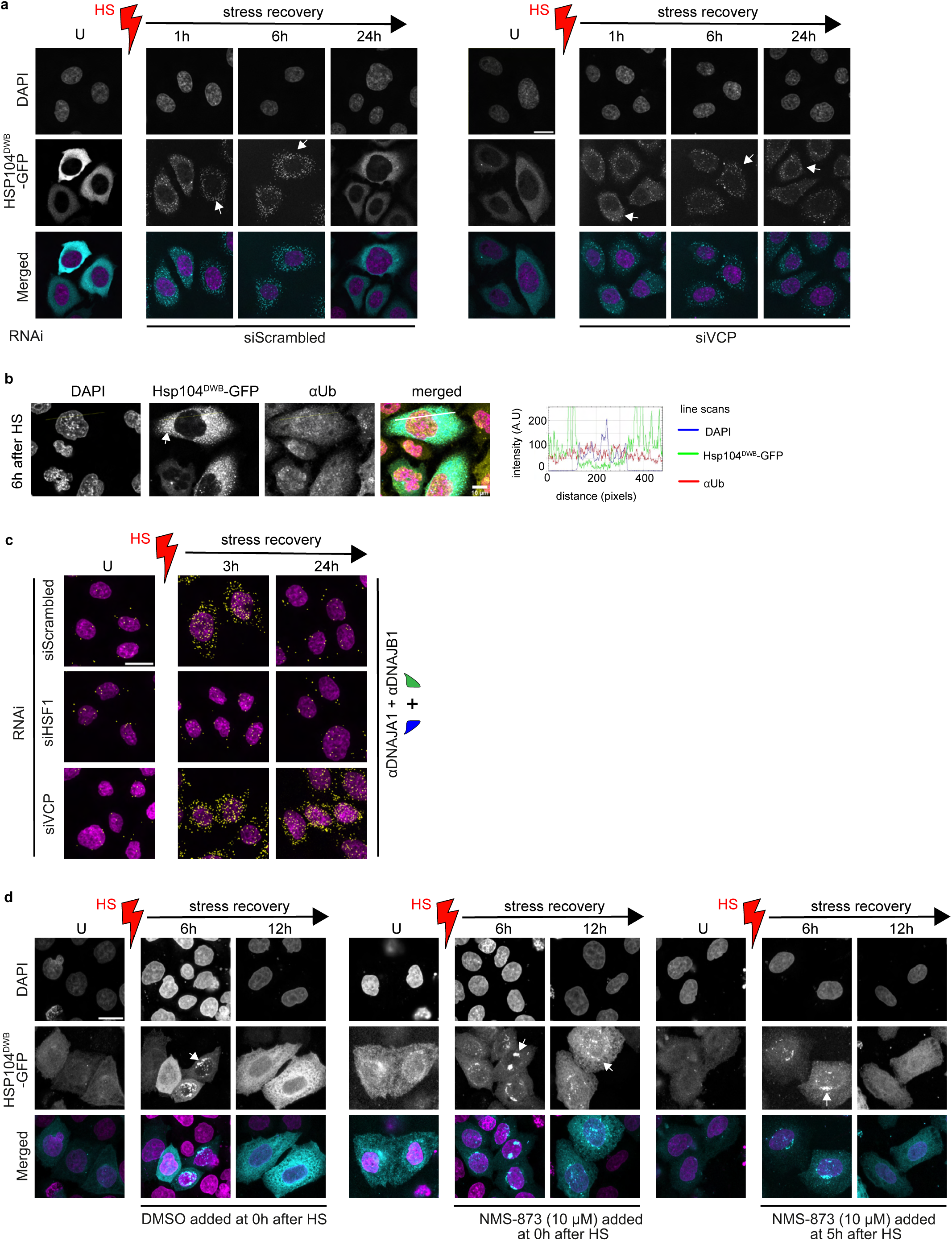
Resolution of Hsp104^DWB^-GFP positive aggregates in VCP depleted/inhibited cells. **a**, Resolution of heat-induced Hsp104^DWB^-GFP positive aggregates (cyan; punctated fluorescent signal pointed by white arrows) is blocked in HeLa cells RNAi depleted of VCP. The non-targeting control siRNA (siScrambled). Scale bar 20 µm (n = 3). Nuclei stained with DAPI (magenta). Heat shock (HS) in red. U denotes unstressed cells. **b**, Heat-induced Hsp104^DWB^-GFP positive aggregates do not have a ubiquitin signal. Colocalization of the immunofluorescence signal from ubiquitin (Ub, yellow) and the fluorescence signal from Hsp104^DWB^-GFP (cyan) in HeLa cells recovering from heat shock (HS) (6h post-HS). Nuclei stained with DAPI (magenta). Line scans are indicated in white. Corresponding line scan plots are shown on the right. Scale bar 10 µm (n = 3). **c**, Disassembly of Hsp70 disaggregase is delayed in VCP depleted HeLa cells (PLA signal, yellow). The non-targeting control siRNA (siScrambled) and depletion of HSF-1 are used as controls for RNAi (see section on HSF-1 mediated assembly of DNAJA-DNAJB1 scaffold). Nuclei stained with DAPI (magenta). Heat shock (HS) in red. U denotes unstressed cells. Scale bar 20 µm (n = 3). **d**, Blocking VCP disaggregase activity with inhibitor NMS-873 (10 µM, high dose) after 5h post-HS does not inhibit the resolution of Hsp104^DWB^-GFP positive aggregates by Hsp70-DNAJA1-DNAJB1 disaggregase. On the contrary, the addition of NMS-873 at 0h post-HS results in a phenotype similar to the depletion of VCP. DMSO, vehicle control for the inhibitor (n = 3). Nuclei stained with DAPI (magenta).

**Supplementary Data Figure 15.**
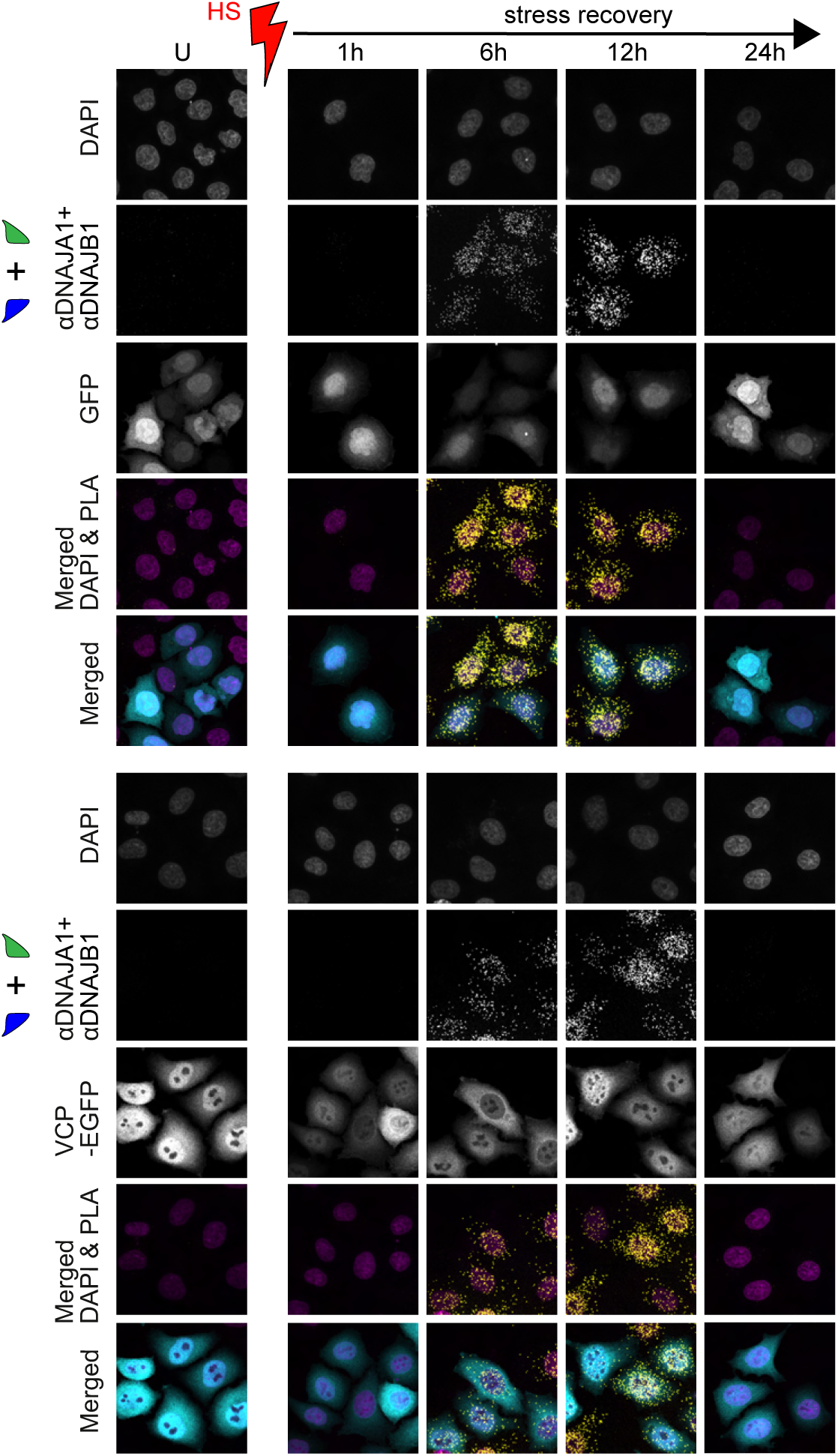
Cells overexpressing VCP still activate Hsp70-DNAJA1-DNAJB1 disaggregase during heat stress recovery. The assembly and disassembly dynamics of DNAJA1-DNAJB1 scaffold of Hsp70 disaggregase is not affected in HeLa cells overexpressing VCP (PLA signal, yellow; VCP-EGFP signal, cyan). GFP-alone is expressed as control. Nuclei stained with DAPI (magenta). Scale bar 20 µm (n = 3).

**Supplementary Data Figure 16.**
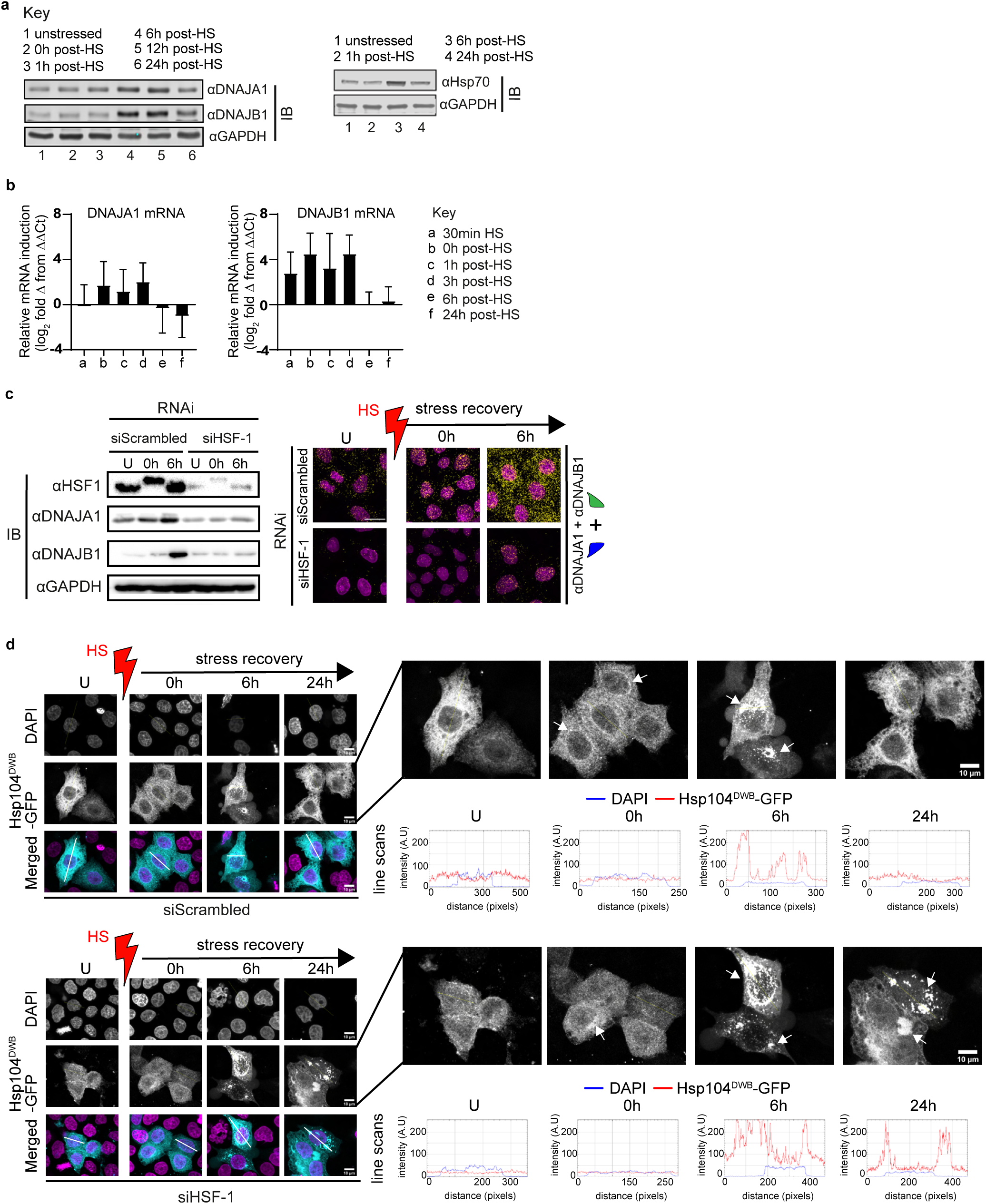
Disaggregation of heat-induced Hsp104^DWB^-GFP positive aggregates requires an intact heat shock response. **a**, Immunoblots showing changes in steady-state protein levels of DNAJA1, DNAJB1 and Hsp70 after HS in HeLa cells. **b**, Induction of DNAJA1 and DNAJB1 mRNA measured by qRT-PCR (log_2_ fold (-ΔCt; relative to GAPDH mRNA) in HeLa cells recovering from HS. s.e.m. calculated from ΔCt value per experiment. Error bars reflect mean +/− s.e.m. (n = 3). **c**, Inhibition of DNAJA1 and DNAJB1 protein synthesis after HS through HSF-1 depletion blocks DNAJA1-DNAJB1 scaffold assembly. Left: Immunoblots show HSF-1 knockdown by RNAi and inhibition of JDP synthesis after HS. GAPDH, loading control. Right: PLA shows a decreased assembly of Hsp70 disaggregase in HSF-1-depleted cells. The non-targeting control siRNA (siScrambled) is used as the control for RNAi. Scale bar 20 µm (n = 3). **d**, Disruption of the heat shock response by depleting HSF-1 blocks solubilization of heat-induced Hsp104^DWB^-GFP (cyan) positive aggregates in recovering HeLa cells. Aggregated proteins are visualized as punctated fluorescence signals and indicated by white arrows. Nuclei stained with DAPI (magenta). Line scans across the nucleoli are indicated in white. Corresponding line scan plots are shown on the right. Scale bar 10 µm (n = 3).

**Supplementary Data Figure 17.**
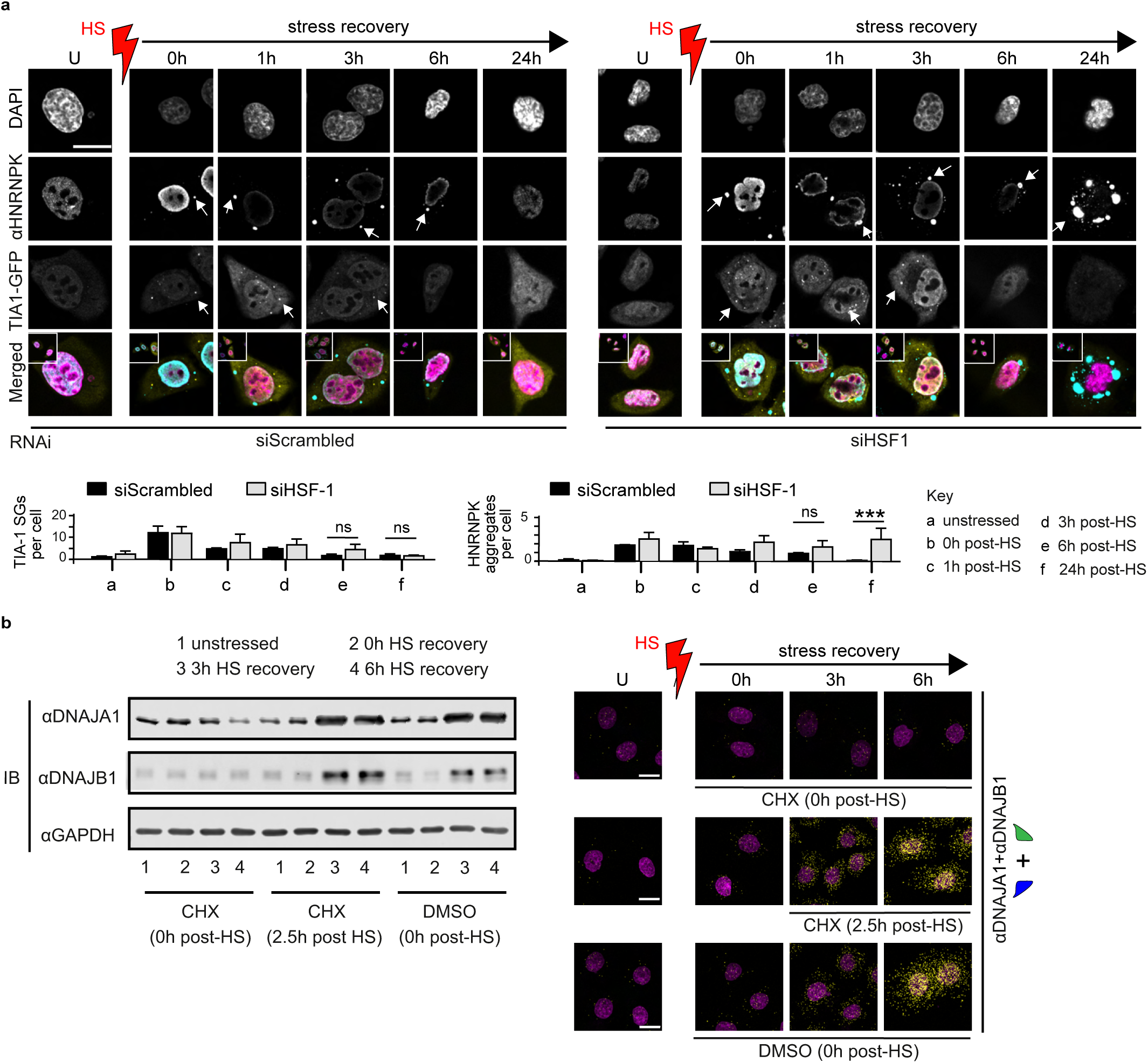
Resolution of different heat-induced aggregate/SG types in cells. **a**, The resolution of TIA-1 containing stress granules occurs independent of the heat shock response. Inactivation of the heat shock response through depletion of HSF-1 does not affect the disassembly of TIA-1-GFP containing SGs (yellow). In contrast, activation of the stress signaling pathway is required to solubilize heat-induced HNRNPK aggregates (cyan). HSF-1 depletion results in inhibition of the disaggregation of HNRNPK aggregates in HeLa cells. The inset depicts zoomed out cells. The non-targeting control siRNA (siScrambled) is used as the control for RNAi. Nuclei stained with DAPI (magenta). Scale bar 20 µm. Scale bar 20 µm. Below: Quantification of TIA-1 SGs and HNRNPK aggregates (n = 3, data are mean +/− s.e.m. * adjusted p-value < 0.05, ** adjusted p-value < 0.01, ns - not significant, one-way ANOVA LSD post hoc test). **b**, DNAJA1 and DNAJB1 molecules synthesized before HS are not involved in Hsp70 disaggregase formation. Left: Cycloheximide (CHX) is added to HeLa cells to block the induction of DNAJA1 and DNAJB1 at specific time points during recovery from heat stress. DNAJA1 and DNAJB1 induction after heat stress is shown by immunoblotting. GAPDH, loading control. DMSO, vehicle control for CHX (n = 2). Right: The newly synthesized DNAJA1 and DNAJB1 molecules are used to assemble the DNAJA1-DNAJB1 scaffold of Hsp70 disaggregase after heat shock (HS, red). The addition of CHX at 0h after HS, which inhibits the synthesis of new DNAJA1 and DNAJB1 molecules, completely blocks the assembly of the JDP scaffold (PLA signal, yellow). Treatment of cells with CHX at 2.5h post-HS does not prevent the synthesis of new DNAJA1 and DNAJB1 molecules, and assembly of the JDP scaffold. Nuclei stained with DAPI (magenta). DMSO, vehicle control for CHX. Scale bar 20 µm (n = 3).

**Supplementary Data Figure 18.**
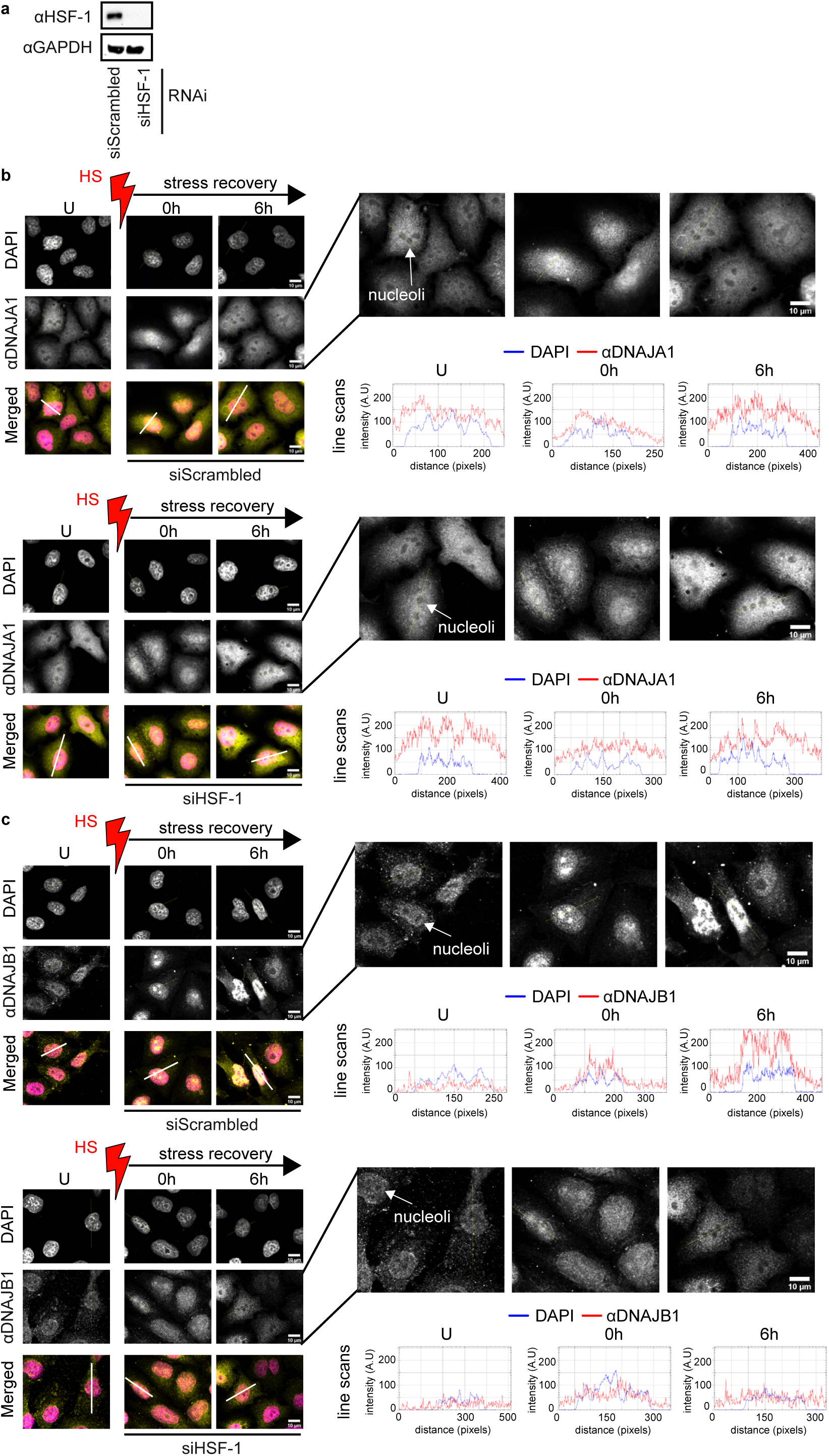
The localization of DNAJA1 and DNAJB1 into nucleoli immediately after heat shock occurs independent of the heat shock response. **a**, Cellular HSF-1 levels after RNAi knockdown are shown by immunoblot. GAPDH, loading control. The non-targeting control siRNA (siScrambled) is used as the control for RNAi. **b**, DNAJA1 molecules synthesized prior to heat stress rapidly localize into nucleoli immediately after heat shock (HS, red), and relocate out of nucleoli before 6h post-HS at similar levels in both HSF-1 depleted and control (siScrambled) HeLa cells. Subcellular localization of DNAJA1 in unstressed (U) and heat shocked (HS) HeLa cells monitored by immunocytochemistry (yellow). Nuclei stained with DAPI (magenta). The nucleoli are unstained by DAPI and appear as dark spots in the nucleus. Line scans across the nucleoli are indicated in white. Corresponding line scan plots are shown on the right. Scale bar 10 µm (n = 3). **c**, As in (**b**), DNAJB1 molecules synthesized prior to HS rapidly locate into nucleoli immediately after HS, and relocates out of nucleoli before 6h post-HS to a similar degree in HSF-1 depleted and control cells. Scale bar 10 µm (n = 3).

**Supplementary Data Figure 19.**
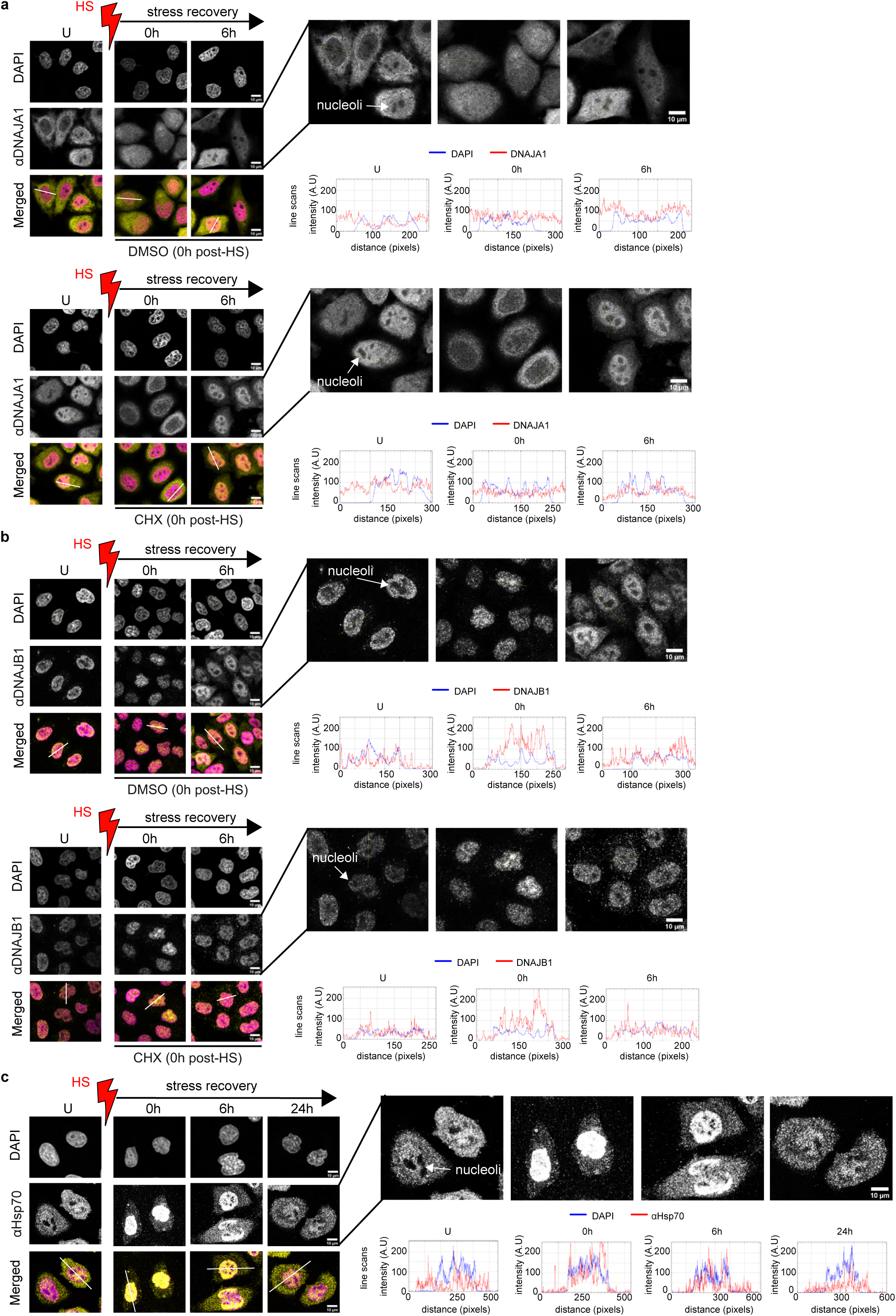
Pre-HS synthesized DNAJA1 and DNAJB1 shuttle to nucleoli immediately after heat stress. **a**, DNAJA1 molecules predominantly synthesized before heat stress show similar level of localization into nucleoli immediately after heat shock (HS, red), and relocate out of the nucleoli before 6h post-HS. Cycloheximide (CHX) was added to inhibit protein synthesis at 0h after HS. DMSO added at 0h post-HS as vehicle control. DNAJA1 detected in unstressed cell (U) and heat shocked HeLa cells by immunocytochemistry (yellow). Nuclei stained with DAPI (magenta). The nucleoli are unstained by DAPI and appear as dark spots in the nucleus. Line scans across the nucleoli are indicated in white. Corresponding line scan plots are shown on the right. Scale bar 10 µm (n = 3). **b**, As in (**a**), Pre-HS synthesized DNAJB1 molecules in CHX treated cells rapidly locate into nucleoli immediately after HS, and relocate out of nucleoli before 6h post-HS similar to control (DMSO treated) cells. Scale bar 10 µm (n = 3). **c**, Hsp70 molecules rapidly localize into nucleoli immediately after heat shock, and relocate out of the nucleoli before 6h post-HS. Scale bar 10 µm (n = 3).

**Supplementary Data Figure 20.**
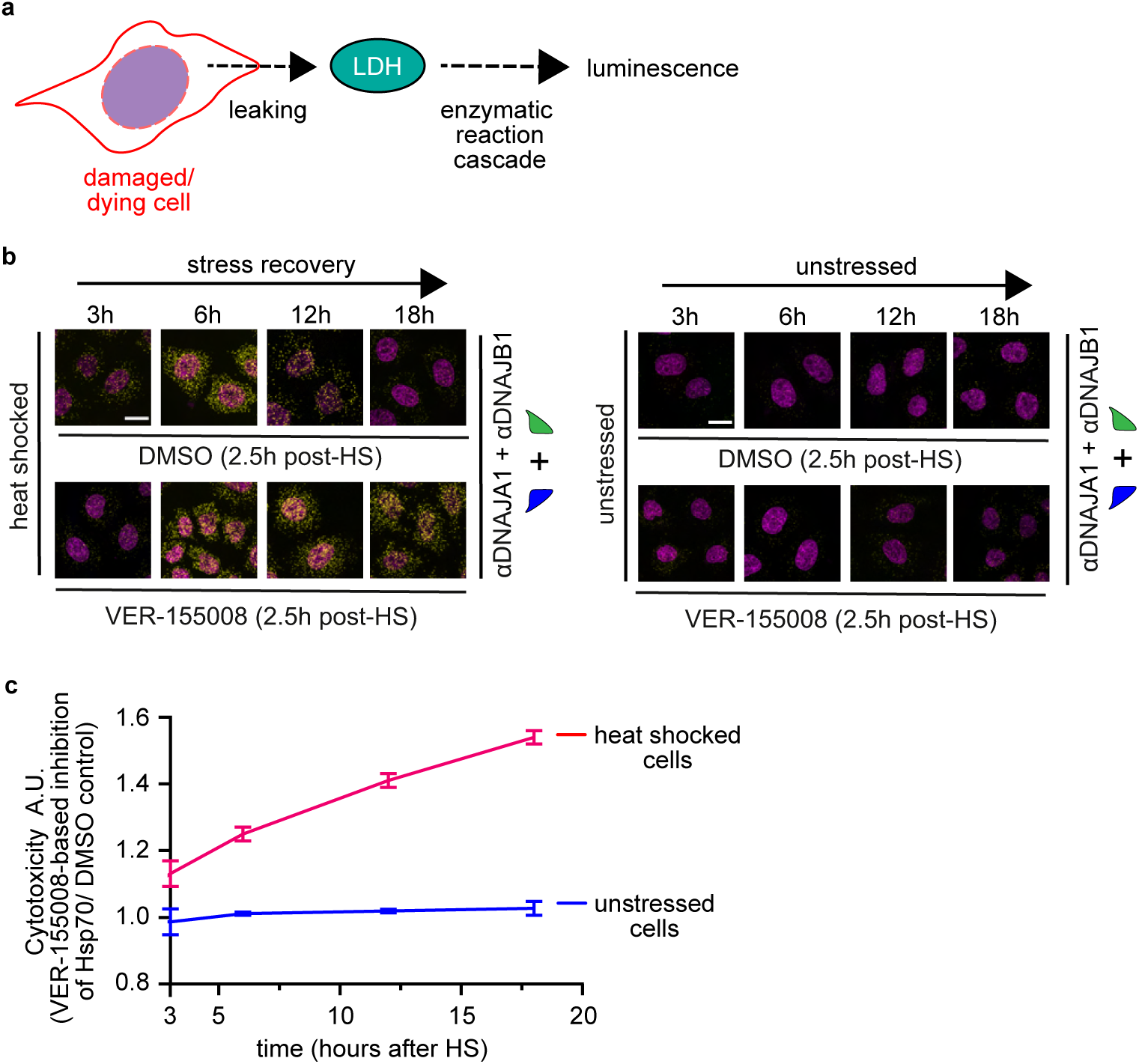
Stress-induced Hsp70 disaggregase provides heat tolerance. **a**, Simplified scheme of Ultra-Glo rLuciferase-based cytotoxicity assay. Leaked LDH is detected as a readout for damaged or dying cells. **b**, Assembly of DNAJA1-DNAJB1 scaffold of the Hsp70 disaggregase (PLA, yellow) in cells treated with VER-155008 at 2.5h after HS. Control, cells without HS. DMSO, vehicle control for VER-155008. The nuclei stained with DAPI (magenta). Scale bar 20 µm (n = 3). **c**, Disruption of stress-induced Hsp70 disaggregase activity increased heat toxicity in cells. Cytotoxicity level in unstressed (blue) and HS recovering HeLa cells (red) with and without VER-155008 treatment. DMSO, vehicle control for the inhibitor. Error bars depicted as s.e.m. (n = 3).

**Supplementary Data Figure 21.**
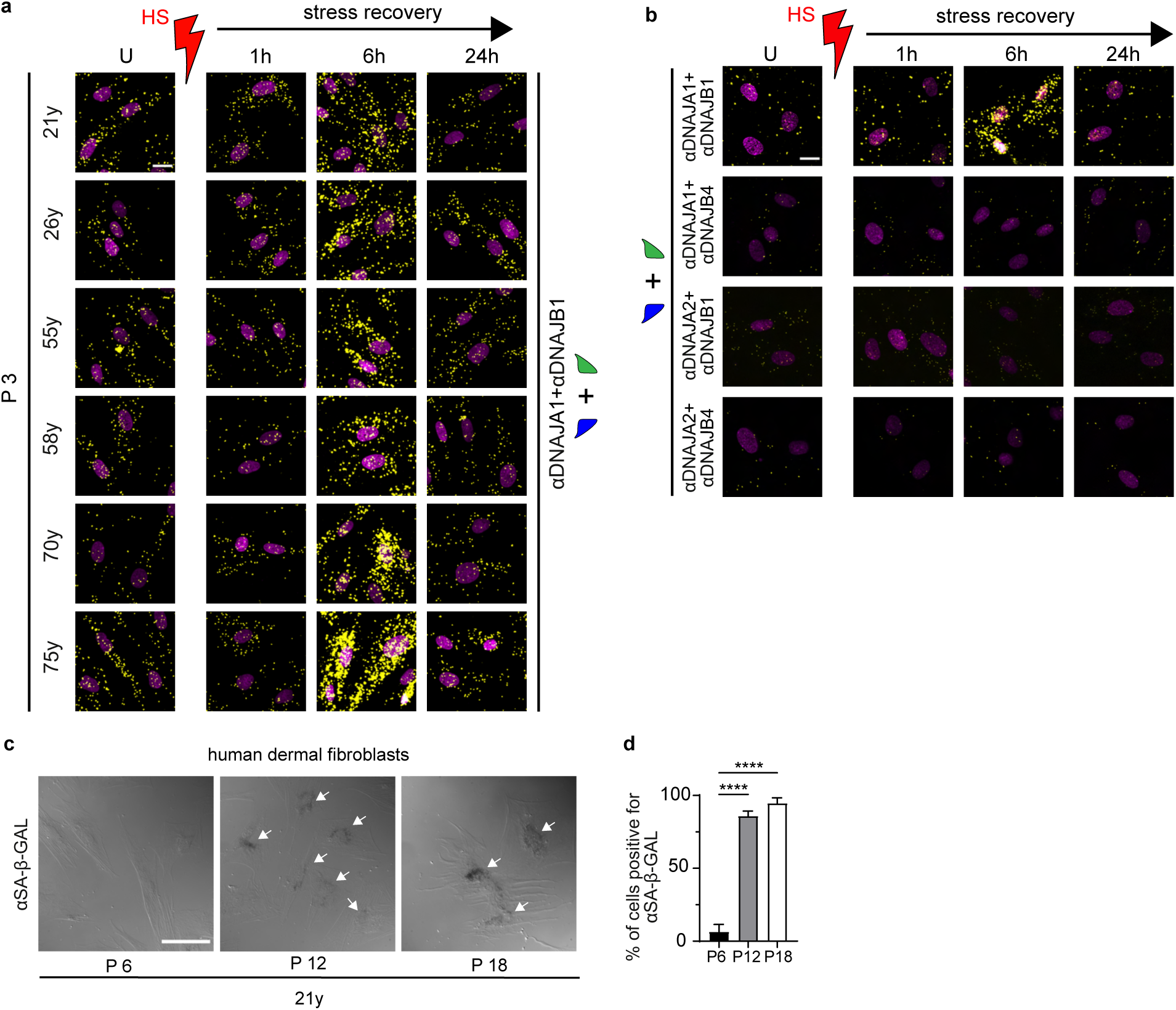
Assembly of Hsp70-DNAJA1-DNAJB1 disaggregase is not affected in dermal fibroblasts obtained from young *vs* old human donors. **a**, Induction of the DNAJA1-DNAJB1 scaffold of Hsp70 disaggregase in heat shocked primary dermal fibroblasts (HDFs) derived from humans of different chronological ages (21 years of age (21y), 26y, 55y, 58y, 70y, and 75y) in cells at passage (P) 3. Nuclei stained with DAPI (magenta). U denotes unstressed cells. Heat shock (HS) in red. Scale bar 20 µm (n = 3). **b**, The assembly dynamics of different JDP scaffolds formed between members of class A (DNAJA1 and DNAJA2) and class B (DNAJB1 and DNAJB4) (PLA signal, yellow) in human dermal fibroblasts after HS. Nuclei stained with DAPI (magenta). Scale bar 20 µm. (n = 3). **c**, Dermal fibroblasts undergoing replicative aging show increased senescence-specific β-galactosidase levels. Detection of senescence-specific β-galactosidase levels in P 6, P 12 and P 18 HDFs derived from 21-year-old (21y) human donor (see Methods). DIC images show cells at P 12 and P 18 stained positive for senescence-specific β-galactosidase (white arrows). Scale bar 20 µm. (n = 4). **d**, Quantification of senescence-specific β-galactosidase positive cells at P 6, P 12 and P 18. Data are mean +/− s.e.m. **** adjusted p-value < 0.0001, one-way ANOVA Tukey post hoc test).

**Supplementary Data Figure 22.**
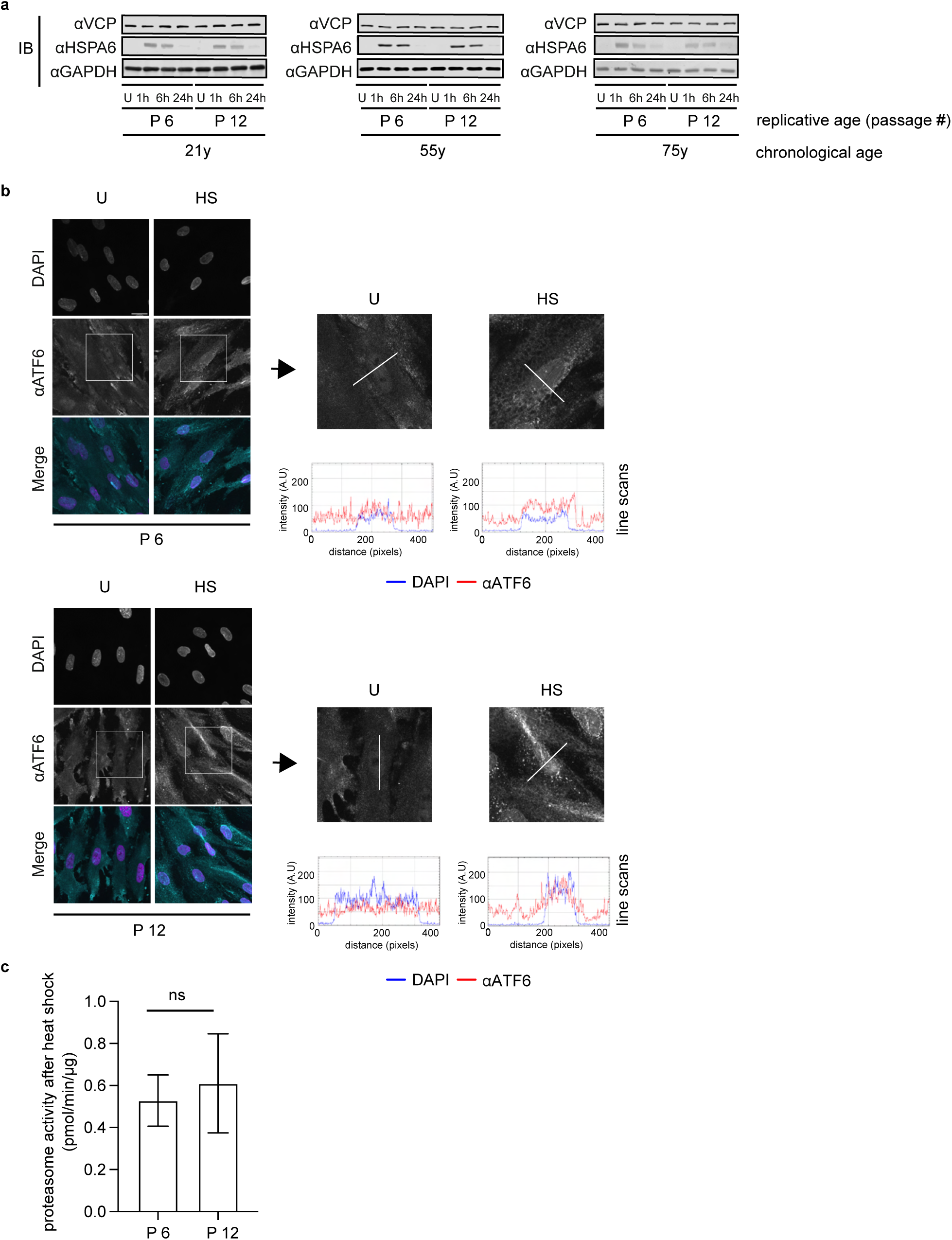
Hsp70-DNAJA1-DNAJB1 disaggregase activity is one of the earliest PQC functions to collapse during replicative aging of cells. **a**, Immunoblot analysis shows similar induction patterns of HSPA6 chaperone after heat shock (HS) in non-senescent P 6 and replicative aging P 12 human dermal fibroblasts (HDFs) derived from donors aged 21, 55 and 75 years indicating that HSR is fully intact in all cells. VCP levels also do not change in P 6 and P 12 HDFs. GAPDH, loading control, (n = 3). **b**, Nuclear shuttling of ATF-6 (cyan) required to activate the endoplasmic reticulum unfolded protein response (UPR_ER_) is equally active in P 6 and P 12 HDFs. Nuclei stained with DAPI (magenta). U and HS denote unstressed and heat shocked cells, respectively. HS performed at 44 °C for two hours in the incubator.^53^ Line scans across the nuclei are indicated in white. Corresponding line scan plots are shown on the right. Scale bar 20 µm (n = 3). Insets depict magnified cells. **c**, P 6 and P 12 HDFs show similar proteasome activity (see Methods) immediately after HS. ns – not significant, one-way ANOVA Tukey post hoc test.

**Supplementary Data Figure 23.**
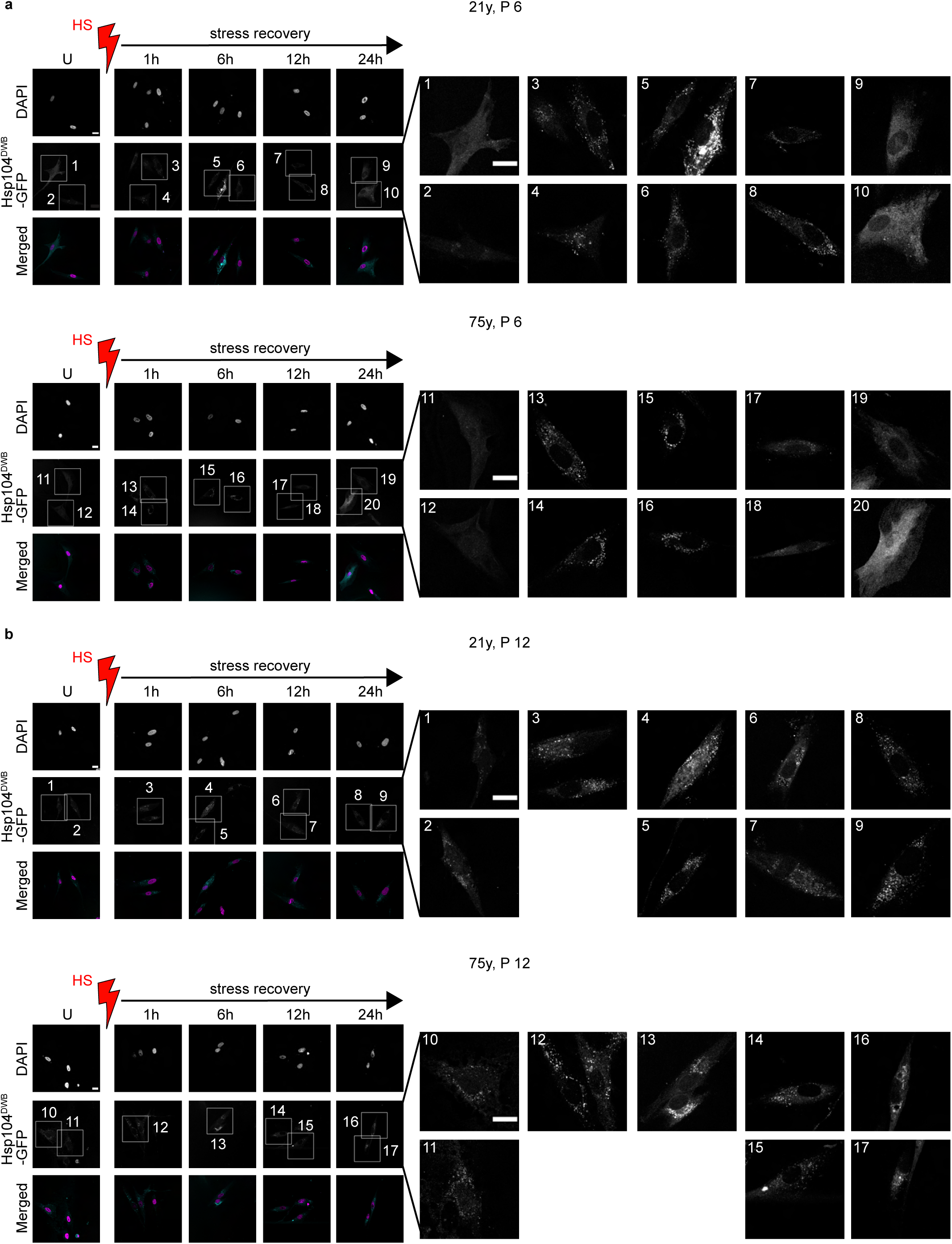
Persistence of heat-induced aggregates in aged human dermal fibroblasts. **a**, Regardless of chronological age, nonsenescent P 6 cells can efficiently solubilize heat-induced protein aggregates during recovery. Levels of Hsp104^DWB^-GFP positive aggregates (cyan) in unstressed and heat-shocked P 6 dermal fibroblasts derived from 21- and 75-year-old humans. Nuclei stained with Hoechst (magenta). White arrows show aggregates. U denotes unstressed cells. Heat shock (HS) in red. Scale bar 20 µm (n = 3). Insets depict magnified cells. **b**, Human dermal fibroblasts undergoing replicative aging (P 12) show defects in solubilizing heat-induced protein aggregates during recovery. Note that even unstressed cells (U) show Hsp104^DWB^-GFP positive aggregates. As in (**a**), Hsp104^DWB^-GFP positive aggregates (cyan) in unstressed and heat-shocked P 12 human dermal fibroblasts (21y and 75y). Nuclei stained with Hoechst (magenta). White arrows show aggregates. Scale bar 20 µm (n = 3). Insets depict magnified cells. P denotes the cell passage.

**Supplementary Data Figure 24.**
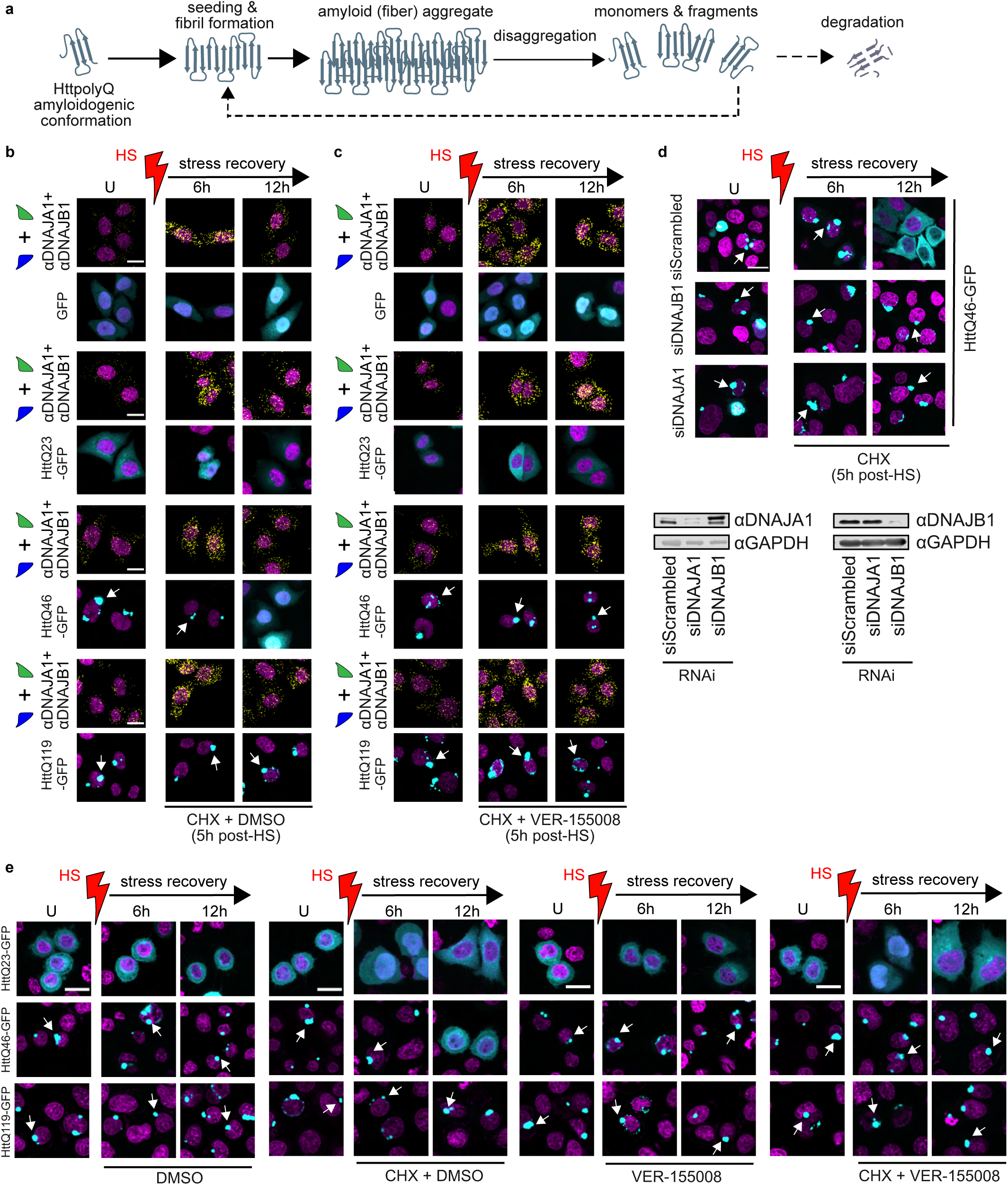
Hsp70-DNAJA1-DNAJB1 disaggregase activity disassembles disease-linked polyQ aggregates in cells. **a**, Disassembly of polyQ amyloids by disaggregation. **b**, Amyloid aggregates formed by HttQ46-GFP can be solubilized by inducing Hsp70-DNAJA1-DNAJB1 disaggregase activity. Top: The PLA signal (yellow) shows the induction of Hsp70-DNAJA1-DNAJB1 disaggregase in HeLa cells after heat shock (HS, red). Bottom: Solubilization of amyloids formed by HttQ-GFP variants with different lengths of glutamine stretches (HttQ23-GFP, HttQ46-GFP and HttQ119-GFP) after activation of Hsp70 disaggregase. HttQ23-GFP, a control that does not aggregate. Punctated GFP fluorescence (cyan) indicates aggregated polyQ (white arrows). Non-aggregating or solubilized polyQ generates a diffused fluorescence signal. Nuclei stained with DAPI (magenta). **c**, As in (**b**), inhibition of heat-induced Hsp70 disaggregase activity 5h after HS by adding VER-155008 blocks HttQ46-GFP amyloid disassembly. DMSO, vehicle control for the inhibitor. Cells were treated with cycloheximide (CHX) at 5h after HS to block the translation of new HttpolyQ proteins. U denotes unstressed cells. Scale bar 20 µm (n = 3). **d**, Depletion of DNAJA1 or DNAJB1 inhibits Hsp70-mediated polyQ disassembly. Top: The disassembly of HttQ46-GFP aggregates is inhibited in DNAJA1 and DNAJB1 depleted HeLa cells. Punctated GFP fluorescence (cyan) indicates aggregated HttQ46-GFP (white arrows). The solubilized HttQ46-GFP generates a diffused fluorescence signal. The non-targeting control siRNA (siScrambled) is used as the control for RNAi. Nuclei stained with DAPI (magenta). Hsp70-DNAJA1-DNAJB1 disaggregase is activated by HS. CHX was added 5 h after HS to inhibit the synthesis of new HttQ46-GFP molecules. Bottom: Immunoblot showing knockdown of DNAJA1 and DNAJB1 in HeLa cells expressing HttQ46-GFP. **e**, Disassembly of HttQ inclusions by heat-induced Hsp70 disaggregase is masked by high levels of aggregation of newly synthesized amyloidogenic proteins. Cells treated with +/− CHX with and without VER-155008. CHX was added 5h after HS to inhibit HttQ-GFP synthesis. VER-155008 was added 5h after HS to inhibit heat-induced Hsp70 disaggregase activity. DMSO, vehicle control for the inhibitor. GFP fluorescence (cyan) indicates the aggregated (punctated signal, indicated by white arrows) and non-aggregated (diffused signal) state of HttQ23-GFP, HttQ46-GFP, and HttQ119-GFP. HttQ23-GFP, HttQ control that does not aggregate. The nuclei stained with DAPI (magenta). Scale bar 20 µm (n = 3).

## Supplementary information 1

**Disassembly of disease-linked polyQ aggregates by Hsp70-DNAJA1-DNAJB1 disaggregase.** For this analysis, we used GFP tagged Huntingtin protein (Htt) exon 1 containing an expanded polyglutamine sequence (polyQ) (Supplementary Data Fig. 24a). These cytotoxic aggregates could form independently of proteotoxic stresses, such as heat. PolyQ aggregation was induced in unstressed HeLa cells (cyan foci, Supplementary Data Fig. 24b) by overexpressing Htt exon-1 carrying varying lengths of sequential glutamine stretches (HttQ46-mEmGFP and HttQ119-GFP; HttQ23-mEmGFP, non-aggregating control). The Hsp70-DNAJA1-DNAJB1 disaggregase was activated by HS and cycloheximide (CHX) was added 5h after HS to prevent aggregation of newly synthesized HttpolyQ molecules. CHX addition at 5h after HS had minimal effects on the induction of Hsp70-DNAJA1-DNAJB1 disaggregase. We observed the solubilization of pre-existing HttQ46-mEmGFP aggregates (disappearance of fluorescent puncta and appearance of a diffuse signal 12h after HS) with induction of the disaggregase (Supplementary Data Fig. 24b). Adding the Hsp70 inhibitor with CHX 5h after HS (Supplementary Data Fig. 24c) or knocking down DNAJA1 and DNAJB1 using RNAi (Supplementary Data Fig. 24d), completely abrogated amyloid solubilization. Together, these results indicate that Hsp70-DNAJA1-DNAJB1 disaggregase also has the ability to target amyloid aggregates in cells. Interestingly, the omission of CHX treatment from the experimental regime prevented the previously observed decrease in the steady-state level of HttQ46-mEmGFP aggregates (Supplementary Data Fig. 24e) despite the induction of Hsp70 disaggregase. Given the highly aggregate-prone nature of HttpolyQ species, we presume that the aggregation rate of newly synthesized HttQ46-mEmGFP monomers was faster than the disaggregation rate of the Hsp70-DNAJA1-DNAJB1 chaperone machine. In contrast, the amyloids formed by Htt exon-1 with a longer Q stretch (HttQ119-GFP) were completely resistant to the action of Hsp70-DNAJA1-DNAJB1 disaggregase even after blocking the aggregation of newly synthesized proteins using CHX (Supplementary Data Fig. 24b, e). These observations highlight important functional boundaries associated with the inherent capacity of this disaggregase to modulate amyloid load in cells.

## Notes

### Competing Interest Statement

The authors have declared no competing interest.

### Summary of Updates

Revised manuscript contains new data. Key additions include data pertaining to: 1. Assembly of Hsp70-DNAJA1-DNAJB1 disaggregase on heat-induced aggregates 2. Additional controls to support that the activity of Hsp70-DNAJA1-DNAJB1 disaggregase is impaired during early stages of replicative aging

